# Mass spectrometry and split luciferase complementation assays reveal the MecA protein interactome of *Streptococcus mutans*

**DOI:** 10.1101/2023.09.08.556943

**Authors:** Hua Qin, David Anderson, Zhengzhong Zou, Dustin Higashi, Christina Borland, Jens Kreth, Justin Merritt

## Abstract

MecA is a highly conserved adaptor protein encoded by prokaryotes from the *Bacillota* phylum. MecA mutants exhibit similar pleiotropic defects in a variety of organisms, although most of these phenotypes currently lack a mechanistic basis. MecA mediates ClpCP-dependent proteolysis of its substrates, but only several such substrates have been reported in the literature and there are suggestions that proteolysis-independent regulatory mechanisms may also exist. Here, we provide the first comprehensive characterization of the MecA interactome and further assess its regulatory role in Clp-dependent proteolysis. Untargeted coimmunoprecipitation assays coupled with mass spectrometry revealed that the MecA ortholog from the oral pathobiont *Streptococcus mutans* likely serves as a major protein interaction network hub by potentially complexing with >100 distinct protein substrates, most of which function in highly conserved metabolic pathways. The interactome results were independently verified using a newly developed prokaryotic split luciferase complementation assay (SLCA) to detect MecA protein-protein interactions *in vivo*. In addition, we further develop a new application of SLCA to support *in vivo* measurements of MecA relative protein binding affinities. SLCA results were independently verified using targeted coimmunoprecipitation assays, suggesting the general utility of this approach for prokaryotic protein-protein interaction studies. Our results indicate that MecA indeed regulates its interactome through both Clp-dependent proteolysis as well as through an as yet undefined proteolysis-independent mechanism that may affect more than half of its protein interactome. This suggests a significant aspect of MecA regulatory function still has yet to be discovered.

## Introduction

In prokaryotes, molecular chaperones and proteases play crucial roles in global protein quality control and the regulation of genetic networks. For this reason, such proteins exhibit extremely diverse protein interactomes, the vast majority of which remain poorly characterized in most bacteria. In the oral pathobiont *Streptococcus mutans*, these functions are largely performed by the Clp protease system, which consists of a ClpP protease subunit paired with any of several associated Clp ATPases: ClpC, ClpE, and ClpX (1–5). Of these three Clp ATPases, ClpC is unique in its requirement for an additional adaptor protein called MecA to stimulate proteolysis by ClpCP complexes (6,7). MecA is highly conserved among low G+C Gram-positive bacteria, but thus far, only a few MecA substrates have been reported within the entire *Bacillota* phylum (6,8–16). In *S. mutans*, MecA plays a critical role in the ClpCP-dependent degradation of the natural competence-specific alternative sigma factor ComX by mediating ternary complex formation with ComX and ClpC. Its N-terminal portion recognizes and binds to the target substrate ComX (residues 1-82), whereas the C-terminal portion complexes with ClpC (residues 123-240) (14). Depletion of ComX due to this interaction ultimately antagonizes natural competence development (14,17). An analogous MecA-dependent proteolytic mechanism also regulates natural competence development in *Bacillus subtilis* (9,11). In addition to natural competence, MecA likely participates in a variety of additional metabolic pathways as well. For example, MecA-overproduction in *B. subtilis* triggers sporulation and biofilm defects (12,16), while a *mecA* deletion reduces cell viability and triggers defective cell division (18). *S. mutans mecA* mutants exhibit similar pleiotropic defects in cell envelope biogenesis, cell division, and biofilm formation, suggesting that it too likely contains a highly diverse, yet largely uncharacterized interactome (19). It is worth noting that among the limited number of confirmed MecA substrates, not all are regulated through proteolysis. In *B. subtilis*, MecA-ClpC complexes have been suggested to bind and modulate the master regulator Spo0A∼P, subsequently preventing its transcription activation ability through a mechanism independent of both proteolysis and phosphorylation (12,16). Thus, there is reason to suspect that MecA regulates its interactome through both functional and proteolytic mechanisms.

Recent improvements in the technologies used to detect and characterize protein-protein interactions (PPI) have stimulated new interest to reveal protein interactomes, leading to the discovery of many unexpected PPI, especially those involved in previously unrecognized moonlighting functions (20–25). Several genetic and biochemical techniques are commonly employed to detect PPI, such as two-hybrid assays, coimmunoprecipitation (co-IP), bimolecular fluorescence complementation (BiFC), and Förster resonance energy transfer (FRET) (26–29). Another related technology, the split luciferase complementation assay (SLCA), has emerged as an increasingly popular genetic approach to study PPI, due to the simplicity of the assay as well as its extraordinary sensitivity. In principle, SLCA works much like the classic two-hybrid approach and BiFC. The luciferase enzyme is split and expressed as two separate chimeric protein fusions that are reconstituted into a functional reporter only when the two fused target proteins of interest form stable complexes. SLCA has been used to measure dynamic changes in PPI in numerous studies of viruses and mammalian cells, but thus far has only been rarely employed for prokaryotic genetic research (30–39).

In recent years, our group has regularly utilized coimmunoprecipitation (co-IP) to assay PPI in *S. mutans* (40–43). While we have found this biochemical approach to be highly reliable and robust, co-IP is also time-consuming, labor-intensive, and can be quite costly when using antibody affinity resins. Thus, we have been interested in augmenting these biochemical studies with a simpler alternative genetic approach. We have previously performed BiFC in *S. mutans* (41), which we found to be suitable for imaging purposes, but this approach did not exhibit the sensitivity and dynamic range necessary for robust PPI characterization. The two-hybrid approach has been employed to study *S. mutans* PPI (17,44,45), but thus far, this approach has required protein expression in ectopic hosts. Our preference has been to study PPI in their native contexts, which precludes the current two-hybrid reporter systems. For this reason, it was of interest to explore the potential utility of SLCA in *S. mutans*. We previously developed a suite of synthetic codon-optimized luciferases for use in streptococci and were able to demonstrate their utility for both typical transcriptional/translational fusion reporter assays as well as for *in vivo* biophotonic imaging applications (46–50). In our experience, luciferases vastly outperform all other reporter systems for their simplicity, precision, and dynamic range, especially when using derivatives of the Renilla luciferase enzyme from *Renilla reniformis*. Furthermore, the GFP-scanning approach has been previously applied in viruses and mammalian cells to identify SLCA split sites in Renilla luciferase that yield strong complementation activity with minimal self-assembly (51–53). Therefore, we were curious whether these same split sites could be adaptable to implement SLCA in *S. mutans*, particularly for the study of proteins with complex interaction networks. Accordingly, the *S. mutans* MecA ortholog was selected as an ideal target protein for the development of a prokaryotic SLCA due to MecA’s role as a conserved pleiotropic regulator with a largely uncharacterized protein interactome.

## Results

### MecA interactome analysis

To identify the MecA protein interactome, we appended a tandem affinity purification (TAP) epitope tag (3x HA + FLAG) onto the C-terminus of MecA. Cultures were grown to mid-logarithmic phase, washed, and then formaldehyde crosslinked before performing antibody affinity chromatography. Protein lysates were first immunopurified using anti-FLAG affinity resin, and then the resulting protein eluates were further purified with anti-HA affinity resin. The final immunopurified protein complexes were analyzed by liquid chromatography with tandem mass spectrometry. Specific enrichment was calculated by comparing the mass spectrometry spectral count values for individual TAP-tagged proteins vs. the wild-type negative control (i.e., no TAP tag). We performed this interactome analysis three times, and identified >100 candidate MecA-interacting proteins exhibiting ≥ 2-fold enrichment (Table 1). Consistent with previous studies in *S. mutans* and/or other bacteria, MecA was found to form heteromeric complexes with the protease ClpP as well as the Clp ATPases ClpC, ClpX, and ClpE (4,14,15,17,54). However, most of the identified MecA interaction partners have not been previously reported. These proteins function in a diverse assortment of highly conserved metabolic pathways, like amino acid biosynthesis (GltB, GltA, Akh, IIvD), DNA metabolism and transcription (UvrA, XseA, GyrB, GyrA, TopA, NrdE), cell division (FtsQ, FtsZ, SepF), cell envelope biogenesis (MurM, GPI, MurA2, MurD), and biofilm formation (GtfB, GtfC, SpaP). Many of the identified MecA protein interactions were detected independently in multiple datasets with strong enrichment, such as GltA and GltB (glutamate synthase subunits), ClpP and ClpC (Clp protease), TopA (type I DNA topoisomerase), SpaP (cell surface antigen I/II), and others (Table 1).

**Table 1.**
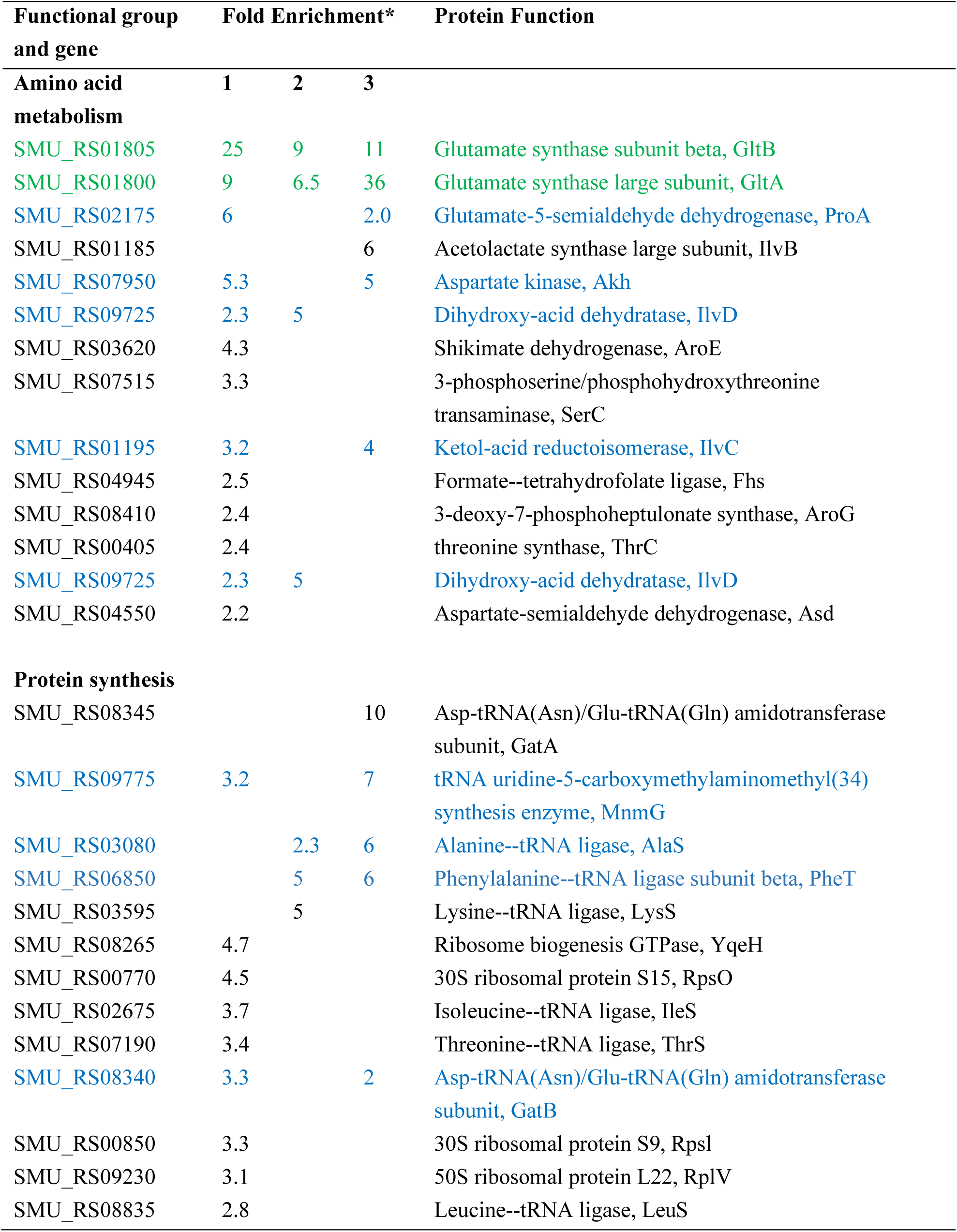
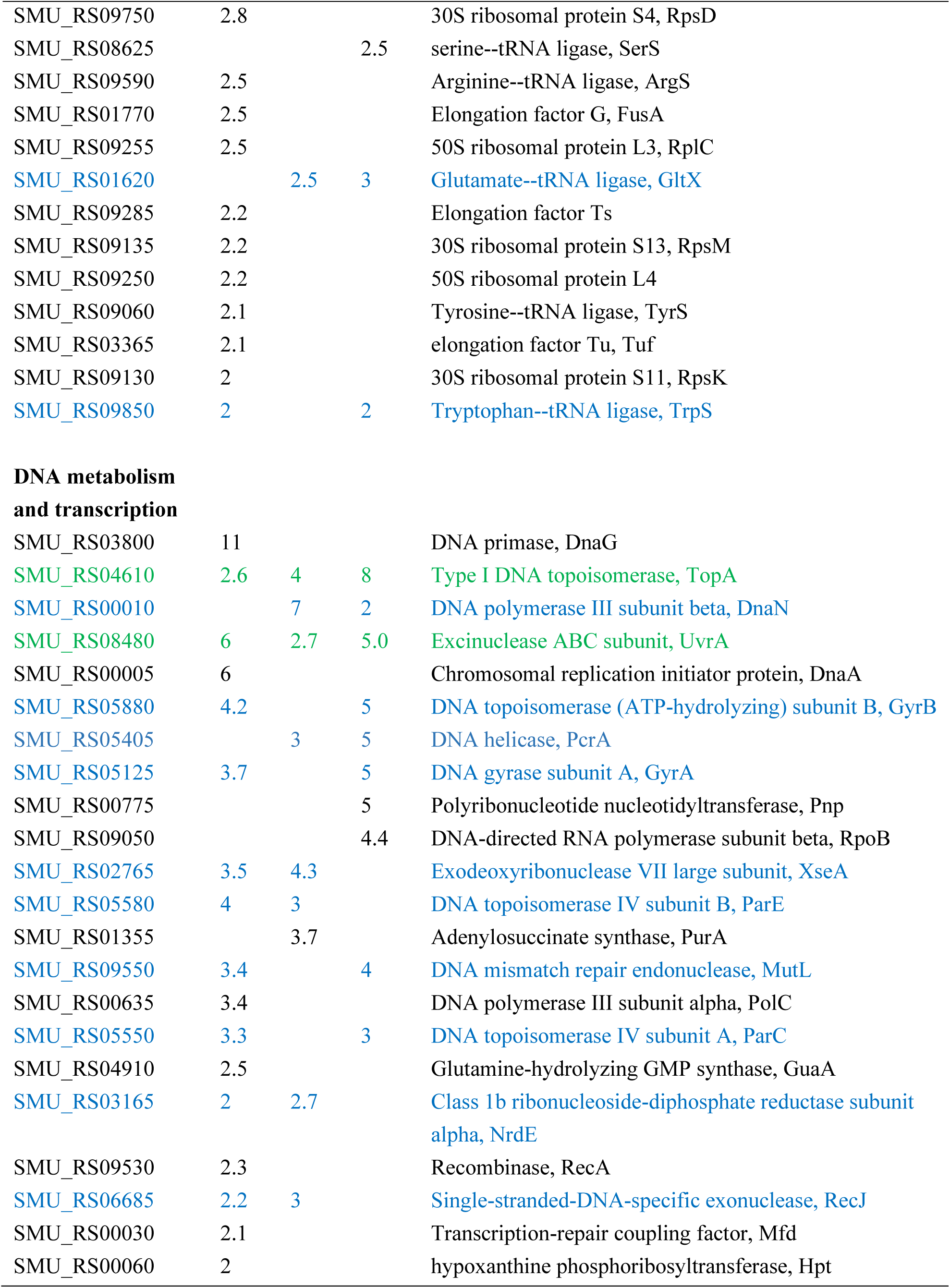
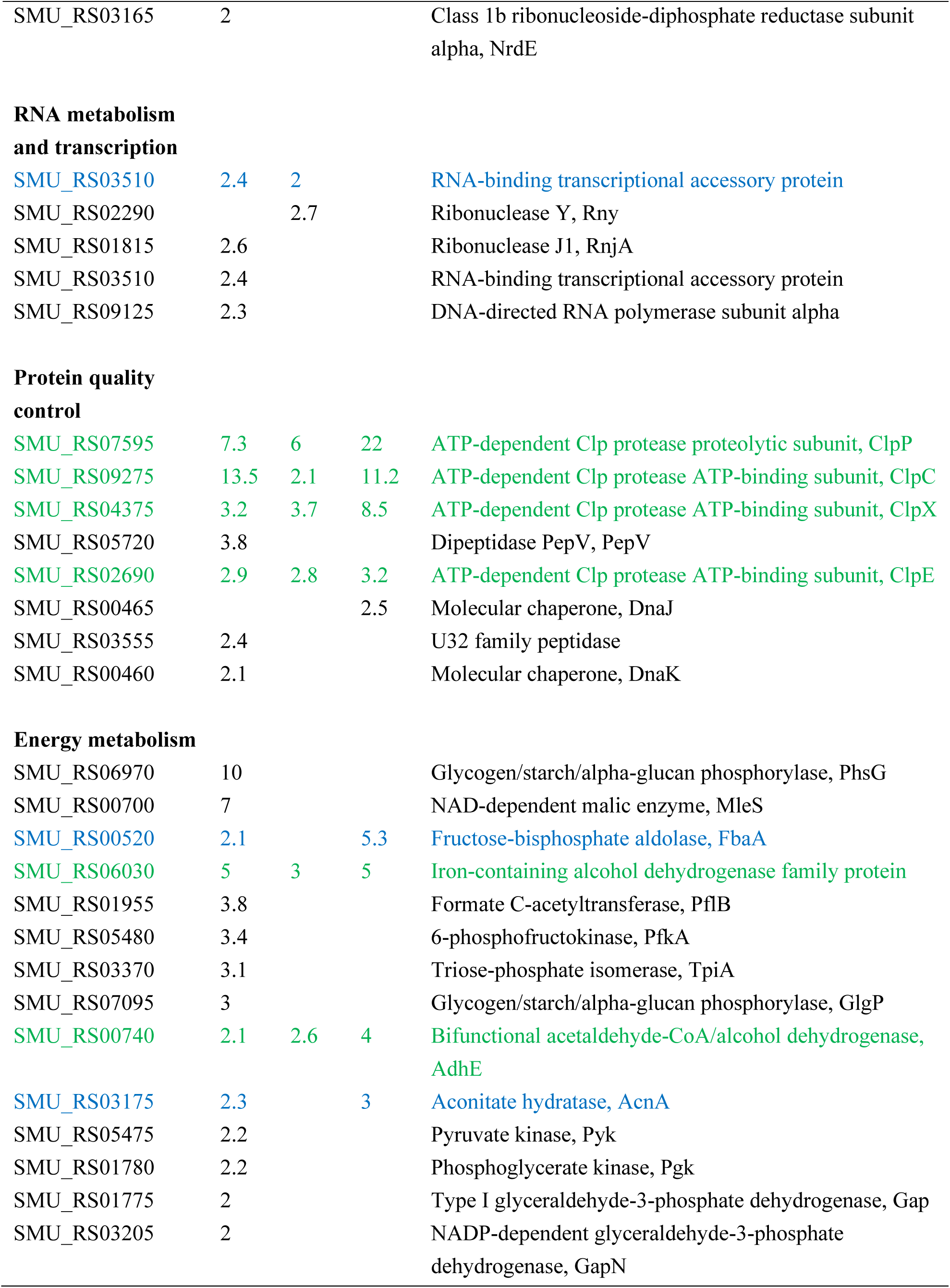
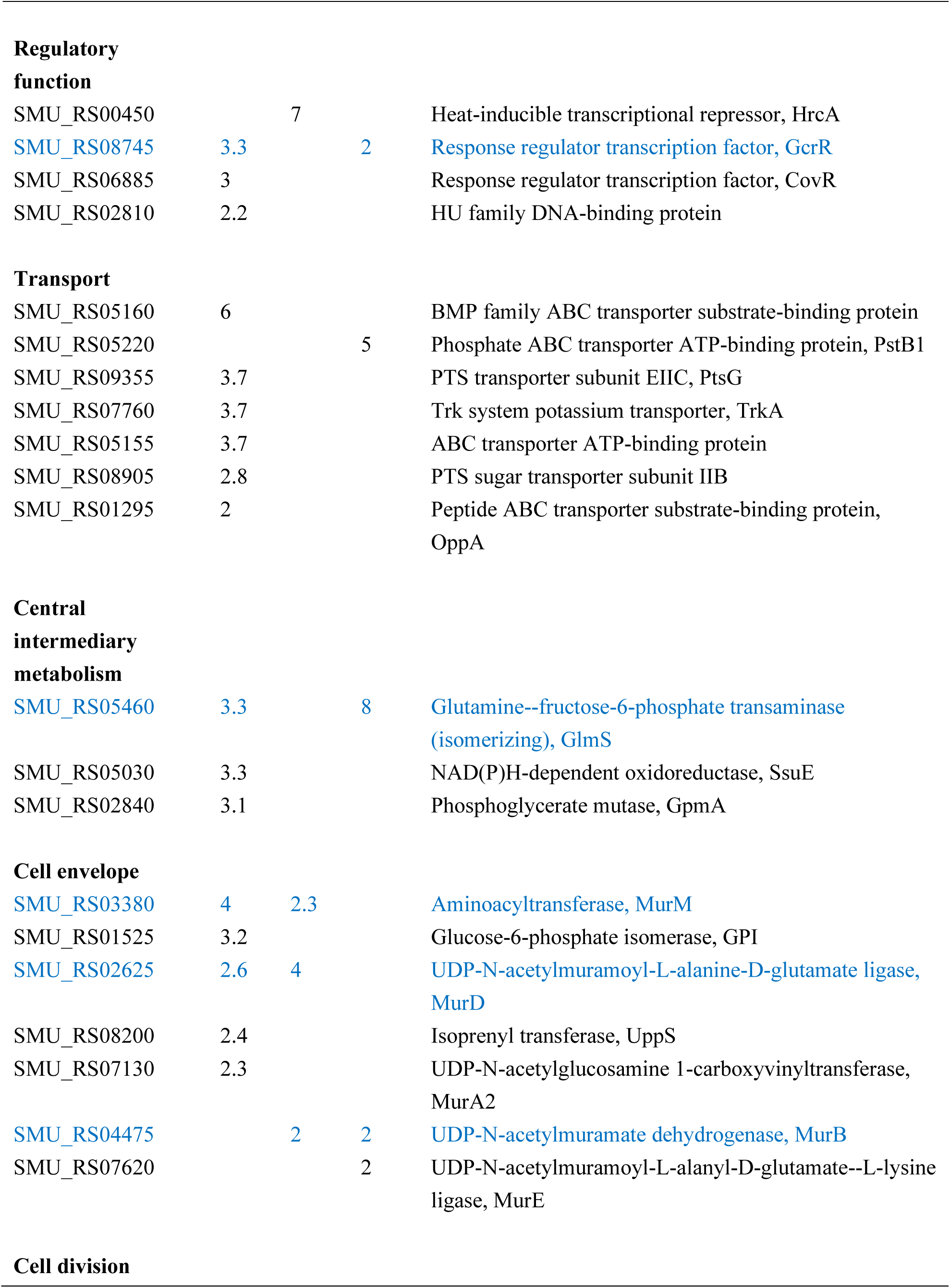
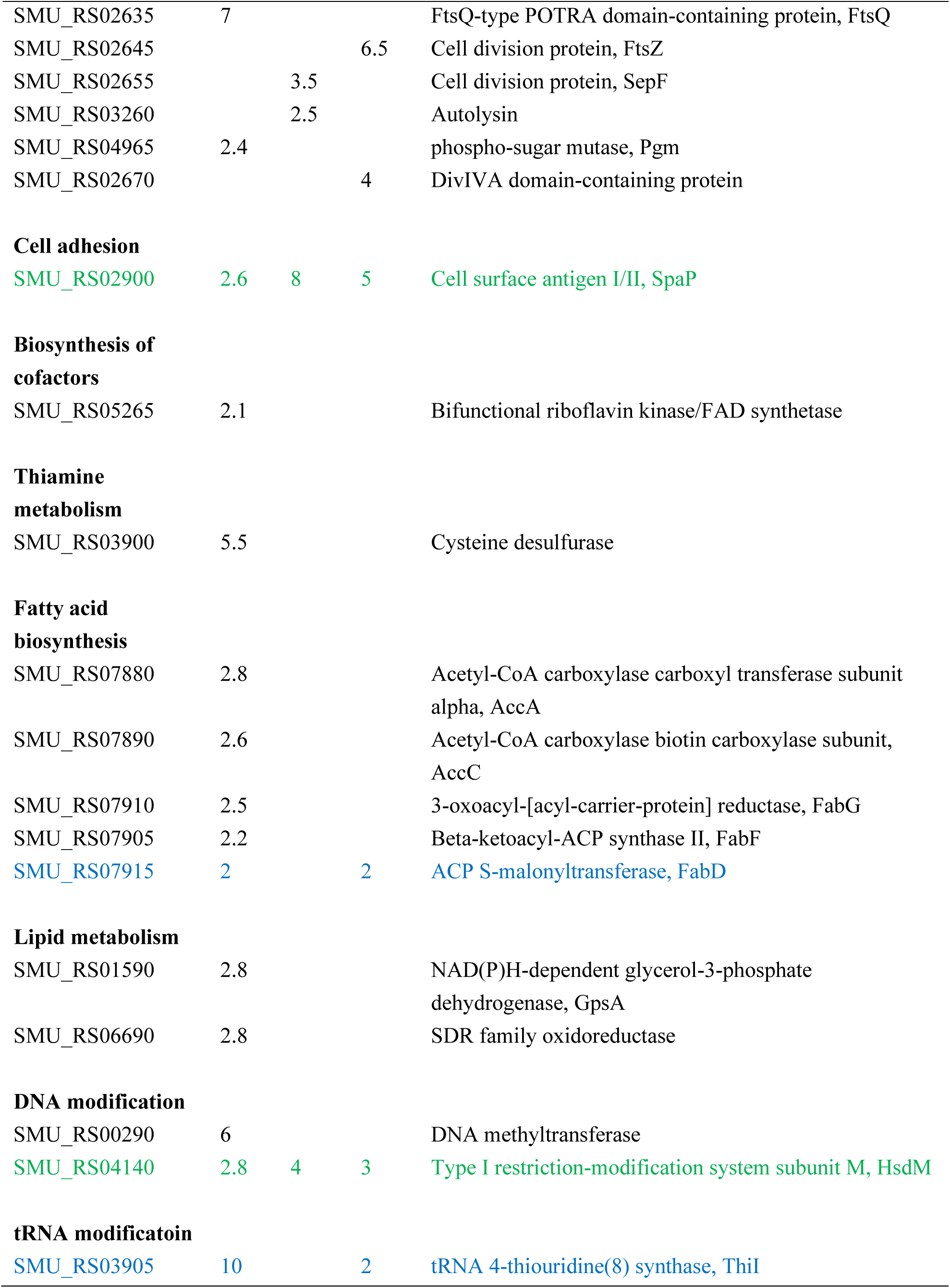
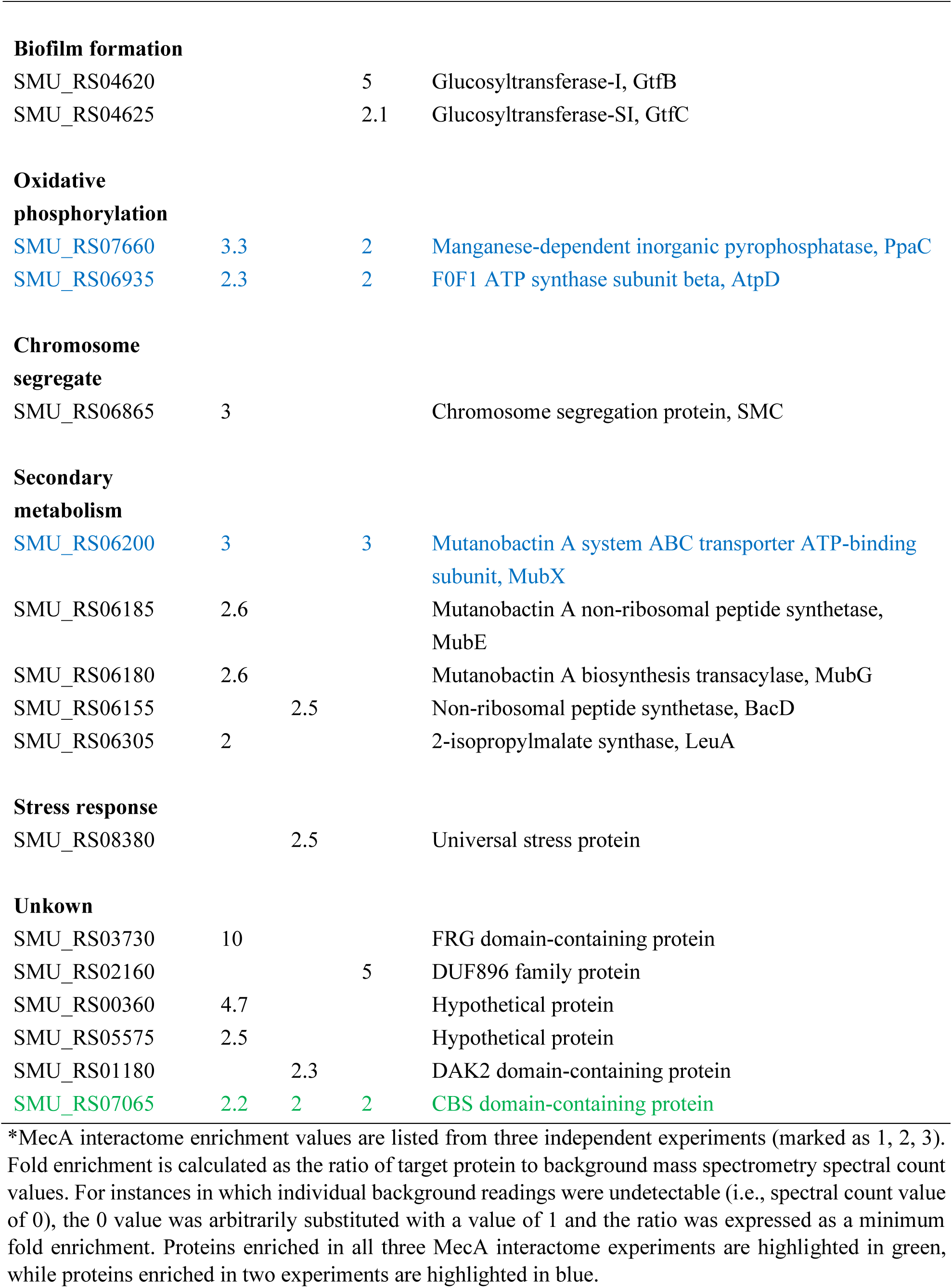
Proteins exhibiting ≥2-fold enrichment in the MecA interactome screen.

### Development of a Green Renilla (RenG) split luciferase complementation assay

Given the large diversity of potential MecA interactions detected via co-IP/mass spectrometry, we were interested to develop a split luciferase complementation assay (SLCA) as a genetic approach to independently confirm these PPI *in vivo*. Based upon a previous split luciferase assay using the Renilla luciferase variant RLuc8 (53), we compared two equivalent sets of split sites in a codon-optimized Green Renilla luciferase variant RenG located between residues 155-156 and 229-230 (46). Both split sites fall within predicted unstructured loop regions in the RenG structural model (Figure 1A and 1B). To facilitate the proper folding of protein chimeras, a flexible linker (either 2× GGGGS or 3× GGGGS) was inserted between the protein of interest and the RenG luciferase fragment (Figure 1C). To test both candidate RenG split sites, we first created C-terminal SLCA fusions to RNases J1 and J2 because these enzymes form extremely avid heteromeric complexes and both enzymes are highly tolerant of protein fusions (40,55,56). As shown in Figure 1D, both RenG SLCA split sites yielded robust signals when fused to RNases J1 and J2, with the 155-156 split site yielding about 25% greater signal intensity compared to the 229-230 split site (Figure 1D and Table S3). As expected, the unfused split RenG luciferase fragments yielded extremely low reporter activity that was <15% higher than the background luminescence values, indicating a complete lack of detectable self-assembly between the luciferase fragments (Figure 1D and Table S3). Given the slightly higher output from the 155-156 RenG split site, this was chosen for all subsequent split luciferase assays.

**Figure 1.**
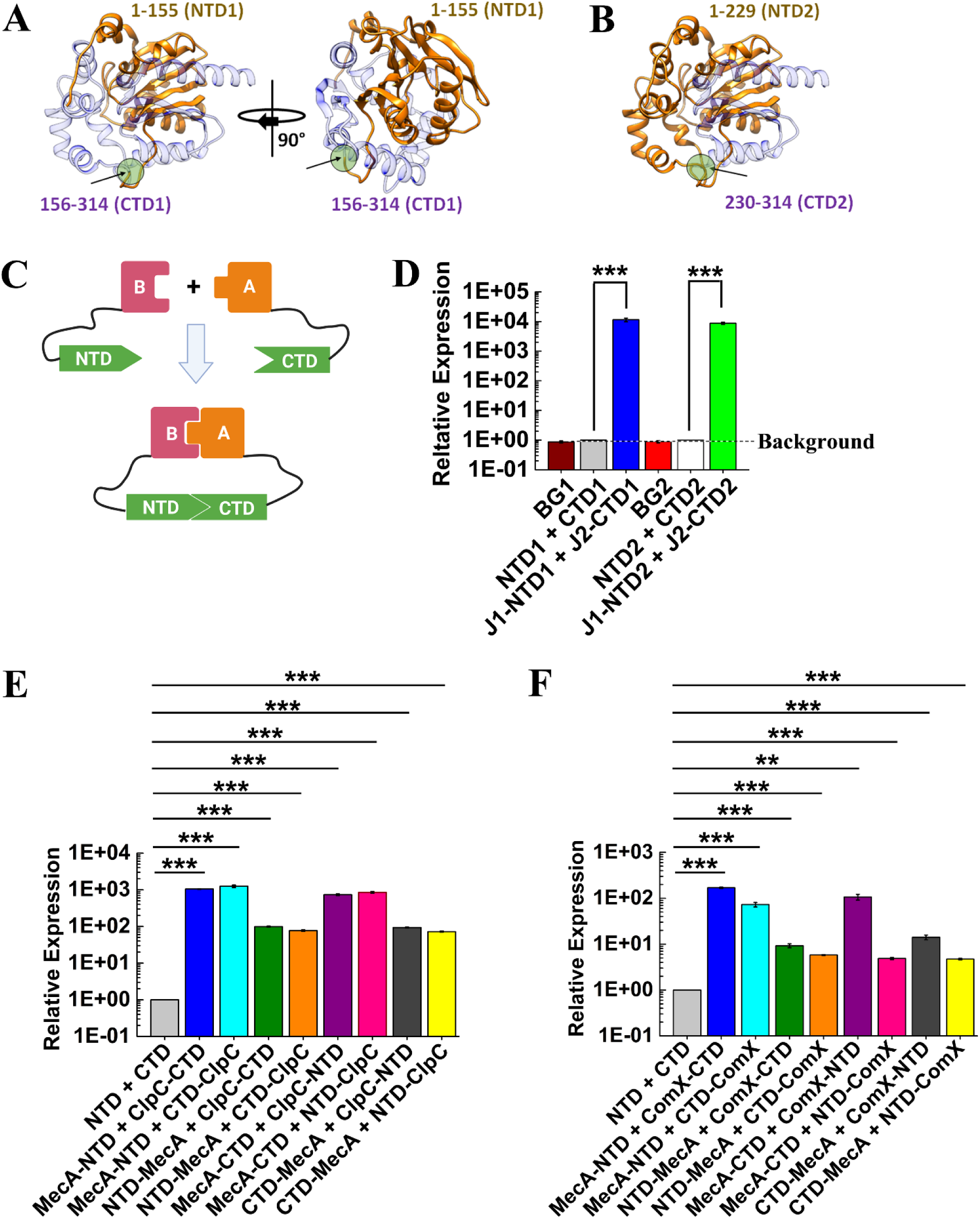
Development of a split luciferase complementation assay. A and B, Structural models of Green Renilla luciferase (RenG) split luciferase segments. The split site is highlighted by a green circle and an arrow. (A). The RenG 1-155 domain (NTD1, shown in orange) and the 156-314 domain (CTD1, shown in purple) interface includes 77 and 86 residues, respectively, encompassing 3403 Å^2^ total and an estimated solvation free energy gain (Δ^i^G) of -46.3 kcal/mol upon complex formation (PDBePISA, ‘Protein interfaces, surfaces and assemblies’ service PISA at the European Bioinformatics Institute. https://www.ebi.ac.uk/pdbe/prot_int/pistart.html. (82)). (B). The RenG 1-299 domain (NTD2, shown in orange) and 230-314 domain (CTD2, shown in purple) interface includes 72 and 58 residues, respectively, encompassing 2613 Å^2^ total and an estimated solvation free energy gain (Δ^i^G) of -39.4 kcal/mol upon complex formation. (C). Schematic representation of the split luciferase complementation assay (SLCA). The RenG N-terminal domain (NTD) is attached to protein B via the linker peptide 2x or 3x GGGGS (illustrated as a thin black line). The RenG C-terminal domain (CTD) is fused to protein A using the same linker. A stable heteromeric complex formed between proteins B and A supports reconstitution of RenG enzymatic activity, yielding a quantifiable reporter signal. (D). Different RenG split sites were tested to analyze RNase J1-RNaseJ2 heteromeric complex formation. The N-terminal domain of RenG (NTD1 or NTD2) was translationally fused to the C-terminus of RNase J1, while the C-terminal domain of RenG (CTD1 or CTD2) was translationally fused to the C-terminus of RNase J2. Luciferase activity was normalized to culture optical density (OD_600_) and then expressed relative to the free split luciferase reporter strain values, which were arbitrarily assigned values of 1. The luminescence values obtained from the wild-type strain (i.e., no luciferase fusion) were assigned as the assay background (BG1 and BG2). (E). The 1-155 domain (NTD) or 156-314 domain (CTD) of RenG were translationally fused to the N- or C-terminus of MecA or ClpC to analyze MecA-ClpC complexes. (F). The NTD or CTD of RenG was translationally fused to the N- or C-terminus of MecA or ComX to analyze MecA-ClpC complexes. All luciferase data are expressed as the means ± s.d. (indicated by error bars) derived from three or four biological replicates. \*\*\**p*<0.001, \*\**p*<0.01 Unpaired two-tailed Student’s *t-*test with Welch’s correction.

After demonstrating the utility of RenG SLCA for measuring RNase J1 and J2 complexes, we were next interested to optimize this assay to examine different components of the MecA interactome. Thus, we began by testing one of the few known MecA-dependent interactions in *S. mutans*, the MecA-ClpC-ComX ternary complex (14). Both MecA-ClpC and MecA-ComX complexes were configured in all eight possible N- and C-terminal chimera combinations to compare their signal outputs (Figure 1E and 1F). Although all of the chimeras yielded robust reporter signals substantially above background values, we observed the strongest signal outputs using RenG SLCA fusions to the C-terminus of MecA combined with SLCA fusions to the N-terminus of ClpC, whereas the MecA-ComX SLCA results were strongest with C-terminal fusions to both proteins (Figure 1E and 1F). The N-terminal domain (NTD) of RenG was the optimal luciferase fragment fusion for MecA in both MecA-ClpC and MecA-ComX SLCA.

### Validation of MecA interactome results using RenG split luciferase complementation assays

Given the promising results obtained with RenG SLCA of RNase J1-J2 and MecA-ClpC-ComX complexes, we next selected a diverse subset of 13 proteins identified in the MecA interactome data (Table 1) and assayed their potential interactions with MecA using the RenG SLCA. The selected proteins represented a wide range of mass spectrometry enrichment values to examine SLCA with both high and low avidity complexes. We first examined fusions of the RenG NTD (1–155) to the N- or C-terminus of MecA paired with RenG C-terminal domain (CTD; 156-314) fusions to the N- or C-terminus of GltB and ThiI. Both MecA-GltB and MecA-ThiI pairs yielded strong SLCA signals, indicating their stable interaction *in vivo* (Figure 2A and 2C). Consistent with previous observations, luciferase signals were noticeably stronger with RenG NTD fusions to the C-terminus of MecA (Figure 2A and 2C). Therefore, for the remainder of the tested SCLA fusions, we only examined MecA reporters with C-terminal fusions to the RenG NTD. In agreement with the interactome data, robust SLCA signals were detected from all of the tested proteins, except for sortase (SrtA), which was not identified in the interactome data (Table 1) and was included as a negative control (Figure 2A – 2N). For most of the assayed MecA interaction partners, C-terminal fusions with the RenG CTD generated the strongest SLCA signals, with DNA primase (DnaG) being the lone exception. Unexpectedly, SLCA signals from the MecA-SpaP interaction were noticeably weaker than the other positive interactions (Figure 2M).

**Figure 2.**
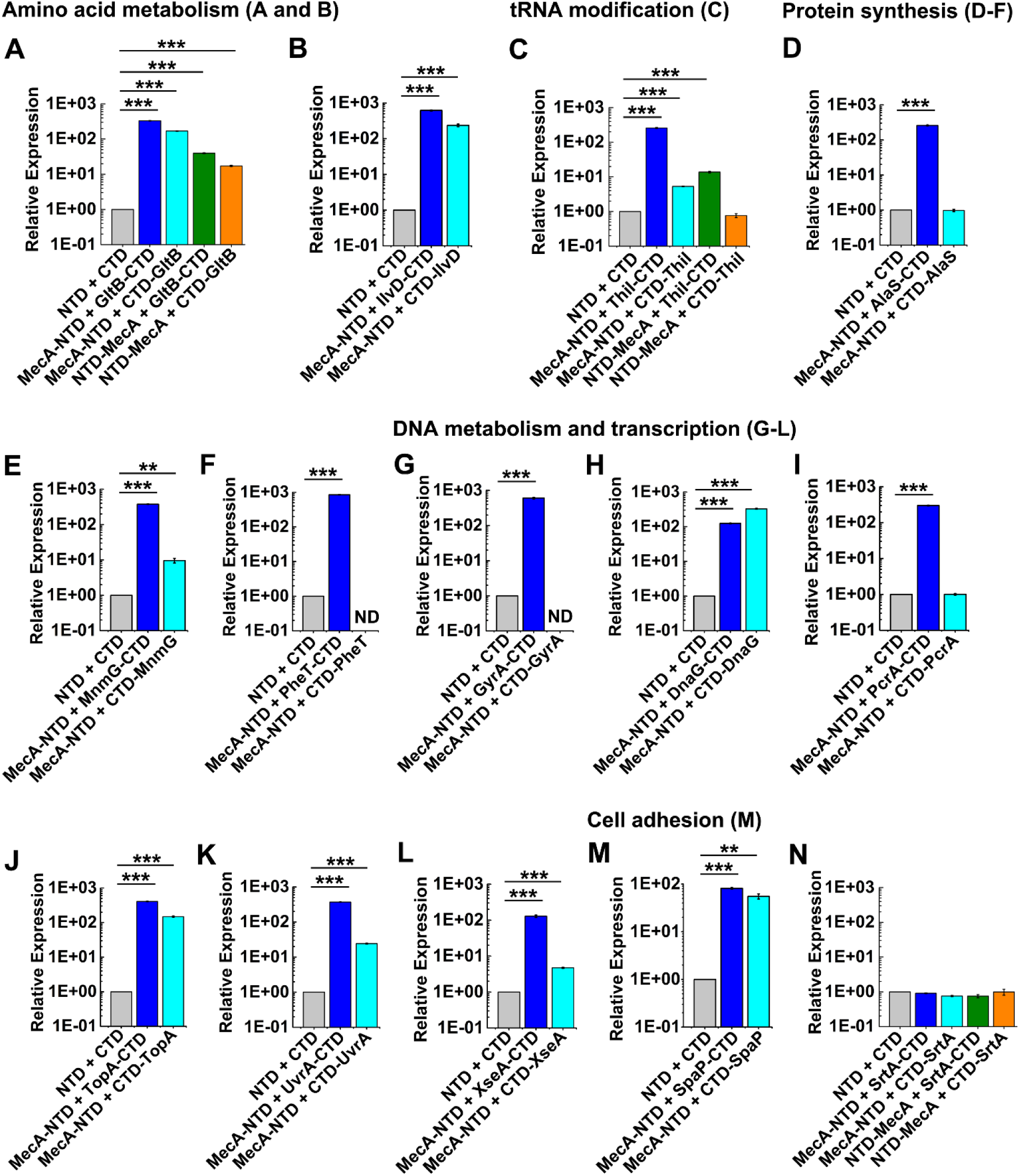
Split luciferase complementation assays of selected MecA interactome proteins. The RenG 1-155 amino acid NTD was translationally fused to the N- or C-terminus of MecA, while the RenG 156-314 CTD was translationally fused to the N- or C-terminus of candidate MecA interacting proteins. Cultures were grown to mid-logarithmic growth phase, normalized to the measured optical density (OD_600_) values, and then expressed relative to the luciferase values of the unfused NTD and CTD split luciferase fragments, which were arbitrarily assigned values of 1. Normalized luciferase values were measured to analyze MecA interactions with (A) GltB, (B) IlvD, (C) ThiI, (D) AlaS, (E) MnmG, (F) PheT, (G) GyrA, (H) DnaG, (I) PcrA, (J) TopA, (K) UvrA, (L) XseA, (M) SpaP, and (N) SrtA. ND (not determined due to apparent lethality). All luciferase data are expressed as means ±s.d. (indicated by error bars) derived from three or four biological replicates. \*\*\**p*<0.001, \*\**p*<0.01 Unpaired two-tailed Student’s *t-*test with Welch’s correction.

### Assessing protein binding affinities using the RenG split luciferase complementation assay

Based upon the robust luciferase signals obtained with split RenG, we were curious whether this assay may also be useful as a simple *in vivo* qualitative assessment of protein interaction avidity. We reasoned that highly avid protein complexes would yield SLCA values much closer to the theoretical maximum luciferase activity compared to low avidity complexes, which exhibit more transient interactions that readily dissociate. We considered the theoretical maximum potential SLCA value to be the reporter signal obtained from a full-length luciferase chimera to the protein of interest. In this case, relative binding affinity could be expressed as a simple ratio between the measured SLCA values versus a chimera with the full-length RenG (i.e., theoretical maximum) (Figure 3A). To determine the theoretical maximum SLCA value, both proteins of interest are separately fused to an intact full-length RenG, and the protein chimera yielding the weaker reporter activity (i.e., lower abundance protein) is set as the maximal luciferase value obtainable via SLCA of the two proteins. As shown in Figure 3B, we created whole RenG chimeras to each of the proteins previously tested via SLCA and observed a wide range of luciferase values that varied by >30-fold. We next divided all of the measured SLCA values by their corresponding maximal potential luciferase values to obtain relative protein binding affinities (Table 2). Consistent with both previous studies and predicted protein complex interface values, SLCA measurements of the RNase J1-J2 complex yielded exceptionally strong relative binding affinities, achieving nearly 98% of the theoretical maximum (Table 2 and Table S3) (40,55). Given the extensive interaction interface between RNase J1 and J2 (Figure S1), these complexes would be predicted to rarely dissociate, which agrees strongly with our previous findings (40) as well as the SLCA results (Table 2). Since the actual binding affinity (K_d_) of the *S. mutans* RNase J1-J2 complex has never been measured, we also assayed the relative binding affinity of ectopically expressed luciferase fusions to the *Bacillus amyloliquefaciens* ribonuclease barnase and its specific inhibitor barstar as a control reaction. The exceptionally high avidity of the barnase-barstar complex has been studied in detail (57–59), and as expected, SLCA measurements similarly indicated a very strong relative binding affinity, achieving more than 53% of the theoretical maximum (Figure S2, Table 2 and Table S3). Based upon a protein interface analysis of structural models for the *S. mutans* MecA-ClpC complex, we predicted that it too would form a highly avid complex, but less so compared to RNase J1-J2 (Figure S1). Accordingly, MecA-ClpC SLCA yielded a 55% relative binding affinity (Table 2 and Table S3), which was quite similar to the barnase-barstar results. While the RNase J1-J2, barnase-barstar, and MecA-ClpC relative binding affinities were all consistent with expectations, only weak correlations were noted between the interactome enrichment values measured via mass spectrometry and the relative binding affinities measured by SLCA (Table 1 and 2). For example, the enrichment of GltB was consistently among the strongest in the mass spectrometry data (Table 1), but the relative binding affinity measured for GltB-MecA complexes suggested only a moderate interaction affinity (Table 2). Mass spectrometry spectral count values are known to be impacted by a variety of variables, like protein abundance, length, and amino acid composition, all of which can subsequently influence the final enrichment values in the interactome dataset. Therefore, to further assess the validity of the relative binding affinities measured via SLCA, we employed a third approach using targeted co-IP assays. Candidate MecA-interacting proteins were translationally fused with a 3× HA epitope tag, whereas a 3× FLAG epitope tagged MecA served as the bait protein. Sortase (SrtA) was again employed as a negative control. MecA protein complexes were immunopurified with anti-FLAG affinity resin, and then the co-purified HA-tagged candidate proteins were detected via anti-HA western blots. When comparing the abundances of MecA interactome proteins found within the input protein lysates vs. the resulting output co-IP samples, it was obvious that the interactome proteins displayed a wide range of MecA binding affinities. ClpC exhibited the highest MecA binding affinity, while GyrA, ThiI, and GltB exhibited moderate binding affinities, AlaS showed weak MecA binding, and as expected, we did not detect any MecA interaction in the SrtA sample (Figure 4). After comparing the SLCA relative binding affinity assays results (Table 2) with the targeted co-IP results (Figure 4), we noticed a clear correlation among all of the assayed interactions, in contrast to the fairly weak correlations observed with the co-IP/mass spectrometry enrichment values.

**Figure 3.**
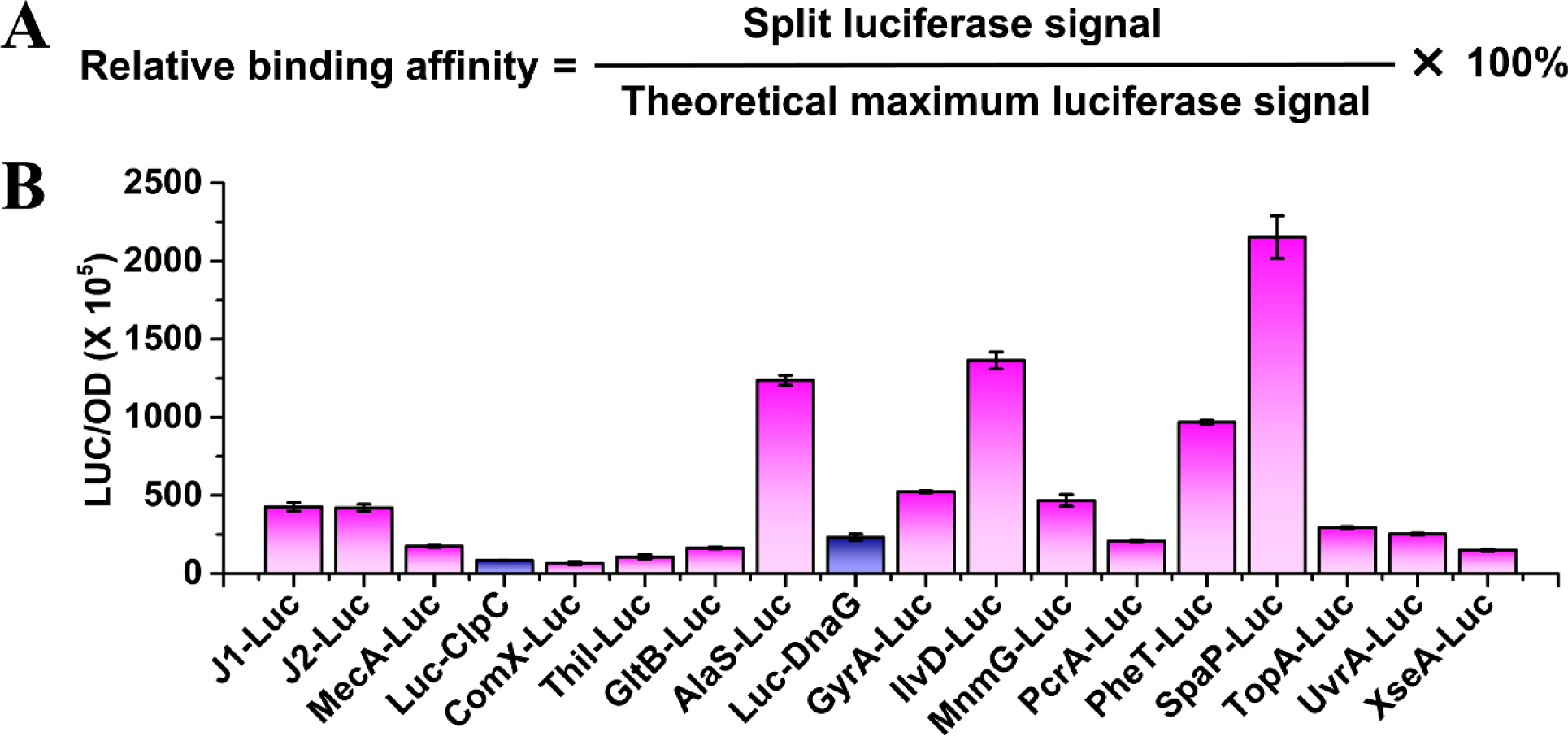
Determination of maximal luciferase values for selected MecA interactome proteins. (A) Relative protein binding affinity is determined by calculating the ratio of measured SLCA values compared to the theoretical maximum luciferase value for the protein complex of interest. (B) Cultures were grown to mid-logarithmic growth phase, and luciferase activity was normalized to the measured optical density (OD_600_) values. All luciferase data are expressed as means ±s.d. (indicated by error bars) derived from three or four biological replicates. Pink bars indicate C-terminal luciferase fusions, while purple bars indicate N-terminal fusions.

**Figure 4.**
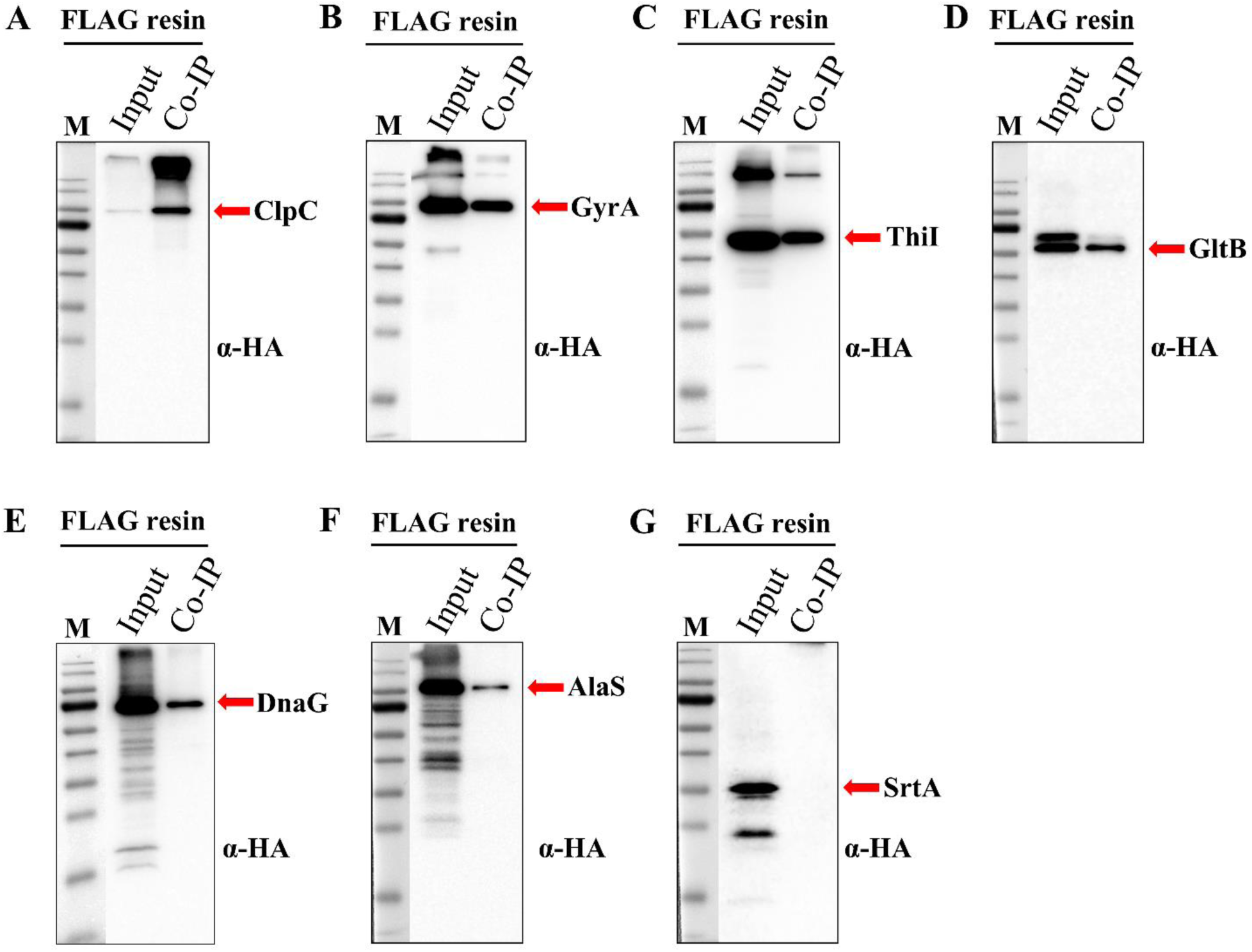
Coimmunoprecipitation of selected MecA interactome proteins. 3×FLAG tagged MecA was used as a bait to coimmunoprecipitate HA tagged MecA interactome proteins having predicted high, medium, and low affinities as reported in Table 2. Total protein was extracted from cultures grown to mid-logarithmic phase. Anti-FLAG affinity resin was used for immunoprecipitation, while anti-HA antibodies (α-HA) were used for western blot detection of coimmunoprecipitated proteins. The pre-stained protein ladder molecular weights from top to bottom: 170, 130, 95, 72 (strong reference band), 56, 43, 34, 26, 17, and 11 KDa. MecA coimmunoprecipitated proteins: (A) ClpC, (B) GyrA, (C) ThiI, (D) GltB, (E) DnaG, (F) AlaS, and (G) SrtA.

**Table 2.** MecA interactome protein relative binding affinity.

| Functional group | MecA interactome | Relative binding affinity (%) |
| --- | --- | --- |
| **RNA metabolism** | RNase J1-RNase J2 | 97.76 ± 8.06 |
| | Barstar-Barnase | 53.09 ± 1.01 |
| **Protein quality control** | MecA-N-ClpC | 55.22 ± 4.16 |
| **Protein synthesis** | MecA-PheT | 17.95 ± 0.77 |
| | MecA-MnmG | 8.00 ± 0.33 |
| | MecA-AlaS | 5.50 ± 0.29 |
| **Amino acid metabolism** | MecA-IlvD | 13.11 ± 0.61 |
| | MecA-GltB | 7.45 ± 0.28 |
| **DNA metabolism and transcription** | MecA-GyrA | 12.87 ± 0.78 |
| | MecA-TopA | 8.64 ± 0.36 |
| | MecA-UvrA | 7.86 ± 0.30 |
| | MecA-N-DnaG | $6.97 \pm 0.32$ |
| | MecA-PcrA | $6.36 \pm 0.28$ |
| | MecA-XseA | $3.16 \pm 0.25$ |
| **tRNA modification** | MecA-Thil | $8.92 \pm 1.07$ |
| **Quorum sensing** | MecA-ComX | $6.30 \pm 0.94$ |
| **Cell adhesion** | MecA-SpaP | $1.72 \pm 0.083$ |

### Assessing the role of MecA in Clp-dependent proteolysis

As previously mentioned, MecA is thought to primarily function as an adaptor protein required for stimulating ClpCP-dependent proteolysis of its interactome. While there is compelling evidence in support of this function (6–10,13–15,17,60), there are also indications that MecA might regulate some proteins at the functional level instead of via proteolysis (12,16). Given the limited number of known MecA substrates, it has been difficult to reconcile these previous observations. Therefore, it was of interest to examine the newly identified *S. mutans* MecA interactome to determine whether it is primarily regulated via Clp-dependent proteolysis. To test this, the abundances of a subset of the MecA interactome was measured in the *mecA* mutant and *mecA*/*clpP* double mutant backgrounds. We reasoned that the proteins targeted by MecA/Clp-dependent proteolysis would exhibit higher abundances in the Δ*mecA* and/or Δ*mecA/clpP* double mutant strains. Indeed, 5 of the 13 MecA-interactome proteins we examined did exhibit discernable regulation by MecA/Clp-dependent proteolysis (GltB, ThiI, DnaG, SpaP and TopA) (Figure 5). ClpC abundance may have been slightly affected as well, but if anything, it seemed to be stimulated by MecA/Clp rather than depleted by it (Figure 5A). In contrast, the abundances of the remaining proteins displayed no apparent differences in the *mecA* and/or *mecA/clpP* backgrounds (Figure 5C, H-O), suggesting that MecA likely regulates much, if not most, of its diverse interactome through an uncharacterized mechanism largely independent of Clp-dependent proteolysis.

**Figure 5.**
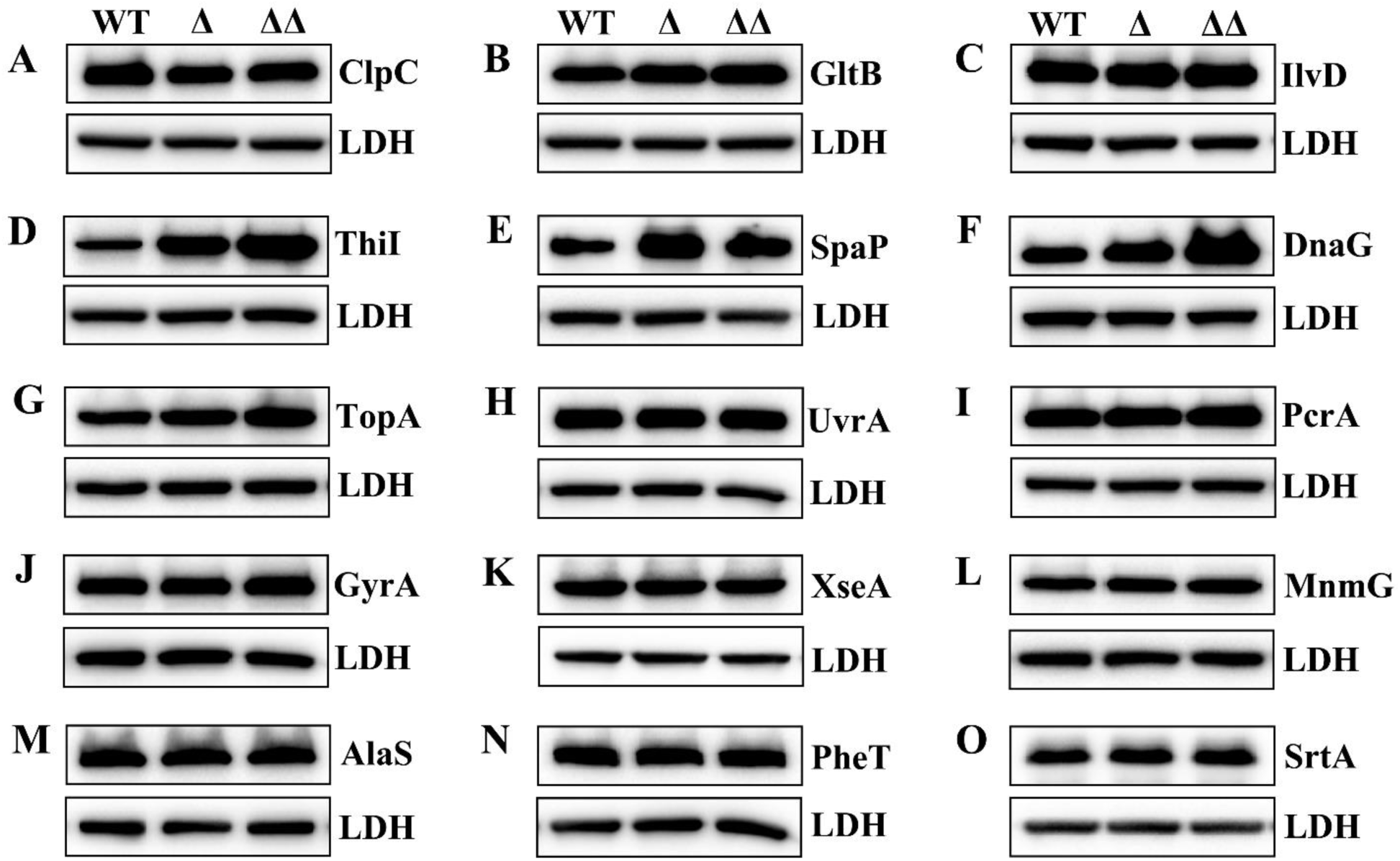
Role of MecA-dependent proteolysis within the MecA interactome. (A – O) The total abundance of MecA-interacting proteins was analyzed in the following backgrounds: wild-type (WT), *mecA* mutant (Δ) and *mecA/clpP* double mutant (ΔΔ). Cultures were grown to mid-logarithmic phase before extracting protein lysates to assess the abundance of MecA-interacting proteins. Lactate dehydrogenase (LDH) was included as a loading control. Functional categories of the assayed proteins: (A) protein quality control, (B and C) amino acid metabolism, (D) tRNA modification, (E) cell adhesion, (F-K) DNA metabolism and transcription, and (L-N) protein synthesis.

## Discussion

Proteins are the workhorses of the cell, performing the vast majority of cellular functions, much of which occur in the context of higher order protein complexes. Consequently, there is a broad interest to better understand the largely hidden ‘social’ nature of proteins, especially those serving as hubs within complex regulatory networks (20–25). In this study, we investigated the interactome of MecA due to its broad conservation, potential role as a major network hub, and the limited number of known MecA interactions. We identified a highly diverse MecA interactome and were able to use this information to develop and validate a new split luciferase complementation assay (SLCA) useful for monitoring PPI in live bacteria. Our SLCA approach employed a synthetic red-shifted Renilla luciferase variant of RLuc8 (61–64) called Green Renilla (RenG), which we have previously demonstrated to yield exceptionally high activity in a variety of bacterial species (46,65). Besides its robust luminescence output, RenG is also particularly well suited for SLCA applications due to its small size (36 kDa) and monomeric functionality. We further developed RenG SLCA to demonstrate its utility as a simple and rapid *in vivo* assessment of PPI avidity within live cells. Since luciferase assays are also ideally suited for temporal studies (50), one could conceivably adapt RenG SLCA to graph kinetic changes in PPI triggered by alterations in growth conditions, signal molecules, and/or environmental stimuli.

The results of our MecA SLCA illustrate the importance of screening different split luciferase fusion combinations when optimizing SLCA for proteins of interest. For our assays, C-terminal fusions generally yielded the strongest signal outputs, but exceptions were noted, such as ClpC (Figure 1E) and DnaG (Figure 2H). In several instances, C-terminal fusions were essential for functionality of the assay (Figure 2D, 2F, 2G, and 2I), but in most cases, the orientations of the fusions simply affected overall signal outputs. MecA protein interactions have been reported to occur with its N-terminal segment (6,10), which is highly consistent with our SLCA results. We found substantially lower SLCA values with N-terminal RenG fusions to MecA (Figure 1E and 1F), presumably due to partial steric interference between the fused luciferase fragment and the MecA interaction partners. We also noted signal output variability depending upon which luciferase fragments were chosen to fuse to the target proteins. However, the magnitude of these differences was generally much smaller, indicating that the location of the luciferase fusion on a target protein is more influential on SLCA output compared to the choice of luciferase fragment. Thus, for a simple determination of +/– protein complex formation, it may not be essential to screen every possible combination of protein chimeras. However, when assessing relative binding affinity via SLCA, determining the optimal orientation of both protein chimeras is recommended to ensure maximal SLCA output. Likewise, all of our chimeras also included a flexible linker to separate the luciferase subunits from their fused proteins. This reduces the chance that a luciferase fusion would impede the proper folding of a target protein. Presumably, the inclusion of linkers would also increase the mobility of the fused luciferase fragments to facilitate their reconstitution following heteromeric complex formation between proteins of interest.

Overall, our SLCA relative binding affinity data appeared to closely mirror independent co-IP measurements of the same protein pairs, supporting the utility of this approach (Table 2 and Figure 4). We noted one potential exception, that of MecA-SpaP. This interaction yielded the weakest relative binding affinity (Table 2), despite the fact that SpaP was enriched in all three of our interactome screens (Table 1). SpaP is a member of the antigen I/II family of cell wall anchored adhesins produced by *S. mutans* and other streptococci (66–71). SpaP contains an N-terminal signal peptide and a C-terminal sortase LPXTG motif used to covalently attach the protein to the cell wall. During its SecA-dependent translocation through the cell membrane, the N-terminal signal sequence of SpaP is removed, while the C-terminal LPXTG motif is later cleaved between the threonine (T) and glycine (G) residues as part of its subsequent attachment to peptidoglycan (72). Consequently, both N- and C-terminal SpaP luciferase fusions only remain as intact chimeras while SpaP is in its immature unprocessed cytoplasmic form. This processing step likely interferes with the proper measurement of relative binding affinity via SLCA because the intact luciferase enzyme can retain its enzymatic function following cleavage from SpaP, whereas free floating luciferase fragments do not (Figure 1D). As a consequence, we suspect that the intact SpaP luciferase fusion yielded a disproportionately strong luciferase signal relative to the SLCA output (Figure 2M and 3B). Thus, protein processing steps are another key consideration when designing SLCA constructs to measure relative binding affinities. It is worth noting that this issue may be addressable by appending a degron to the N- or C-terminal portions of the linker sequence connecting the target protein-luciferase chimera (56). With this configuration, the intact luciferase enzyme should be efficiently targeted for degradation immediately following its cleavage from the chimera, thus limiting reporter activity to the immature unprocessed form of SpaP. Even with this processing issue for the SpaP construct, we still measured SLCA values almost 2 orders of magnitude above background values, indicating a genuine interaction with MecA, as suggested by the original interactome analysis (Table 1 and Figure 2M).

While MecA is a highly conserved pleiotropic regulator, substantial knowledge gaps remain regarding its specific functional role in the cell. MecA has been primarily investigated for its role as an adapter protein for ClpC during assembly of the ClpCP protease complex (6–8,10,14,15,17,60). This ability is critical for stimulating proteolysis by ClpCP (6,7,73,74). However, several studies have also provided evidence to suggest that MecA can affect bacterial physiology through mechanisms independent of ClpCP proteolysis (12,18,19). Therefore, it was of particular interest to determine whether Clp-dependent proteolysis is a common feature of the *S. mutans* MecA interactome. As shown in Figure 5, of the 13 MecA-interacting proteins we assayed, only GltB (glutamate synthase subunit β), ThiI (tRNA 4-thiouridine(8) synthase), DnaG (DNA primase), SpaP (cell wall adhesin), and TopA (type I DNA topoisomerase) exhibited increased abundances in the *mecA* and/or *mecA/clpP* backgrounds, indicating these proteins are likely to be targeted for degradation by MecA/Clp-dependent proteolysis. However, we did not observe evidence of MecA/Clp-dependent proteolysis for the majority of proteins examined, suggesting a large fraction of the MecA interactome is regulated via an unknown mechanism (Figure 5). Furthermore, our MecA interactome data provide strong evidence to support its functional role as a pleiotropic regulator, as we detected a highly diverse interactome that included proteins from a wide variety of functional categories and metabolic pathways (Table 1). Some of these interactions may also partially explain the commonly observed phenotypes associated with *mecA* deletions or overexpression mutations in different organisms. For example, *mecA* is required for normal biofilm formation in multiple organisms including *S. mutans* (12,19). We detected multiple MecA interaction partners that have been reported to influence biofilm development like GltA/B (75–77), SpaP (69,70,78), and the glucosyltransferases GtfB and GtfC (79,80) (Table 1). Cell division and chromosomal defects are also commonly reported MecA phenotypes (18,19). Accordingly, the *S. mutans* MecA interactome includes broadly conserved proteins such as FtsZ (Z-ring formation), FtsQ (divisome formation), SepF (septum formation), GyrA/B (DNA gyrase), PcrA (DNA helicase), TopA (topoisomerase) and others. Given the relatively small fraction of proteins exhibiting MecA/Clp-dependent proteolysis (Figure 5), there is reason to suspect that MecA likely affects a large portion of its interactome by modulating protein function, rather than by simply stimulating proteolytic degradation. The results presented in the current study provide the critical insights needed to further explore how this regulation may be achieved, which would be a key advance in our understanding of MecA as a central hub of multiple protein interaction networks.

## Material and Methods

### Bacterial strains, culture media, and growth conditions

The bacterial strains used in this study are listed in Table S1. All *S. mutans* strains were grown either in an anaerobic chamber containing an atmosphere of 90% N_2_, 5% CO_2_, and 5% H_2_ at 37 °C or in a 5% CO_2_ incubator at 37 °C. *S. mutans* strains were cultured in Todd Hewitt medium (Difco) supplemented with 0.3% (w/v) yeast extract (THYE). For the selection of antibiotic-resistant colonies, THYE plates were supplemented with 1 mg ml^-1^ spectinomycin, 800 *µ*g ml^-1^ kanamycin, 12.5 *µ*g ml^-1^ erythromycin, or 0.02 M DL-4-chlorophenylalanine (4-CP).

### DNA manipulation and strain construction

Primers used in this study are listed in Table S2. Specific details of strain construction are described in Supplemental Experimental Procedures. The protocol for cloning-independent allelic replacements and markerless mutagenesis of *S. mutans* have been described previously (81). Individual PCR reactions were performed using Phusion DNA Polymerase (Thermo Scientific), while overlap extension polymerase chain reaction (OE-PCR) was performed using AccuPrime DNA Polymerase (Life Technologies). The *S. mutans* reference strain UA159 served as the wild-type parental strain for all experiments.

### Coimmunoprecipitation

*S. mutans* strains were cultured anaerobically to mid-log phase at 37 °C in THYE medium containing the appropriate antibiotics. Cells were pelleted and washed with ice cold phosphate-buffered saline (PBS), and then crosslinked in PBS buffer containing 1% (v/v) formaldehyde for 10-15 min at 25 °C before quenching with ice cold 0.125 M glycine. Cells were disrupted with sonication in lysis buffer [50 mM Tris, 150 mM NaCl, 1% (v/v) Triton X-100, protease inhibitor cocktail (Sigma-Aldrich), pH 7.4], and clarified lysates were incubated with anti-FLAG M2 affinity gel (Sigma-Aldrich) for 3 h or overnight with end-over-end rotating at 4 °C. After washing four times with lysis buffer, protein was eluted from the matrix with 100 *µ*l 3×FLAG competitor peptide (150 ng µl^-1^ final concentration) with rotating at 4 °C for 1 h (for TAP tag protein) or incubating with 1% (w/v) SDS with gentle shaking at 25 °C for 5 min. For the purification of TAP tagged proteins, the eluates obtained from competitor peptide elutions were resuspended in lysis buffer and incubated with Pierce^TM^ anti-HA agarose (Thermo Fisher) for 3 h with end-over-end rotating at 4 °C. After washing the resin four times with lysis buffer, protein was eluted with 100 *µ*l 1% (w/v) SDS with gentle shaking at 25 °C for 5 min. The wild-type strain UA159 served as a no-epitope negative control sample. Immunopurified supernatants were analyzed by SDS-polyacrylamide gel electrophoresis (PAGE) and either stained with Coomassie Blue or analyzed by western blot. Prior to performing in-gel digestion for mass spectrometry analysis, a small fraction of each sample was separated via 12% SDS-PAGE to verify sample quantity and quality.

### In-gel digestion and liquid chromatography tandem mass spectrometry analysis

Proteins were separated in NuPAGE^TM^ 10% bis-tris gels (Invitrogen) for 5 min at 150 V and stained with Coomassie Blue. Visible bands were excised from the gel and cut into approximately 1 mm^2^ pieces. In-gel digestion was performed using Trypsin Gold (Promega) and ProteaseMAX^TM^ surfactant (Promega) according to the manufacturer’s protocols. Mass spectrometry analysis was performed by the Oregon Health & Science University (OHSU) proteomics shared resource.

### Luciferase activity assay

Luciferase assays were performed as previously described (46). Overnight *S. mutans* cultures were diluted 1:40 in THYE and grown to mid-log phase. Luciferase activity was detected after adding 1 *µ*l of 0.75 mg ml^-1^ coelenterazine-h solution (NanoLight Technologies) to 100 *u*l of cell culture. Luciferase activity was measured with a GloMAX Discover 96-well luminometer (Promega). Luciferase data were normalized by dividing luciferase values by their corresponding optical density (OD_600_) values. Data are presented as the means ±standard deviations of at least three biological replicates.

### Western blot

*S. mutans* strains were cultured anaerobically to mid-log phase at 37 °C in THYE medium containing the appropriate antibiotics. Cells were pelleted and washed with ice cold PBS, and then resuspended in ice cold TBS buffer containing protease inhibitors. Cells were disrupted with a Precellys Evolution (Bertin Technologies) bead disruptor. Equal amounts of clarified total protein lysates were separated by 12% SDS-PAGE, followed by electroblotting onto nitrocellulose membranes (Bio-Rad). The membranes were blocked with 5% (w/v) non-fat milk in TBS buffer containing 0.1% (v/v) Tween 20 and then incubated with primary antibodies [OctA-Probe antibody (H-5) anti-FLAG, Santa Cruz Biotechnology or Monoclonal anti-HA antibody, Sigma Aldrich] diluted 1:2,000. After incubating with 1:10,000 diluted HRP-conjugated secondary antibodies (Santa Cruz Biotechnology), immunoreactive bands were detected with the ChemiGlow West Chemiluminescence Substrate Kit (Protein Simple). FLAG epitope-labeled lactate dehydrogenase (LDH) served as a loading control.

