## Supplemental material for "Mass spectrometry and split luciferase complementation assays reveal the MecA protein interactome of *Streptococcus mutans*"

### Supplemental Experimental Procedures

#### DNA manipulation and strain construction

The bacterial strains used in this study are listed in Table S1. The protocol for cloning-independent allelic replacements and markerless mutagenesis of *S. mutans* has been described previously (Xie et al, 2011). Individual PCR reactions were performed using Phusion DNA Polymerase (Thermo Scientific), while overlap extension polymerase chain reaction (OE-PCR) was performed using AccuPrime DNA Polymerase (Life Technologies). Primers used in this study is described in Table S2.

#### Construction of TAP tagged *mecA* strain for coimmunoprecipitation analysis

A TAP tag (with triple HA epitope and one FLAG epitope) was translationally fused to the C-terminus of MecA, a flexible linker 2× GGGGS was added between MecA and the TAP tag. Briefly, the flanking upstream and downstream sequences of *mecA* were PCR amplified with the primer pairs MecA-LF/ MecA(TAP)-R and (erm)MecA dn-F/ MecA-RR, respectively. The TAP tag was amplified with the primer pair HA3-FLAG-C-F/ J25 FLAG-R using a synthesized TAP fragment DNA template (Integrated DNA Technologies). An erythromycin resistance cassette *ermB* was amplified from the plasmid pJY4164 (Achen et al, 1986) with the primer pair (TAP)erm-F/erm-R. The four fragments all contained segments of complementary sequence that facilitated their assembly via overlap extension PCR (OE-PCR) with the primer pair MecA-LF/MecA-RR. The resulting amplicons were transformed into the wild-type strain UA159 and selected for erythromycin resistance to produce the strain MecATAP.

#### Construction strains for RNase J1-J2 split luciferase complementation assays

A markerless mutagenesis strategy was employed to translationally fuse the N-terminal domain (NTD1, 1-155 amino acid; NTD2, 1-229 amino acid) of the Green Renilla luciferase (RenG) enzyme to the C-terminus of RNase J1, while the RenG C-terminal domain (CTD1, 156-314 amino acid; CTD2, 230-314 amino acid) was translationally fused to the C-terminus of RNase J2. Both constructs included a flexible linker 2× GGGGS between the protein of interest and the RenG fragment. Construct assembly was performed as follows. The partial ORF and downstream sequence of *rnjA* (RNase J1) were amplified with the primer pairs J1-LF/J1(IFDC2)-R and (IFDC2)J1 dn-F/J1-RR, respectively. The counterselectable IFDC2 cassette was amplified from the plasmid pIFDC2 (Xie et al, 2011) using the primer pair IFDC2-F/IFDC2-R. The resulting PCR amplicons were mixed and assembled via OE-PCR using the primer pair J1-LF/J1-RR. The OE-PCR amplicon was transformed into the strain UA159 and selected for erythromycin resistance to produce the intermediate strain J1IFDC2. Next, the partial ORF and downstream fragment of *rnjA* were amplified with the primer pairs J1-LF/J1-linker-R and J1 dn-F/ J1-RR, respectively. The N-terminal 1-155 amino acid fragment of the Green Renilla luciferase ORF (*renG*) was amplified from the genomic DNA of strain ldhRenGSm (Merritt et al, 2016) with the primer pair (linker)RenG-F-2/RenG155(J1 dn)-R. These three fragments were mixed and assembled via OE-PCR with the primer pair J1-LF/J1-RR to create the first OE-PCR amplicon. The N-terminal 1-229 amino acid fragment of *renG* was PCR amplified from the genomic DNA of strain ldhRenGSm (Merritt et al, 2016) with the primer pair (linker)RenG-F-2/RenG229(J1 dn)-R. Amplicons of the two *rnjA* upstream and downstream homologous flanking sequences and the *renG* 1-229 amplicon were mixed and assembled via OE-PCR with the primer pair J1-LF/J1-RR to create the second OE-PCR amplicon. The two resulting OE-PCR amplicons were separately transformed into the strain J1IFDC2 and selected on THYE medium containing 4-CP to remove the IFDC2 cassette and obtain the strains J1-NTD1 and J1-NTD2, respectively.

The partial ORF and downstream sequence of *rnjB* (RNase J2) were PCR amplified with the primer pairs J2-LF/J2(IFDC2)-R and J2 dn-F/J2-RR, while the IFDC2 cassette was amplified from the plasmid pIFDC2 using the primer pair IFDC2-F/IFDC2-R. The resulting PCR amplicon was mixed and assembled via OE-PCR using the primer pair J2-LF/J2-RR, and then transformed into the strains J1-NTD1 and J1-NTD2 using erythromycin selection to produce the intermediate strains J1NTD1-J2IFDC2 and J1NTD2-J2IFDC2. Next, the partial ORF and downstream fragment of *rnjB* were PCR amplified with the primer pair J2-LF/J2-linker-R and J2 dn-F/J2-RR, respectively. The C-terminal *renG* 156-314 amino acid fragment was amplified from the genomic DNA of strain ldhRenGSm with the primer pair (linker)RenG156-F/RenG156-314(J2 dn)-R. The resulting amplicons were mixed and assembled via OE-PCR using the primer pair J2-LF/J2-RR to generate the first

OE-PCR amplicon. The C-terminal *renG* 230-314 amino acid fragment was PCR amplified from the genomic DNA of strain ldhRenGSm with the primer pair (linker)RenG230-F/RenG156-314(J2 dn)-R. The *renG* C-terminal 230-314 amino acid fragment was mixed with the partial ORF and downstream fragment of *rnjB* and assembled via OE-PCR using the primer pair J2-LF/J2-RR to generate the second OE-PCR amplicon. The first OE-PCR amplicon was transformed into the strain J1NTD1-J2IFDC2, while the second OE-PCR amplicon was transformed into the strain J1NTD2-J2IFDC2. Both transformation reactions were selected on THYE medium containing 4-CP to remove the IFDC2 cassette and obtain the strains J1-NTD1+J2-CTD1 and J1-NTD2+J2-CTD2, respectively.

The partial ORF and downstream sequence of *rnjA* were amplified with the primer pairs J1-LF/J1(rbs-RenG1-155)-R and (erm)J1 dn-F/J1-RR, respectively. The RenG NTD1 (1-155 amino acid) fragment was amplified from the genomic DNA of strain ldhRenGSm with the primer pair (LDH-rbs)RenG-F/RenG1-155(erm)-R-2. An erythromycin resistance cassette *ermB* was amplified from the plasmid pJY4164 with the primer pair erm-F/erm-R. The four fragments were mixed and assembled via OE-PCR with the primer pair J1-LF/J1-RR to generate the first OE-PCR amplicon. Next, the RenG NTD2 (1-229 amino acid) fragment was amplified from the genomic DNA of strain ldhRenGSm with the primer pair (LDH-rbs)RenG-F/RenG1-229(erm)-R. The four amplicons containing the partial ORF and downstream fragment of *rnjA*, NTD2, and the erythromycin resistance cassette were mixed and assembled via OE-PCR with the primer pair J1-LF/J1-RR to generate the second OE-PCR amplicon. The resulting OE-PCR amplicons were transformed into the wild-type strain UA159 and selected for erythromycin resistance to produce the strains NTD1 and NTD2.

The partial ORF and downstream sequence of *rnjB* were PCR amplified with the primer pairs J2-LF/J2(rbs-RenG156-314)-R and (Kan)J2 dn-F-2/J2-RR. The RenG CTD1 (156-314 amino acid) fragment was amplified from the genomic DNA of strain ldhRenGSm with the primer pair (LDH rbs)RenG156-314-F/RenG-R. A kanamycin resistance cassette *aphAIII* was amplified from the plasmid pWVKTs (Gutierrez et al, 1996) with the primer pair (RenG)Kan-F-2/KanR-R-2. The four fragments were mixed and assembled via OE-PCR with the primer pair J2-LF/J2-RR. The resulting OE-PCR amplicon was transformed into the strain NTD1 and selected for kanamycin resistance to generate the strain NTD1 + CTD1.

Next, the partial ORF and downstream sequence of *rnjB* were amplified with the primer pairs J2-LF/J2(LDH rbs)-R and (Kan)J2 dn-F-2/J2-RR. The RenG CTD2 (230-314 amino acid) fragment was amplified from the genomic DNA of strain ldhRenGSm with the primer pair (LDH rbs)RenG230-314-F/RenG-R. A kanamycin resistance cassette *aphAIII* was amplified from the plasmid pWVKTs with the primer pair (RenG)Kan-F-2/KanR-R-2. The four fragments were mixed and assembled via OE-PCR with the primer pair J2-LF/J2-RR. The resulting OE-PCR amplicon was transformed into the strain NTD2 and selected for kanamycin resistance to generate the strain NTD2 + CTD2.

**Construction of strains for MecA-ClpC split luciferase complementation assays**

DNA encoding the RenG N-terminal domain (NTD, 1-155 amino acid) or C-terminal domain (CTD, 156-314 amino acid) was translationally fused either to the C-terminus or N-terminus of MecA, while the C-terminal domain or N-terminal domain of RenG was translationally fused to the either terminus of ClpC. A flexible linker 3× GGGGS was added between the protein of interest and RenG. Construct assembly was performed as follows. The upstream and downstream flanking sequences of *mecA* were PCR amplified with the primer pairs MecA-LF-1/MecA(IFDC2)-R and (IFDC2)MecA dn-F/MecA-RR-1, respectively. The IFDC2 cassette was amplified from the plasmid pIFDC2 using the primer pair IFDC2-F/IFDC2-R. The resulting PCR amplicons were mixed and assembled via OE-PCR using the primer pair MecA-LF-1/MecA-RR-1. The OE-PCR amplicon was transformed into the strains UA159 and ΔM (Qin et al, 2021) and selected for erythromycin resistance to produce the intermediate strains MecAIFDC2 and MecAIFDC2/ΔM. Next, the *mecA* upstream and downstream fragments were PCR amplified with the primer pairs MecA-LF-1/MecA(linker)-R and (RenG1-155)MecA dn-F/MecA-RR-1, respectively. DNA encoding the N-terminal domain of RenG was amplified from the genomic DNA of strain ldhRenGSm with the primer pair (MecA)linker-F/RenG1-155-R. These three amplicons were mixed and assembled via OE-PCR using the primer pair MecA-LF-1/MecA-RR-1 to generate the first OE-PCR amplicon. The *mecA* upstream and downstream fragments were PCR amplified with the primer pairs MecA-LF-1/MecA(linker)-R and (RenG)MecA dn-F/MecA-RR-1, respectively. DNA encoding the C-terminal domain of RenG was amplified from the genomic DNA of strain ldhRenGSm with the primer pair (MecA)linker-F/RenG-R. The resulting PCR amplicons were mixed and assembled via OE-PCR using the primer pair MecA-LF-1/MecA-RR-1 to generate the second OE-PCR amplicon. The OE-PCR amplicons were transformed into the strains MecAIFDC2 and MecAIFDC2/ΔM, and selected on THYE medium containing 4-CP to remove the IFDC2 cassette to obtain the strains MecA-NTD, MecA-CTD, MecA-NTD/ΔM, and MecA-CTD/ΔM.

The upstream and downstream flanking sequences of *mecA* were PCR amplified with the primer pairs MecA-LF-2/MecA up(IFDC2)-R and (IFDC2)MecA-F/MecA-RR-2, respectively. The IFDC2 cassette was amplified from the plasmid pIFDC2 using the primer pair IFDC2-F/IFDC2-R. The resulting PCR amplicons were mixed and assembled via OE-PCR using the primer pair MecA-LF-2/MecA-RR-2. The OE-PCR amplicon was transformed into the strains UA159 and ΔM, and selected for erythromycin resistance to produce the intermediate strains IFDC2-MecA and IFDC2-MecA/ΔM. Next, the upstream and downstream fragments of *mecA* were PCR amplified with the primer pairs MecA-LF-2/MecA up(RenG1-155)-R and (linker)MecA-F/MecA-RR-2, respectively. DNA encoding the N-terminal domain of RenG was amplified from the genomic DNA of strain ldhRenGSm with the primer pair RenG-F-2/RenG1-155(linker)-R. The three resulting PCR amplicons were mixed and assembled via OE-PCR with the primer pair MecA-LF-2/MecA-RR-2 to obtain the first OE-PCR amplicon. Next, the upstream and downstream fragment of *mecA* were PCR amplified with the primer pairs MecA-LF-2/MecA up(RenG156-314)-R and (linker)MecA-F/MecA-RR-2, respectively. DNA encoding the C-terminal domain of RenG was PCR amplified from the genomic DNA of strain ldhRenGSm with the primer pair RenG156-314-F/RenG156-314(linker)-R. The resulting PCR amplicons were mixed and assembled via OE-PCR with the primer pair MecA-LF-2/MecA-RR-2 to obtain the second OE-PCR amplicon. The OE-PCR amplicons were transformed into the strains IFDC2-MecA and IFDC2-MecA/ΔM, and selected on THYE medium containing 4-CP to remove the IFDC2 cassette to obtain the strains NTD-MecA, CTD-MecA, NTD-MecA/ΔM and CTD-MecA/ΔM.

The partial ORF and downstream sequence of *clpC* were PCR amplified with the primer pairs ClpC-LF-2/ClpC(linker)-R and (erm)ClpC dn-F/ClpC-RR-2, respectively. DNA encoding the N-terminal domain of RenG was PCR amplified from the genomic DNA of strain MecA-NTD with the primer pair linker15-F/RenG1-155-R, while the erythromycin resistance cassette *ermB* was amplified from the plasmid pJY4164 with the primer pair (RenG1-155)erm-F/erm-R. These four PCR amplicons were mixed and assembled via OE-PCR with the primer pair ClpC-LF-2/ClpC-RR-2. The resulting amplicon was transformed into the strains MecA-CTD and CTD-MecA, and selected for erythromycin resistance to generate the strains MecA-CTD + ClpC-NTD and CTD-MecA + ClpC-NTD. Next, DNA encoding the C-terminal domain of RenG was PCR amplified from the genomic DNA of strain MecA-CTD with the primer pair linker15-F/RenG-R, while the erythromycin resistance cassette *ermB* was amplified from the plasmid pJY4164 with the primer pair (RenG)erm-F/erm-R. The four amplicons were mixed and assembled via OE-PCR using the primer pair ClpC-LF-2/ClpC-RR-2. The resulting OE-PCR amplicon was transformed individually into the strains MecA-NTD and NTD-MecA, and selected for erythromycin resistance to generate the strains MecA-NTD + ClpC-CTD and NTD-MecA + ClpC-CTD.

The upstream sequence and partial ORF of *clpC* were PCR amplified with the primer pairs ClpC-LF-1/ClpC up(erm)-R and (linker)ClpC-F/ClpC-RR-1, respectively. An erythromycin resistance cassette *ermB* was PCR amplified from the plasmid pJY4164 with the primer pair erm-F/erm(rbs)-R-1. DNA encoding the N-terminal domain of RenG was PCR amplified from the genomic DNA of strain MecA-NTD with the primer pair (rbs)RenG-F/RenG1-155(linker)-R. The PCR amplicons were mixed and assembled via OE-PCR using the primer pair ClpC-LF-1/ClpC-RR-1. The resulting OE-PCR amplicon was transformed into the strains MecA-CTD and CTD-MecA, and selected for erythromycin resistance to obtain the strains MecA-CTD + NTD-ClpC, CTD-MecA + NTD-ClpC. Next, the erythromycin resistance cassette *ermB* was amplified from the plasmid pJY4164 with the primer pair erm-F/erm(rbs)-R-2. DNA encoding the C-terminal domain of RenG was amplified from the genomic DNA of strain MecA-CTD with the primer pair (rbs)RenG156-314-F/RenG156-314(linker)-R. The four amplicons were mixed and assembled via OE-PCR using the primer pair ClpC-LF-1/ClpC-RR-1. The resulting OE-PCR amplicon was transformed into the strains MecA-NTD and NTD-MecA, and selected for erythromycin resistance to obtain the strains MecA-NTD + CTD-ClpC and NTD-MecA + CTD-ClpC.

**Construction of strains for MecA-ComX split luciferase complementation assays**

The RenG N-terminal domain (NTD, 1-155 amino acid) or C-terminal domain (CTD, 156-314 amino acid) was translationally fused to either the C-terminus or N-terminus of MecA, while the RenG C-terminal domain or N-terminal domain was translationally fused to either terminus of ComX. A flexible linker 3× GGGGS was inserted between the protein of interest and the RenG fragment. Constructs were assembled as follows. The upstream and downstream of flanking sequences of *comX* were PCR amplified with the primer pairs ComX-LF/ComX(IFDC2)-R and (IFDC2)ComX dn-F/ComX-RR, respectively. The IFDC2 cassette was amplified from the plasmid pIFDC2 using the primer pair IFDC2-F/IFDC2-R. The resulting PCR amplicons were mixed and assembled via OE-PCR using the primer pair ComX-LF/ComX-RR. The OE-PCR amplicon was transformed into the strains MecA-NTD/ΔM, MecA-CTD/ΔM, NTD-MecA/ΔM and CTD-MecA/ΔM and selected for erythromycin resistance to produce the intermediate strains MecA-NTD + ComX-IFDC2/ΔM, MecA-CTD + ComX-IFDC2/ΔM, NTD-MecA + ComX-IFDC2/ΔM and CTD-MecA + ComX-IFDC2/ΔM. Next, the upstream and downstream flanking sequences of *comX* were PCR amplified with the primer pairs ComX-LF/ComX(linker)-R and (RenG1-155)ComX dn-F/ComX-RR, respectively. DNA encoding the N-terminal domain of RenG was amplified from the genomic DNA of strain MecA-NTD with the primer pair linker15-F/RenG1-155-R. The PCR amplicons were mixed and assembled via OE-PCR using the primer pair ComX-LF/ComX-RR. The resulting OE-PCR amplicon was transformed into the strains MecA-CTD + ComX-IFDC2/ΔM and CTD-MecA + ComX-IFDC2/ΔM and selected on THYE medium containing 4-CP to remove the IFDC2 cassette and obtain the strains MecA-CTD + ComX-NTD and CTD-MecA + ComX-NTD. DNA encoding the C-terminal domain of RenG was amplified from the genomic DNA of strain MecA-CTD with the primer pair linker15-F/RenG-R. The upstream and downstream sequences of *comX* were amplified with the primer pairs ComX-LF/ComX(linker)-R and (RenG)ComX dn-F, respectively. These three fragments were mixed and assembled via OE-PCR using primer pair ComX-LF/ComX-RR. The resulting OE-PCR amplicon was transformed into the strains MecA-NTD + ComX-IFDC2/ΔM and NTD-MecA + ComX-IFDC2/ΔM and selected on THYE medium containing 4-CP to remove the IFDC2 cassette to obtain the strains MecA-NTD + ComX-CTD and NTD-MecA + ComX-CTD.

The upstream and downstream flanking sequences of *comX* were PCR amplified with the primer pairs ComX-LF/ComX up(IFDC2)-R and (IFDC2)ComX-F/ComX-RR, respectively. The IFDC2 cassette was amplified from the plasmid pIFDC2 using the primer pair IFDC2-F/IFDC2-R. The resulting PCR amplicons were mixed and assembled via OE-PCR using the primer pair ComX-LF/ComX-RR. The OE-PCR amplicon was transformed into the strains MecA-NTD/ΔM, MecA-CTD/ΔM, NTD-MecA/ΔM, and CTD-MecA/ΔM, and selected for erythromycin resistance to produce the intermediate strains MecA-NTD + IFDC2-ComX /ΔM, MecA-CTD + IFDC2-ComX /ΔM, NTD-MecA + IFDC2-ComX /ΔM and CTD-MecA + IFDC2-ComX /ΔM. Next, the upstream and downstream of flanking sequences of *comX* were amplified with the primer pairs ComX-LF/ComX up(RenG1-155)-R and (linker)ComX-F/ComX-RR, respectively. DNA encoding the RenG N-terminal domain was amplified from the genomic DNA of strain MecA-NTD with the primer pair RenG-F-2/RenG1-155(linker)-R. These three amplicons were mixed and assembled via OE-PCR using the primer pair ComX-LF/ComX-RR. The resulting OE-PCR amplicon was transformed into the strains MecA-CTD + IFDC2-ComX /ΔM and CTD-MecA + IFDC2-ComX /ΔM, and selected on THYE medium containing 4-CP to remove the IFDC2 cassette and obtain the strains MecA-CTD + NTD-ComX and CTD-MecA + NTD-ComX. Next, the upstream and downstream flanking sequences of *comX* were PCR amplified with the primer pairs ComX-LF/ComX up (RenG156-314)-R and (linker)ComX-F/ComX-RR. DNA encoding the RenG C-terminal domain was PCR amplified from the genomic DNA of strain MecA-CLuc with the primer pair RenG156-314-F/RenG156-314(linker)-R. These three PCR amplicons were mixed and assembled via OE-PCR using the primer pair ComX-LF/ComX-RR. The resulting OE-PCR amplicon was transformed into the strains MecA-NTD + IFDC2-ComX /ΔM and NTD-MecA + IFDC2-ComX /ΔM, and selected on THYE medium containing 4-CP to remove the IFDC2 cassette and obtain the strains MecA-NTD + CTD-ComX and NTD-MecA + CTD-ComX.

**Construction of strains for MecA-ThiI split luciferase complementation assays**

The partial ORF and downstream sequence of *thiI* were PCR amplified with the primer pairs ThiI-LF-1/ThiI(linker)-R and (erm)ThiI dn-F/ThiI-RR-1, respectively. DNA encoding the RenG C-terminal domain was PCR amplified from the genomic DNA of strain MecA-CTD with the primer pair linker15-F/RenG-R, while an erythromycin resistance cassette *ermB* was amplified from the plasmid pJY4164 with the primer pair (RenG)erm-F/erm-R. These four PCR amplicons were mixed and assembled via OE-PCR using the primer pair ThiI-LF-1/ThiI-RR-1 to generate the first OE-PCR product. Next, the upstream sequence and the partial ORF of *thiI* were PCR amplified with the primer pairs ThiI-LF-2/ThiI up(erm)-R and (linker)ThiI-F/ThiI-RR-2, respectively, while an erythromycin resistance cassette *ermB* was amplified from the plasmid pJY4164 with the primer pair erm-F/erm(ThiI rbs)-R. DNA encoding the RenG C-terminal domain was PCR amplified from the genomic DNA of strain CLuc-N-MecA with the primer pair (ThiI rbs)RenG156-F/linker15-R. These four PCR amplicons were mixed and assembled via OE-PCR using the primer pair thiI-LF-2/thiI-RR-2 to generate the second OE-PCR product. The two

OE-PCR amplicons were separately transformed into the strains MecA-NTD and NTD-MecA, and selected for erythromycin resistance to obtain the strains MecA-NTD + ThiI-CTD, MecA-NTD + CTD-ThiI, NTD-MecA + ThiI-CTD and NTD-MecA + CTD-ThiI.

**Construction of strains for MecA-GltB split luciferase complementation assays**

The partial ORF and downstream sequence of *gltB* were PCR amplified with the primer pairs GltB-LF-2/GltB(IFDC2)-R and (IFDC2)GltB dn-F/GltB-RR-2, respectively. The IFDC2 cassette was amplified from the plasmid pIFDC2 using the primer pair IFDC2-F/IFDC2-R. These three PCR amplicons were mixed and assembled via OE-PCR using the primer pair GltB-LF-2/GltB-RR-2. The resulting PCR amplicon was transformed into the strains MecA-NTD and NTD-MecA, and selected for erythromycin resistance to obtain the intermediate strains MecA-NTD + GltB-IFDC2 and NTD-MecA + GltB-IFDC2. Next, the partial ORF and downstream sequence of *gltB* were PCR amplified with the primer pairs GltB-LF-2/GltB(linker)-R and (RenG)GltB dn-F/GltB-RR-2, respectively. DNA encoding the RenG C-terminal domain was PCR amplified from the genomic DNA of strain MecA-CTD with the primer pair linker15-F/RenG-R. These three amplicons were mixed and assembled via OE-PCR using primer pair GltB-LF-2/GltB-RR-2. The resulting OE-PCR amplicon was transformed into the strains MecA-NTD + GltB-IFDC2 and NTD-MecA + GltB-IFDC2, and selected on THYE medium containing 4-CP to remove the IFDC2 cassette and obtain the strains MecA-NTD + GltB-CTD and NTD-MecA + GltB-CTD.

The upstream sequence and the partial ORF of *gltB* were amplified with the primer pairs GltB-LF-1/GltB up(IFDC2)-R and (IFDC2)GltB-F/GltB-RR-1, respectively. The IFDC2 cassette was amplified from the plasmid pIFDC2 using the primer pair IFDC2-F/IFDC2-R. These three PCR amplicons were mixed and assembled via OE-PCR using the primer pair GltB-LF-1/GltB-RR-1. The resulting OE-PCR amplicon was transformed into the strains MecA-NTD and NTD-MecA, and selected for erythromycin resistance to obtain the intermediate strains MecA-NTD + IFDC2-GltB and NTD-MecA + IFDC2-GltB. Next, the upstream sequence and the partial ORF of *gltB* were PCR amplified with the primer pairs GltB-LF-1/GltB up(RenG156-314)-R and (linker)GltB-F/GltB-RR-1, respectively. DNA encoding the RenG C-terminal domain was PCR amplified from the genomic DNA of strain CTD-MecA with the primer pair RenG156-314-F/linker15-R. These three PCR amplicons were mixed and assembled via OE-PCR using the primer pair GltB-LF-1/GltB-RR-1. The resulting OE-PCR amplicon was transformed into the strains MecA-NTD + IFDC2-GltB and NTD-MecA + IFDC2-GltB, and selected on THYE medium containing 4-CP to remove the IFDC2 cassette and obtain the strains MecA-NTD + CTD-GltB and NTD-MecA + CTD-GltB.

**Construction of strains for MecA-AlaS split luciferase complementation assays**

The partial ORF and downstream sequence of *alaS* were PCR amplified with the primer pairs AlaS-LF-1/AlaS(linker)-R and (kan)AlaS dn-F/AlaS-RR-1, respectively. DNA encoding the RenG C-terminal domain was PCR amplified from the genomic DNA of strain MecA-CTD with the primer pair linker15-F/RenG-R, while a kanamycin resistance cassette *aphAIII* was amplified from the plasmid pWVKTs with the primer pair (RenG)kan-F-2/kanR-R-2. These four PCR amplicons were mixed and assembled via OE-PCR using the primer pair AlaS-LF-1/AlaS-RR-1. The resulting OE-PCR amplicon was transformed into the strain MecA-NTD and selected for kanamycin resistance to generate the strain MecA-NTD + AlaS-CTD.

The upstream sequence and partial ORF of *alaS* were PCR amplified with the primer pairs AlaS-LF-2/AlaS up(IFDC2)-R and (IFDC2)AlaS up-F/AlaS-RR-2, respectively. The IFDC2 cassette was amplified from the plasmid pIFDC2 using the primer pair IFDC2-F/IFDC2-R. These three PCR amplicons were mixed and assembled via OE-PCR using the primer pair AlaS-LF-2/AlaS-RR-2. The resulting OE-PCR amplicon was transformed into the strain MecA-NTD and selected for erythromycin resistance to generate the intermediate strain MecA-NTD + IFDC2-AlaS. Next, the upstream sequence and partial ORF of *alaS* were PCR amplified with the primer pair AlaS-LF-2/AlaS up(RenG156-314)-R and (linker)AlaS-F/AlaS-RR-2, respectively. DNA encoding the RenG C-terminal domain was PCR amplified from the genomic DNA of strain CLuc-N-MecA with the primer pair RenG156-314-F/linker15-R. These three PCR amplicons were mixed and assembled via OE-PCR using the primer pair AlaS-LF-2/AlaS-RR-2. The resulting OE-PCR amplicon was transformed into the strain MecA-NTD + IFDC2-AlaS and selected on THYE medium containing 4-CP to remove the IFDC2 cassette and generate the strain MecA-NTD + CTD-AlaS.

**Construction of strains for MecA-DnaG split luciferase complementation assays**

The partial ORF and the downstream sequence of *dnaG* were PCR amplified with the primer pairs DnaG-LF/DnaG(linker)-R and (kan)DnaG dn-F-2/DnaG-RR, respectively. DNA encoding the RenG C-terminal domain was PCR amplified from the genomic DNA of strain MecA-CTD with the primer pair linker15-F/RenG-R, while a kanamycin resistance cassette *aphAIII* was amplified from the plasmid pWVKTs with the primer pair (RenG)kan-F-2/kanR-R-2. These four PCR amplicons were mixed and assembled via OE-PCR using the primer pair DnaG-LF/DnaG-RR. The resulting OE-PCR amplicon was transformed into the strain MecA-NTD and selected for kanamycin resistance to obtain the strain MecA-NTD + DnaG-CTD.

The upstream sequence and the partial ORF of *dnaG* were PCR amplified with the primer pairs DnaG-LF-2/DnaG up(IFDC2)-R and (IFDC2)DnaG up-F/DnaG-RR-2, respectively. The IFDC2 cassette was amplified from the plasmid pIFDC2 using the primer pair IFDC2-F/IFDC2-R. These three PCR amplicons were mixed and assembled via OE-PCR using the primer pair DnaG-LF-2/DnaG-RR-2. The resulting OE-PCR amplicon was transformed into the strains UA159 and MecA-NTD, and selected for erythromycin resistance to generate the intermediate strains IFDC2-DnaG and MecA-NTD + IFDC2-DnaG. Next, the upstream sequence and the partial ORF of *dnaG* were PCR amplified with the primer pairs DnaG-LF-2/DnaG up(RenG156-314)-R and (linker)DnaG-F/DnaG-RR-2, respectively. DNA encoding the RenG C-terminal domain was PCR amplified from the genomic DNA of strain CTD-MecA with the primer pair RenG156-314-F/linker15-R. These three amplicons were mixed and assembled via OE-PCR using the primer pair DnaG-LF-2/DnaG-RR-2. The resulting OE-PCR amplicon was transformed into the strain MecA-NTD + IFDC2-DnaG and selected on THYE medium containing 4-CP to remove the IFDC2 cassette and obtain the strain MecA-NTD + CTD-DnaG.

**Construction of strains for MecA-GyrA split luciferase complementation assays**

The partial ORF and the downstream sequence of *gyrA* were PCR amplified with the primer pairs GyrA-LF/GyrA(linker)-R and (kan)GyrA dn-F/GyrA-RR, respectively. DNA encoding the RenG C-terminal domain was PCR amplified from the genomic DNA of strain MecA-CTD with the primer pair linker15-F/RenG-R and a kanamycin resistance cassette *aphAIII* was amplified from the plasmid pWVKTs with the primer pair (RenG)kan-F-2/kanR-R-2. These four amplicons were mixed and assembled via OE-PCR using the primer pair GyrA-LF/GyrA-RR. The resulting OE-PCR amplicon was transformed into the strain MecA-NTD and selected for kanamycin resistance to generate the strain MecA-NTD + GyrA-CTD.

**Construction of strains for MecA-IlvD split luciferase complementation assays**

The partial ORF and the downstream sequence of *ilvD* were PCR amplified with the primer pairs IlvD-LF/IlvD(linker)-R and (kan)IlvD dn-F/IlvD-RR, respectively. DNA encoding the RenG C-terminal domain together with a kanamycin resistance cassette *aphAIII* fragment was amplified from the genomic DNA of the strain MecA-NTD + AlaS-CTD with the primer pair linker15-F/KanR-R-2. These three PCR amplicons were mixed and assembled via OE-PCR using the primer pair IlvD-LF/IlvD-RR. The resulting OE-PCR amplicon was transformed into the strain MecA-NTD and selected for kanamycin resistance to generate the strain MecA-NTD + IlvD-CTD.

The upstream sequence and the partial ORF of *ilvD* were PCR amplified with the primer pairs IlvD-LF-2/IlvD up(IFDC2)-R and (IFDC2)IlvD up-F/IlvD-RR-2, respectively. The IFDC2 cassette was amplified from the plasmid pIFDC2 using the primer pair IFDC2-F/IFDC2-R. These three PCR amplicons were mixed and assembled via OE-PCR using the primer pair IlvD-LF-2/IlvD-RR-2. The resulting OE-PCR amplicon was transformed into the strain MecA-NTD and selected for erythromycin resistance to obtain the intermediate strain MecA-NTD + IFDC2-IlvD. Next, the upstream sequence and the partial ORF of *ilvD* were PCR amplified with the primer pairs IlvD-LF-2/IlvD up(RenG156-314)-R and (linker)IlvD-F/IlvD-RR-2, respectively. DNA encoding the RenG C-terminal domain was PCR amplified from the genomic DNA of strain CTD-MecA with the primer pair RenG156-314-F/linker15-R. These three PCR amplicons were mixed and assembled via OE-PCR using the primer pair IlvD-LF-2/IlvD-RR-2. The resulting OE-PCR amplicon was transformed into the strain

MecA-NTD + IFDC2-IlvD and selected on THYE medium containing 4-CP to remove the IFDC2 cassette and obtain the strain MecA-NTD + CTD-IlvD.

**Construction of strains for MecA-MnmG split luciferase complementation assays**

The partial ORF and the downstream sequence of *mnmG* were PCR amplified with the primer pairs MnmG-LF/MnmG(linker)-R and (kan)MnmG dn-F/MnmG-RR, respectively. DNA encoding the RenG C-terminal domain together with a kanamycin resistance cassette *aphAIII* fragment was PCR amplified from the genomic DNA of the strain MecA-NTD + AlaS-CTD with the primer pair linker15-F/KanR-R-2. These three PCR amplicons were mixed and assembled via OE-PCR using the primer pair MnmG-LF/MnmG-RR. The resulting OE-PCR amplicon was transformed into the strain MecA-NTD and selected for kanamycin resistance to generate the strain MecA-NTD + MnmG-CTD.

The upstream sequence and the partial ORF of *mnmG* were PCR amplified with the primer pairs MnmG-LF-2/MnmG up(IFDC2)-R and (IFDC2)MnmG up-F/MnmG-RR-2, respectively. The IFDC2 cassette was amplified from the plasmid pIFDC2 using the primer pair IFDC2-F/IFDC2-R. These three PCR amplicons were mixed and assembled via OE-PCR using the primer pair MnmG-LF-2/MnmG-RR-2. The resulting OE-PCR amplicon was transformed into the strain MecA-NTD and selected for erythromycin resistance to obtain the intermediate strain MecA-NTD + IFDC2-MnmG. Next, the upstream sequence and the partial ORF of *mnmG* were PCR amplified with the primer pairs MnmG-LF-2/MnmG up(RenG156-314)-R and (linker)MnmG-F/MnmG-RR-2, respectively. DNA encoding the RenG C-terminal domain was PCR amplified from the genomic DNA of strain CTD-MecA with the primer pair RenG156-314-F/linker15-R. These three PCR amplicons were mixed and assembled via OE-PCR using the primer pair MnmG-LF-2/MnmG-RR-2. The resulting OE-PCR amplicon was transformed into the strain MecA-NTD + IFDC2-MnmG and selected on THYE medium containing 4-CP to remove the IFDC2 cassette and obtain the strain MecA-NTD + CTD-MnmG.

**Construction of strains for MecA-PcrA split luciferase complementation assays**

The partial ORF and the downstream sequence of *pcrA* were PCR amplified with the primer pairs PcrA-LF/PcrA(linker)-R and (kan)PcrA dn-F/PcrA-RR, respectively. DNA encoding the RenG C-terminal domain together with a kanamycin resistance cassette *aphAIII* was PCR amplified from the genomic DNA of the strain MecA-NTD + AlaS-CTD with the primer pair linker15-F/KanR-R-2. These three PCR products were mixed and assembled via OE-PCR using the primer pair PcrA-LF/PcrA-RR. The resulting OE-PCR amplicon was transformed into the strain MecA-NTD and selected for kanamycin resistance to generate the strain MecA-NTD + PcrA-CTD.

The upstream sequence and the partial ORF of *pcrA* were PCR amplified with the primer pairs PcrA-LF-2/PcrA up(IFDC2)-R and (IFDC2)PcrA up-F/PcrA-RR-2, respectively. The IFDC2 cassette was amplified from the plasmid pIFDC2 using the primer pair IFDC2-F/IFDC2-R. These three PCR amplicons were mixed and assembled via OE-PCR using the primer pair PcrA-LF-2/PcrA-RR-2. The resulting OE-PCR amplicon was transformed into the strain MecA-NTD and selected for erythromycin resistance to obtain the intermediate strain MecA-NTD + IFDC2-PcrA. Next, the upstream sequence and the partial ORF of *pcrA* were PCR amplified with the primer pairs PcrA-LF-2/PcrA up(RenG156-314)-R and (linker)PcrA-F/PcrA-RR-2, respectively. DNA encoding the RenG C-terminal domain was PCR amplified from the genomic DNA of strain CTD-MecA with the primer pair RenG156-314-F/linker15-R. These three PCR amplicons were mixed and assembled via OE-PCR using the primer pair PcrA-LF-2/PcrA-RR-2. The resulting OE-PCR amplicon was transformed into the strain MecA-NTD + IFDC2-PcrA and selected on THYE medium containing 4-CP to remove the IFDC2 cassette and obtain the strain MecA-NTD + CTD-PcrA.

**Construction of strains for MecA-PheT split luciferase complementation assays**

The partial ORF and the downstream sequence of *pheT* were PCR amplified with the primer pairs PheT-LF/PheT(linker)-R and (kan)PheT dn-F/PheT-RR, respectively. DNA encoding the RenG C-terminal domain together with a kanamycin resistance cassette *aphAIII* fragment was PCR amplified from the genomic DNA of the strain MecA-NTD + AlaS-CTD with the primer pair linker15-F/KanR-R-2. These three PCR amplicons were mixed and assembled via OE-PCR using the primer pair PheT-LF/PheT-RR. The resulting OE-PCR amplicon was transformed into the strain MecA-NLuc and selected for kanamycin resistance to generate the strain MecA-NTD + PheT-CTD.

**Construction of strains for MecA-SpaP split luciferase complementation assays**

The partial ORF and the downstream sequence of *spaP* were PCR amplified with the primer pairs SpaP-LF/SpaP(linker)-R and (kan)SpaP dn-F/SpaP-RR, respectively. DNA encoding the RenG C-terminal domain together with a kanamycin resistance cassette *aphAIII* fragment was amplified from the genomic DNA of the strain MecA-NTD + AlaS-CTD with the primer pair linker15-F/KanR-R-2. These three PCR amplicons were mixed and assembled via OE-PCR using the primer pair SpaP-LF/SpaP-RR. The resulting OE-PCR amplicon was transformed into the strain MecA-NTD and selected for kanamycin resistance to generate the strain MecA-NTD + SpaP-CTD.

The upstream sequence and the partial ORF of *spaP* were PCR amplified with the primer pairs SpaP-LF-2/SpaP up(IFDC2)-R and (IFDC2)SpaP up-F/SpaP-RR-2, respectively. The IFDC2 cassette was amplified from the plasmid pIFDC2 using the primer pair IFDC2-F/IFDC2-R. These three PCR amplicons were mixed and assembled via OE-PCR using the primer pair SpaP-LF-2/SpaP-RR-2. The resulting OE-PCR amplicon was transformed into the strain MecA-NTD and selected for erythromycin resistance to obtain the intermediate strain MecA-NTD + IFDC2-SpaP. Next, the upstream sequence and the partial ORF of *spaP* were PCR amplified with the primer pairs SpaP-LF-2/SpaP up(RenG156-314)-R and (linker)SpaP-F/SpaP-RR-2, respectively. DNA encoding the RenG C-terminal domain was PCR amplified from the genomic DNA of strain CTD-MecA with the primer pair RenG156-314-F/linker15-R. These three PCR amplicons were mixed and assembled via OE-PCR using the primer pair SpaP-LF-2/SpaP-RR-2. The resulting OE-PCR amplicon was transformed into the strain MecA-NTD + IFDC2-SpaP and selected on THYE medium containing 4-CP to remove the IFDC2 cassette and obtain the strain MecA-NTD + CTD-SpaP.

**Construction of strains for MecA-TopA split luciferase complementation assays**

The partial ORF and the downstream sequence of *topA* were PCR amplified with the primer pairs TopA-LF/TopA(linker)-R and (kan)TopA dn-F/TopA-RR, respectively. DNA encoding the RenG C-terminal domain together with a kanamycin resistance cassette *aphAIII* was PCR amplified from the genomic DNA of the strain MecA-NTD + AlaS-CTD with the primer pair linker15-F/KanR-R-2. These three PCR amplicons were mixed and assembled via OE-PCR using the primer pair TopA-LF/TopA-RR. The resulting OE-PCR amplicon was transformed into the strain MecA-NTD and selected for kanamycin resistance to generate the strain MecA-NTD + TopA-CTD.

The upstream sequence and the partial ORF of *topA* were PCR amplified with the primer pairs TopA-LF-2/TopA up(IFDC2)-R and (IFDC2)TopA up-F/TopA-RR-2, respectively. The IFDC2 cassette was amplified from the plasmid pIFDC2 using the primer pair IFDC2-F/IFDC2-R. These three PCR amplicons were mixed and assembled via OE-PCR using the primer pair TopA-LF-2/TopA-RR-2. The resulting OE-PCR amplicon was transformed into the strain MecA-NTD and selected for erythromycin resistance to obtain the intermediate strain MecA-NTD + IFDC2-TopA. Next, the upstream sequence and the partial ORF of *topA* were PCR amplified with the primer pairs TopA-LF-2/TopA up(RenG156-314)-R and (linker)TopA-F/TopA-RR-2, respectively. DNA encoding the RenG C-terminal domain was PCR amplified from the genomic DNA of strain CTD-MecA with the primer pair RenG156-314-F/linker15-R. These three PCR amplicons were mixed and assembled via OE-PCR using the primer pair TopA-LF-2/TopA-RR-2. The resulting OE-PCR amplicon was transformed into the strain MecA-NTD + IFDC2-TopA and selected on THYE medium containing 4-CP to remove the IFDC2 cassette and obtain the strain MecA-NTD + CTD-TopA.

**Construction of strains for MecA-UvrA split luciferase complementation assays**

The partial ORF and the downstream sequence of *uvrA* were PCR amplified with the primer pairs UvrA-LF/UvrA(linker)-R and (kan)UvrA dn-F/UvrA-RR, respectively. DNA encoding the RenG C-terminal domain together with a kanamycin resistance cassette *aphAIII* was PCR amplified from the genomic DNA of the strain MecA-NTD + AlaS-CTD with the primer pair linker15-F/KanR-R-2. These three PCR amplicons were mixed and assembled via OE-PCR using the primer pair UvrA-LF/UvrA-RR. The resulting OE-PCR amplicon was transformed into the strain MecA-NTD and selected for kanamycin resistance to generate the strain MecA-NTD + UvrA-CTD.

The upstream sequence and the partial ORF of *uvrA* were PCR amplified with the primer pairs UvrA-LF-2/UvrA up(IFDC2)-R and (IFDC2)UvrA up-F/UvrA-RR-2, respectively. The IFDC2 cassette was amplified from the plasmid pIFDC2 using the primer pair IFDC2-F/IFDC2-R. These three PCR amplicons were mixed and assembled via OE-PCR using the primer pair UvrA-LF-2/UvrA-RR-2. The resulting OE-PCR amplicon was transformed into the strain MecA-NTD and selected for erythromycin resistance to obtain the intermediate strain MecA-NTD + IFDC2-UvrA. Next, the upstream sequence and the partial ORF of *uvrA* were PCR amplified with the primer pairs UvrA-LF-2/UvrA up(RenG156-314)-R and (linker)UvrA-F/UvrA-RR-2, respectively. DNA encoding the RenG C-terminal domain was amplified from the genomic DNA of strain CTD-MecA with the primer pair RenG156-314-F/linker15-R. These three PCR amplicons were mixed and assembled via OE-PCR using the primer pair UvrA-LF-2/UvrA-RR-2. The resulting OE-PCR amplicon was transformed into the strain MecA-NTD + IFDC2-TopA and selected on THYE medium containing 4-CP to remove the IFDC2 cassette and obtain the strain MecA-NTD + CTD-UvrA.

**Construction of strains for MecA-XseA split luciferase complementation assays**

The partial ORF and the downstream sequence of *xseA* were PCR amplified with the primer pairs XseA-LF/XseA(linker)-R and (kan)XseA dn-F/XseA-RR, respectively. DNA encoding the RenG C-terminal domain together with a kanamycin resistance cassette *aphAIII* was amplified from the genomic DNA of the strain MecA-NTD + AlaS-CTD with the primer pair linker15-F/KanR-R-2. These three PCR amplicons were mixed and assembled via OE-PCR using the primer pair XseA-LF/XseA-RR. The resulting PCR amplicon was transformed into the strain MecA-NLuc and selected for kanamycin resistance to generate the strain MecA-NTD + XseA-CTD.

The upstream sequence and the partial ORF of *xseA* were PCR amplified with the primer pairs XseA-LF-2/XseA up(IFDC2)-R and (IFDC2)XseA up-F/XseA-RR-2, respectively. The IFDC2 cassette was amplified from the plasmid pIFDC2 using the primer pair IFDC2-F/IFDC2-R. These three PCR amplicons were mixed and assembled via OE-PCR using the primer pair XseA-LF-2/XseA-RR-2. The resulting OE-PCR amplicon was transformed into the strain MecA-NTD and selected for erythromycin resistance to obtain the intermediate strain MecA-NTD + IFDC2-XseA. Next, the upstream sequence and the partial ORF of *uvrA* were PCR amplified with the primer pairs XseA-LF-2/XseA up(RenG156-314)-R and (linker)XseA-F/XseA-RR-2, respectively. DNA encoding the RenG C-terminal domain was PCR amplified from the genomic DNA of strain CTD-MecA with the primer pair RenG156-314-F/linker15-R. These three PCR amplicons were mixed and assembled via OE-PCR using the primer pair XseA-LF-2/XseA-RR-2. The resulting PCR amplicon was transformed into the strain MecA-NTD + IFDC2-TopA and selected on THYE medium containing 4-CP to remove the IFDC2 cassette and obtain the strain MecA-NTD + CTD-N-XseA.

**Construction of strains for MecA-SrtA split luciferase complementation assays**

The upstream and downstream sequences of *srtA* were PCR amplified with the primer pairs SrtA-LF/SrtA dn(IFDC2)-R and (IFDC2)SrtA dn-F/SrtA-RR, respectively. The IFDC2 cassette was amplified from the plasmid pIFDC2 using the primer pair IFDC2-F/IFDC2-R. These three PCR amplicons were mixed and assembled via OE-PCR using the primer pair SrtA-LF/SrtA-RR. The resulting OE-PCR amplicon was transformed into the strains MecA-NTD and NTD-MecA, and selected for erythromycin resistance to obtain the intermediate strains MecA-NTD + SrtA-IFDC2 and NTD-MecA + SrtA-IFDC2. Next, the upstream and downstream sequences of *srtA* were PCR amplified with the primer pairs SrtA-LF/SrtA(linker)-R and (RenG)SrtA dn-F/SrtA-RR, respectively. DNA encoding the RenG C-terminal domain was PCR amplified from the genomic DNA of strain MecA-CTD with the primer pair linker15-F/RenG-R. These three amplicons were mixed and assembled via OE-PCR using primer pair SrtA-LF/SrtA-RR. The resulting OE-PCR amplicon was transformed into the strains MecA-NTD + SrtA-IFDC2 and NTD-MecA + SrtA-IFDC2, and selected on THYE medium containing 4-CP to remove the IFDC2 cassette and obtain the strains MecA-NTD + SrtA-CTD and NTD-MecA + SrtA-CTD.

The upstream and downstream sequences of *srtA* were PCR amplified with the primer pairs SrtA-LF-2/SrtA up(IFDC2)-R and (IFDC2)srtA-F/SrtA-RR-2, respectively. The IFDC2 cassette was amplified from the plasmid pIFDC2 using the primer pair IFDC2-F/IFDC2-R. These three PCR amplicons were mixed and assembled via OE-PCR using the primer pair SrtA-LF-2/SrtA-RR-2. The resulting PCR amplicon was transformed into the strains MecA-NTD and NTD-MecA, and selected for erythromycin resistance to obtain the intermediate strains MecA-NTD + IFDC2-SrtA and NTD-MecA + IFDC2-SrtA. Next, the upstream and downstream sequences of *srtA* were PCR amplified with the primer pairs SrtA-LF-2/SrtA up(RenG156-314)-R and (linker)SrtA-F/SrtA-RR-2, respectively. DNA encoding the RenG C-terminal domain was PCR amplified from the genomic DNA of strain CTD-MecA with the primer pair RenG156-314-F/linker15-R. These three PCR amplicons were mixed and assembled via OE-PCR using the primer pair SrtA-LF-2/SrtA-RR-2. The resulting OE-PCR amplicon was transformed into the strains MecA-NTD + IFDC2-SrtA and NTD-MecA + IFDC2-SrtA, and selected on THYE medium containing 4-CP to remove the IFDC2 cassette and obtain the strains MecA-NTD + CTD-SrtA and NTD-MecA + CTD-SrtA.

**Construction of strains containing intact luciferase fusions**

The partial ORF and the downstream sequence of *rnjA* were PCR amplified with the primer pairs J1-LF/J1-linker-R and (erm)J1 dn-F/J1-RR, respectively. A PCR amplicon containing the *renG* ORF and erythromycin resistance cassette *ermB* was amplified from the genomic DNA of strain RMRenG (Qin et al, 2021) with the primer pair linker15-F/erm-R. These three PCR amplicons were mixed and assembled via OE-PCR using the primer pair J1-LF/J1-RR. The resulting OE-PCR amplicon was transformed into the strain UA159 and selected for erythromycin resistance to obtain the strain J1-Luc.

The partial ORF and the downstream sequence of *rnjB* were PCR amplified with the primer pairs J2-LF/J2-linker-R and (erm)J2 dn-F/J2-RR, respectively. A PCR amplicon containing the *renG* ORF and erythromycin resistance cassette *ermB* was PCR amplified from the genomic DNA of strain RMRenG with the primer pair linker15-F/erm-R. These three PCR amplicons were mixed and assembled via OE-PCR using the primer pair J2-LF/J2-RR. The resulting OE-PCR amplicon was transformed into the strain UA159 and selected for erythromycin resistance to obtain the strain J2-Luc.

The upstream and downstream sequences of *mecA* were PCR amplified with the primer pairs MecA-LF-1/MecA(linker)-R and (RenG)MecA dn-F/MecA-RR-1, respectively. The *renG* ORF was PCR amplified from the genomic DNA of strain ldhRenGSm with the primer pair (linker)RenG-F/RenG-R. These three PCR amplicons were mixed and assembled via OE-PCR using the primer pair MecA-LF-1/MecA-RR-1. The resulting OE-PCR amplicon was transformed into strain MecAIFDC2 and selected on THYE medium containing 4-CP to remove the IFDC2 cassette and obtain the strain MecA-Luc.

The upstream sequence and the partial ORF of *clpC* were PCR amplified with the primer pairs ClpC-LF-1/ClpC up(erm)-R and (linker)ClpC-F/ClpC-RR-1, respectively. An erythromycin resistance cassette *ermB* was PCR amplified from the plasmid pJY4164 with the primer pair erm-F/erm(rbs)-R-2, while the *renG* ORF was amplified from the genomic DNA of strain ldhRenGSm with the primer pair (rbs)RenG-F/RenG156-314(linker)-R. These four PCR amplicons were mixed and assembled via OE-PCR using the primer pair ClpC-LF-1/ClpC-RR-1. The resulting OE-PCR amplicon was transformed into strain UA159 and selected for erythromycin resistance to obtain the strain Luc-ClpC.

The upstream and downstream flanking sequences of *comX* were PCR amplified from the genomic DNA of ComX-NLuc with the primer pairs ComX-LF/linker15-R and (RenG)ComX dn-F/ComX-RR, respectively. The *renG* ORF was PCR amplified from the genomic DNA of strain ldhRenGSm with the primer pair (linker)RenG-F/RenG-R. These three PCR amplicons were mixed and assembled via OE-PCR using the primer pair ComX-LF/ComX-RR. The resulting OE-PCR amplicon was transformed into the strain ComX-IFDC2/ΔM and selected on THYE medium containing 4-CP to remove the IFDC2 cassette and obtain the strain ComX-Luc.

The partial ORF and the downstream sequence of *thiI* were PCR amplified with the primer pairs ThiI-LF-1/ThiI(linker)-R and (kan)ThiI dn-F/ThiI-RR-1, respectively. The *renG* ORF was PCR amplified from the genomic DNA of strain ldhRenGSm with the primer pair (linker)RenG-F/RenG(kan)-R. A kanamycin resistance cassette *aphAIII* was amplified from the plasmid pWVKTs with the primer pair Kan-F/Kan-R. These four PCR amplicons were mixed and assembled via OE-PCR using the primer pair ThiI-LF-1/ThiI-RR-1. The resulting OE-PCR amplicon was transformed into strain UA159 and selected for kanamycin resistance to obtain the strain ThiI-Luc.

The partial ORF and the downstream sequence of *gltB* were PCR amplified with the primer pairs GltB-LF-2/GltB(linker)-R and (kan)GltB dn-F/GltB-RR-2, respectively. The *renG* ORF was PCR amplified from the genomic DNA of strain ldhRenGSm with the primer pair (linker)RenG-F/RenG(kan)-R. A kanamycin resistance cassette *aphAIII* was amplified from the plasmid pWVKTs with the primer pair Kan-F/Kan-R. These four PCR amplicons were mixed and assembled via

OE-PCR using the primer pair GltB-LF-2/GltB-RR-2. The resulting OE-PCR amplicon was transformed into strain UA159 and selected for kanamycin resistance to obtain the strain GltB-Luc.

The partial ORF and the downstream sequence of *alaS* were PCR amplified with the primer pairs AlaS-LF/AlaS(linker)-R-2 and (kan)AlaS dn-F/AlaS-RR, respectively. The *renG* ORF was PCR amplified from the genomic DNA of strain ldhRenGSm with the primer pair (linker)RenG-F/RenG(kan)-R. A kanamycin resistance cassette *aphAIII* was amplified from the plasmid pWVKTs with the primer pair (RenG)kan-F-2/Kan-R-2. These four PCR amplicons were mixed and assembled via OE-PCR using the primer pair AlaS-LF/AlaS-RR. The resulting OE-PCR amplicon was transformed into strain UA159 and selected for kanamycin resistance to obtain the strain AlaS-Luc.

The upstream and the partial ORF of *dnaG* were PCR amplified with the primer pairs DnaG-LF-2/DnaG up(RenG)-R and (linker)DnaG-F/DnaG-RR-2, respectively. The *renG* ORF was PCR amplified from the genomic DNA of strain ldhRenGSm with the primer pair (DnaG up)RenG-F/RenG156-314(linker)-R. These three amplicons were mixed and assembled by OE-PCR using the primer pair DnaG-LF-2/-DnaG-RR-2. The resulting OE-PCR amplicon was transformed into the strain IFDC2-DnaG and selected on THYE medium containing 4-CP to remove the IFDC2 cassette and obtain the strain Luc-DnaG.

The partial ORF and the downstream sequence of *gyrA* were PCR amplified with the primer pairs GyrA-LF/GyrA(linker)-R-2 and (kan)GyrA dn-F/GyrA-RR, respectively. The *renG* ORF was PCR amplified from the genomic DNA of strain ldhRenGSm with the primer pair (linker)RenG-F/RenG(kan)-R. A kanamycin resistance cassette *aphAIII* was amplified from the plasmid pWVKTs with the primer pair (RenG)kan-F-2/Kan-R-2. These four PCR amplicons were mixed and assembled via OE-PCR using the primer pair GyrA-LF/GyrA-RR. The resulting OE-PCR amplicon was transformed into strain UA159 and selected for kanamycin resistance to obtain the strain GyrA-Luc.

The partial ORF and the downstream sequence of *ilvD* were PCR amplified with the primer pairs IlvD-LF/IlvD(linker)-R-2 and (kan)IlvD dn-F/IlvD-RR, respectively. The *renG* ORF was PCR amplified from the genomic DNA of strain ldhRenGSm with the primer pair (linker)RenG-F/RenG(kan)-R. A kanamycin resistance cassette *aphAIII* was amplified from the plasmid pWVKTs with the primer pair (RenG)kan-F-2/Kan-R-2. These four PCR amplicons were mixed and assembled via OE-PCR using the primer pair IlvD-LF/IlvD-RR. The resulting OE-PCR amplicon was transformed into strain UA159 and selected for kanamycin resistance to obtain the strain IlvD-Luc.

The partial ORF and the downstream sequence of *mnmG* were PCR amplified with the primer pairs MnmG-LF/MnmG(linker)-R-2 and (kan)MnmG dn-F/MnmG-RR, respectively. The *renG* ORF was PCR amplified from the genomic DNA of strain ldhRenGSm with the primer pair (linker)RenG-F/RenG(kan)-R. A kanamycin resistance cassette *aphAIII* was amplified from the plasmid pWVKTs with the primer pair (RenG)kan-F-2/Kan-R-2. These four PCR amplicons were mixed and assembled via OE-PCR using the primer pair MnmG-LF/MnmG-RR. The resulting OE-PCR amplicon was transformed into strain UA159 and selected for kanamycin resistance to obtain the strain MnmG-Luc.

The partial ORF and the downstream sequence of *pcrA* were PCR amplified with the primer pairs PcrA-LF/PcrA(linker)-R-2 and (kan)PcrA dn-F/PcrA-RR, respectively. The *renG* ORF was PCR amplified from the genomic DNA of strain ldhRenGSm with the primer pair (linker)RenG-F/RenG(kan)-R. A kanamycin resistance cassette *aphAIII* was amplified from the plasmid pWVKTs with the primer pair (RenG)kan-F-2/Kan-R-2. These four PCR amplicons were mixed and assembled via OE-PCR using the primer pair PcrA-LF/PcrA-RR. The resulting OE-PCR amplicon was transformed into strain UA159 and selected for kanamycin resistance to obtain the strain PcrA-Luc.

The partial ORF and the downstream sequence of *pheT* were PCR amplified with the primer pairs PheT-LF/PheT(linker)-R-2 and (kan)PheT dn-F/PheT-RR, respectively. The *renG* ORF was PCR amplified from the genomic DNA of strain ldhRenGSm with the primer pair (linker)RenG-F/RenG(kan)-R. A kanamycin resistance cassette *aphAIII* was amplified from the plasmid pWVKTs with the primer pair (RenG)kan-F-2/Kan-R-2. These four PCR amplicons were mixed and assembled via OE-PCR using the primer pair PheT-LF/PheT-RR. The resulting OE-PCR amplicon was transformed into strain UA159 and selected for kanamycin resistance to obtain the strain PheT-Luc.

The partial ORF and the downstream sequence of *spaP* were PCR amplified with the primer pairs SpaP-LF/SpaP(linker)-R-2 and (kan)SpaP dn-F/SpaP-RR, respectively. The *renG* ORF was PCR amplified from the genomic DNA of strain ldhRenGSm with the primer pair (linker)RenG-F/RenG(kan)-R. A kanamycin resistance cassette *aphAIII* was amplified from the plasmid pWVKTs with the primer pair (RenG)kan-F-2/Kan-R-2. These four PCR amplicons were mixed and assembled via OE-PCR using the primer pair SpaP-LF/SpaP-RR. The resulting OE-PCR amplicon was transformed into strain UA159 and selected for kanamycin resistance to obtain the strain SpaP-Luc.

The partial ORF and the downstream sequence of *topA* were PCR amplified with the primer pairs TopA-LF/TopA(linker)-R-2 and (kan)TopA dn-F/TopA-RR, respectively. The *renG* ORF was PCR amplified from the genomic DNA of strain ldhRenGSm with the primer pair (linker)RenG-F/RenG(kan)-R. A kanamycin resistance cassette *aphAIII* was amplified from the plasmid pWVKTs with the primer pair (RenG)kan-F-2/Kan-R-2. These four PCR amplicons were mixed and assembled via OE-PCR using the primer pair TopA-LF/TopA-RR. The resulting OE-PCR amplicon was transformed into strain UA159 and selected for kanamycin resistance to obtain the strain TopA-Luc.

The partial ORF and the downstream sequence of *uvrA* were PCR amplified with the primer pairs UvrA-LF/UvrA(linker)-R-2 and (kan)UvrA dn-F/UvrA-RR, respectively. The *renG* ORF was PCR amplified from the genomic DNA of strain ldhRenGSm with the primer pair (linker)RenG-F/RenG(kan)-R. A kanamycin resistance cassette *aphAIII* was amplified from the plasmid pWVKTs with the primer pair (RenG)kan-F-2/Kan-R-2. These four PCR amplicons were mixed and assembled via OE-PCR using the primer pair UvrA-LF/UvrA-RR. The resulting OE-PCR amplicon was transformed into strain UA159 and selected for kanamycin resistance to obtain the strain UvrA-Luc.

The partial ORF and the downstream sequence of *xseA* were PCR amplified with the primer pairs XseA-LF/XseA(linker)-R-2 and (kan)XseA dn-F/XseA-RR, respectively. The *renG* ORF was PCR amplified from the genomic DNA of strain ldhRenGSm with the primer pair (linker)RenG-F/RenG(kan)-R. A kanamycin resistance cassette *aphAIII* was amplified from the plasmid pWVKTs with the primer pair (RenG)kan-F-2/Kan-R-2. These four PCR amplicons were mixed and assembled via OE-PCR using the primer pair XseA-LF/XseA-RR. The resulting OE-PCR amplicon was transformed into strain UA159 and selected for kanamycin resistance to obtain the strain XseA-Luc.

**Construction of strains for Barstar-Barnase split luciferase complementation and intact luciferase fusions**

DNA encoding the RenG N-terminal domain (NTD, 1-155 amino acid) was translationally fused to the C-terminus of barstar, while the RenG C-terminal domain (CTD, 156-314 amino acid) was translationally fused to the C-terminus of barnase. A flexible linker 3× GGGGS was inserted between the protein of interest and the RenG fragment. Codon optimized barstar and barnase sequences were synthesized. The barstar ORF was then transcriptionally fused to the *ldh* gene (lactate dehydrogenase), while the barnase ORF was transcriptionally fused to the *rpoB* gene (beta subunit of RNA polymerase). The strains were constructed as follows. The *ldh* ORF and downstream sequence were PCR amplified with the primer pairs LDH-F/LDH(rbs-Barstar)-R and (erm)LDH dn-F/LDH-RR, respectively. DNA encoding the barstar ORF was amplified from a synthetic gBlock (Integrated DNA Technologies) with the primer pair (rbs)Barstar-F/linker15-R. DNA encoding the RenG N-terminal domain together with an erythromycin resistance cassette *ermB* was PCR amplified from the genomic DNA of strain MecA-CTD + ClpC-NTD with the primer pair linker15-F/erm-R. These four amplicons were mixed and assembled via OE-PCR using the primer pair LDH-F/LDH-RR. The resulting OE-PCR amplicon was transformed into strain UA159 and selected for kanamycin resistance to obtain the strain Barstar-NTD.

The partial ORF and the downstream sequence of *rpoB* were PCR amplified with the primer pairs RpoB-LF/RpoB(rbs-Barnase)-R and (Kan)RpoB dn-F/RpoB-RR, respectively. DNA encoding the barnase ORF was PCR amplified from a synthetic gBlock (Integrated DNA Technologies) with the primer pair (rbs)Barnase-F/linker15-R. DNA encoding the RenG C-terminal domain together with a kanamycin resistance cassette *aphAIII* was PCR amplified from the genomic DNA of the strain MecA-NTD + AlaS-CTD with the primer pair linker15-F/KanR-R-2. These four amplicons were mixed and assembled via OE-PCR using the primer pair RpoB-LF/RpoB-RR. The resulting OE-PCR amplicon was transformed into the strain Barstar-NTD and selected for kanamycin resistance to obtain the strain Barstar-NTD + Barnase-CTD.

A PCR amplicon encoding the *ldh* and barstar ORFs was generated from the genomic DNA of the strain Barstar-NTD with the primer pair LDH-F/linker15-R. A PCR amplicon encoding the *renG* ORF and an erythromycin resistance cassette *ermB* was PCR amplified from the genomic DNA of strain J1-Luc with the primer pair linker15-F/erm-R. The downstream sequence of *ldh* was PCR amplified with the primer pair (erm)LDH dn-F/LDH-RR. These three amplicons were mixed and assembled via OE-PCR using the primer pair LDH-F/LDH-RR. The resulting OE-PCR amplicon was transformed into the strain UA159 and selected for erythromycin resistance to obtain the strain Barstar-Luc.

A PCR amplicon containing the partial *rpoB* ORF and the barnase ORF was generated from the genomic DNA of the strain Barstar-NTD + Barnase-CTD with the primer pair RpoB-LF/linker15-R. DNA encoding the *renG* ORF and a kanamycin resistance cassette *aphAIII* was PCR amplified from the genomic DNA of strain ThiI-Luc with the primer pair linker15-F/KanR-R-2. The downstream sequence of *rpoB* was PCR amplified with the primer pair (Kan)RpoB dn-F/RpoB-RR. These three amplicons were mixed and assembled via OE-PCR using the primer pair RpoB-LF/RpoB-RR. The resulting OE-PCR amplicon was transformed into the strain UA159 and selected for kanamycin resistance to obtain the strain Barnase-Luc.

**Construction of strains for coimmunoprecipitation analysis**

The 3× FLAG epitope tag was translationally fused to the C terminus of MecA and the 3× HA epitope tag was translationally fused to either the C- or N-terminus of MecA interacting proteins. Strains were constructed as follows. The upstream and downstream sequences of *mecA* were PCR amplified with the primer pairs MecA-LF-1/MecA(lnker)-R-2 and (erm)MecA dn-F-2/MecA-RR-1, respectively. DNA encoding the 3× FLAG epitope tag was PCR amplified from the plasmid pJYRFG (Qin et al, 2021) with the primer pair C-Flag3-F/C-Flag3-R. An erythromycin resistance cassette *ermB* was amplified from the plasmid pJY4164 with the primer pair (FLAG3)erm-F/erm-R. These four PCR amplicons were mixed and assembled via OE-PCR with the primer pair MecA-LF-1/MecA-RR-1. The resulting OE-PCR amplicon was transformed into strain UA159 and selected for erythromycin resistance to obtain the strain MecAFG.

The upstream region and the partial ORF of *clpC* were PCR amplified with the primer pairs ClpC-LF-1/ClpC up(kan)-R and (linker)ClpC-F/ClpC-RR-1, respectively. A kanamycin resistance cassette *aphAIII* was PCR amplified from the plasmid pWVKTs with the primer pair (ClpC up)kan-F/kan(rbs)-R. DNA encoding the 3× HA epitope tag was PCR amplified from the genomic DNA of strain RdprA$^{OE}$/ΔM (Qin et al, 2021) with the primer pair (kan)HA3-F/HA3(liker)-R. These four PCR amplicons were mixed and assembled via OE-PCR using the primer pair ClpC-LF-1/ClpC-RR-1. The resulting OE-PCR amplicon was transformed into the strain MecAFG and selected for kanamycin resistance to obtain the strain MecAFG + HA-ClpC.

The partial ORF and the downstream sequence of *gyrA* were PCR amplified with the primer pairs GyrA-LF/GyrA(linker)-R and (kan)GyrA dn-F/GyrA-RR, respectively. DNA encoding the 3× HA epitope tag was amplified from the genomic DNA of strain RdprA$^{OE}$/ΔM with the primer pair (linker)C-HA3-F/C-HA3-R. A kanamycin resistance cassette *aphAIII* was amplified from the plasmid pWVKTs with the primer pair (HA3)kan-F-2/KanR-R-2. These four PCR amplicons were mixed and assembled via OE-PCR using the primer pair GyrA-LF/GyrA-RR. The resulting OE-PCR amplicon was transformed into the strain MecAFG and selected for kanamycin resistance to obtain the strain MecAFG + GyrAHA.

The partial ORF and the downstream sequence of *thiI* were PCR amplified with the primer pairs ThiI-LF-1/ThiI(linker)-R-2 and (kan)ThiI dn-F/ThiI-RR-1, respectively. DNA encoding the 3× HA epitope tag was amplified from the genomic DNA of strain RdprA$^{OE}$/ΔM with the primer pair C-HA3-F/C-HA3-R. A kanamycin resistance cassette *aphAIII* was amplified from the plasmid pWVKTs with the primer pair (HA3)kan-F/Kan-R. These four PCR amplicons were mixed and assembled via OE-PCR using the primer pair ThiI-LF-1/ThiI-RR-1. The resulting OE-PCR amplicon was transformed into the strain MecAFG and selected for kanamycin resistance to obtain the strain MecAFG + ThiIHA.

The partial ORF and the downstream sequence of *gltB* were PCR amplified with the primer pairs GltB-LF-2/GltB(linker)-R-2 and (kan)GltB dn-F/GltB-RR-2, respectively. DNA encoding the 3× HA epitope tag containing a kanamycin resistance cassette *aphAIII* fragment was PCR amplified from the genomic DNA of the strain MecAFG + ThiIHA with the primer pair C-HA3-F/Kan-R. These three amplicons were mixed and assembled via OE-PCR using the primer pair GltB-LF-2/GltB-RR-2. The resulting OE-PCR amplicon was transformed into the strain MecAFG and selected for kanamycin resistance to obtain the strain MecAFG + GltBHA.

The upstream sequence and the partial ORF of *dnaG* were PCR amplified with the primer pairs DnaG-LF-2/DnaG up(IFDC2)-R and (IFDC2)DnaG up-F/DnaG-RR-2, respectively. The IFDC2 cassette was amplified from the plasmid pIFDC2 using the primer pair IFDC2-F/IFDC2-R. These three PCR amplicons were mixed and assembled via OE-PCR using the primer pair DnaG-LF-2/DnaG-RR-2. The resulting OE-PCR amplicon was transformed into the strain MecAFG and selected for erythromycin resistance to generate the intermediate strain MecAFG + IFDC2-DnaG. Next, the upstream sequence and the partial ORF of *dnaG* were PCR amplified with the primer pairs DnaG-LF-2/DnaG up(HA3)-R and (linker)DnaG-F/DnaG-RR-2, respectively. DNA encoding the 3× HA epitope tag was amplified from the genomic DNA of strain RdprA$^{OE}$/ΔM with the primer pair N-HA3-F/N-HA3(linker)-R. These three PCR amplicons were mixed and assembled via OE-PCR using the primer pair DnaG-LF-2/DnaG-RR-2. The resulting OE-PCR amplicon was transformed into the strain MecAFG + IFDC2-DnaG and selected on THYE medium containing 4-CP to remove the IFDC2 cassette and obtain the strain MecAFG + HA-DnaG.

The partial ORF and the downstream sequence of *alaS* were PCR amplified with the primer pairs AlaS-LF/AlaS(linker)-R and (kan)AlaS dn-F/AlaS-RR, respectively. DNA encoding the 3× HA epitope tag was amplified from the genomic DNA of strain MecAFG + ThiIHA with the primer pair linker15-F/C-HA3-R. A kanamycin resistance cassette *aphAIII* was amplified from the plasmid pWVKTs with the primer pair (HA3)kan-F-2/KanR-R-2. These four PCR amplicons were mixed and assembled via OE-PCR using the primer pair AlaS-LF/AlaS-RR. The resulting OE-PCR amplicon was transformed into the strain MecAFG and selected for kanamycin resistance to obtain the strain MecAFG + AlaSHA.

The upstream and downstream flanking sequences of *srtA* were PCR amplified with the primer pairs SrtA-LF/SrtA(linker)-R-2 and (kan)SrtA dn-F/SrtA-RR, respectively. DNA encoding the 3× HA epitope tag was amplified from the genomic DNA of strain MecAFG + ThiIHA with the primer pair C-HA3-F/C-HA3-R. A kanamycin resistance cassette *aphAIII* was amplified from the plasmid pWVKTs with the primer pair (HA3)kan-F/Kan-R. These four PCR amplicons were mixed and assembled via OE-PCR using the primer pair SrtA-LF/SrtA-RR. The resulting OE-PCR amplicon was transformed into the strain MecAFG and selected for kanamycin resistance to obtain the strain MecAFG + SrtAHA.

**Construction of strains to measure ClpC, GltB, ThiI, DnaG, AlaS, GyrA, and SrtA protein abundance**

To analyze the expression pattern of selected MecA interacting proteins, DNA encoding a 3×HA epitope tag was added to the 3' of each of the genes, while a FLAG epitope tag was added to the 3' of *ldh* in each of the strains to use as a protein loading control. Strains were constructed as follows. The upstream and downstream sequences of *mecA* were PCR amplified with the primer pair MecA-LF-2/MecA up(erm)-R and (erm)MecA dn-F/MecA-RR-1, respectively. An erythromycin resistance cassette *ermB* was amplified from the plasmid pJY4164 with the primer pair erm-F/erm-R. These three PCR amplicons were mixed and assembled via OE-PCR using the primer pair MecA-LF-2/MecA-RR-1. The resulting OE-PCR amplicon was transformed into strain LDHFG (Qin et al, 2021) and selected for erythromycin resistance to obtain the strain ΔMecA.

A PCR amplicon containing the upstream and the partial ORF of *clpC*, a 3×HA epitope tag, and a kanamycin resistance cassette *aphAIII* was generated from the genomic DNA of the strain MecAFG + HA-ClpC with the primer pair ClpC-LF-1/ClpC-RR-1. The resulting amplicon was transformed into the strains LDHFG and ΔMecA and selected for kanamycin resistance to generate the strains HA-ClpC and HA-ClpC/ ΔMecA. The upstream and downstream flanking sequences of *clpP* were PCR amplified with the primer pairs ClpP-LF/ClpP(Tet)-R and (Tet)ClpP dn-F/ClpP-RR, respectively. A tetracycline resistance cassette was amplified from strain GN03XC (Niu et al, 2010) with the primer pair Tet-F/Tet-R. These three amplicons were mixed and assembled via OE-PCR using the primer pair ClpP-LF/ClpP-RR. The resulting OE-PCR amplicon was transformed into the strain HA-ClpC/ ΔMecA and selected for tetracycline resistance to obtain the strain HA-ClpC/ ΔAP.

A PCR amplicon containing the partial ORF and the downstream sequence of *gltB*, a 3×HA epitope tag, and a kanamycin resistance cassette *aphAIII* was generated from the genomic DNA of the strain MecAFG + GltBHA with the primer pair GltB-LF-2/GltB-RR-2. The resulting amplicon was transformed into the strains LDHFG and ΔMecA and selected for kanamycin resistance to generate the strains GltBHA and GltBHA/ ΔMecA. The upstream and downstream flanking sequences of *clpP* and a tetracycline resistance cassette was amplified from the genomic DNA of the strain HA-ClpC/ ΔAP with the primer pair ClpP-LF/ClpP-RR. The resulting amplicon was transformed into the strain GltBHA/ ΔMecA and selected for tetracycline resistance to obtain the strain GltBHA/ ΔAP.

A similar approach was employed to construct epitope tagged versions of ThiI, DnaG, AlaS, GyrA and SrtA, except the fragment containing the partial ORF and the upstream or downstream sequence of the corresponding gene, a 3×HA epitope tag, and a kanamycin resistance cassette *aphAIII* was amplified from the genomic DNA of the strain MecAFG + ThiIHA with the primer pair ThiI-LF-1/ThiI-RR-1, the strain MecAFG + HA-DnaG with the primer pair DnaG-LF-2/DnaG-RR-2, the strain MecAFG + AlaSHA with the primer pair AlaS-LF/AlaS-RR, the strain MecAFG + GyrAHA with the primer pair GyrA-LF/GyrA-RR, and the strain MecAFG + SrtAHA with the primer pair SrtA-LF/SrtA-RR, respectively. The amplicons were transformed into the strains LDHFG and ΔMecA and selected for kanamycin resistance to generate the strains ThiIHA, ThiIHA/ ΔMecA, HA-DnaG, HA-DnaG/ ΔMecA, AlaSHA, AlaSHA/ ΔMecA, GyrAHA, GyrAHA/ ΔMecA, SrtAHA, and SrtAHA/ ΔMecA. The upstream and downstream flanking sequences of *clpP* and a tetracycline resistance cassette was amplified from the genomic DNA of the strain HA-ClpC/ ΔAP with the primer pair ClpP-LF/ClpP-RR. The resulting amplicon was transformed into the above strains and selected for tetracycline resistance to obtain the strains ThiIHA/ ΔAP, HA-DnaG/ ΔAP, AlaSHA/ ΔAP, GyrAHA/ ΔAP and SrtAHA/ ΔAP.

**Construction of strains to measure SpaP, TopA, IlvD, PcrA, MnmG, PheT, XseA, and UvrA protein abundance**

The partial ORF and the downstream sequence of *spaP* were amplified with the primer pairs SpaP-LF/SpaP(linker)-R and (kan)SpaP dn-F/SpaP-RR, respectively. A 3×HA epitope tag and a kanamycin resistance cassette *aphAIII* was amplified from the genomic DNA of the strain MecAFG + AlaSHA with the primer pair (linker)C-HA3-F/KanR-R-2. These three amplicons were mixed and assembled via OE-PCR using the primer pair SpaP-LF/SpaP-RR. The resulting amplicon was transformed into the strains LDHFG and ΔMecA and selected for kanamycin resistance to generate the strains SpaPHA and SpaPHA/ ΔMecA. The upstream and downstream flanking sequences of *clpP* and a tetracycline resistance cassette was amplified from the genomic DNA of the strain HA-ClpC/ ΔAP with the primer pair ClpP-LF/ClpP-RR. The resulting amplicon was transformed into the strain SpaPHA/ ΔMecA and selected for tetracycline resistance to obtain the strain SpaPHA/ ΔAP.

A similar approach was used for construction of TopA, IlvD, PcrA, MnmG, PheT, XseA and UvrA epitope tagged strains, except the partial ORF and the downstream sequence of *topA* were PCR amplified with the primer pairs TopA-LF/TopA(linker)-R and (kan)TopA dn-F/TopA-RR, the partial ORF and the downstream sequence of *ilvD* were amplified with the primer pairs IlvD-LF/IlvD(linker)-R and (kan)IlvD dn-F/IlvD-RR, the partial ORF and the downstream sequence of *pcrA* were amplified with the primer pairs PcrA-LF/PcrA(linker)-R and (kan)PcrA dn-F/PcrA-RR, the partial ORF and the downstream sequence of *mnmG* were amplified with the primer pairs MnmG-LF/MnmG(linker)-R and (kan)MnmG dn-F/MnmG-RR, the partial ORF and the downstream sequence of *pheT* were amplified with the primer pairs PheT-LF/PheT(linker)-R and (kan)PheT dn-F/PheT-RR, the partial ORF and the downstream sequence of *xseA* were amplified with the primer pairs XseA-LF/XseA(linker)-R and (kan)XseA dn-F/XseA-RR, the partial ORF and the downstream sequence of *uvrA* were amplified with the primer pairs UvrA-LF/UvrA(linker)-R and (kan)UvrA dn-F/UvrA-RR. A 3×HA epitope tag and a kanamycin resistance cassette *aphAIII* was amplified from the genomic DNA of the strain MecAFG + AlaSHA with the primer pair (linker)C-HA3-F/KanR-R-2. The three corresponding amplicons for each construct were assembled via OE-PCR, transformed into the strains LDHFG and ΔMecA, and then selected for kanamycin resistance to generate the strains TopAHA, TopAHA/ ΔMecA, IlvDHA, IlvDHA/ ΔMecA, PcrAHA, PcrAHA/ ΔMecA, MnmGHA, MnmGHA/ ΔMecA, PheTHA, PheTHA/ ΔMecA, XseAHA, XseAHA/ ΔMecA, UvrAHA and UvrAHA/ ΔMecA. The upstream and downstream flanking sequences of *clpP* and a tetracycline resistance cassette was amplified from the genomic DNA of the strain HA-ClpC/ ΔAP with the primer pair ClpP-LF/ClpP-RR. The resulting OE-PCR amplicon was transformed into the strains TopAHA/ ΔMecA, IlvDHA/ ΔMecA, PcrAHA/ ΔMecA, MnmGHA/ ΔMecA, PheTHA/ ΔMecA, XseAHA/ ΔMecA and UvrAHA/ ΔMecA, and selected for tetracycline resistance to obtain the strains TopAHA/ ΔAP, IlvDHA/ ΔAP, PcrAHA/ ΔAP, MnmGHA/ ΔAP, PheTHA/ ΔAP, XseAHA/ ΔAP and UvrAHA/ ΔAP.

### Supplemental Tables

**Table S1. Strains and plasmids used in this study**

| Strain | Relevant characteristics | Reference |
|---|---|---|
| UA159 | *S. mutans* genome reference strain | (Ajdić et al, 2002) |
| MecATAP | UA159:: (*mecA*-TAP), $Em^R$ | This study |
| J1IFDC2 | UA159:: (*rnaseJ1*-IFDC2), $Em^R$ | This study |
| ldhRenGSm | UA159:: (*ldh*$_{rbs}$-optimized green renilla luc), 4-$CP^S$ | Merritt et al, 2016 |
| J1-NTD1 | UA159:: (*rnaseJ1*-$RenG_{1\text{-}155}$), 4-$CP^S$ | This study |
| J1-NTD2 | UA159:: (*rnaseJ1*-$RenG_{1\text{-}299}$), 4-$CP^S$ | This study |
| J1NTD1-J2IFDC2 | UA159:: (*rnaseJ1*-$RenG_{1\text{-}155}$, *rnaseJ2*-IFDC2), 4-$CP^S$, $Em^R$ | This study |
| J1NTD2-J2IFDC2 | UA159:: (*rnaseJ1*-$RenG_{1\text{-}299}$, *rnaseJ2*-IFDC2), 4-$CP^S$, $Em^R$ | This study |
| J1-NTD1+J2-CTD1 | UA159:: (*rnaseJ1*-$RenG_{1\text{-}155}$, *rnaseJ2*-$RenG_{156\text{-}314}$), 4-$CP^S$ | This study |
| J1-NTD2+J2-CTD2 | UA159:: (*rnaseJ1*-$RenG_{1\text{-}299}$, *rnaseJ2*-$RenG_{230\text{-}314}$), 4-$CP^S$ | This study |
| NTD1 | UA159:: (*rnaseJ1*p-$RenG_{1\text{-}155}$), $Em^R$ | This study |
| NTD2 | UA159:: (*rnaseJ1*p-$RenG_{1\text{-}229}$), $Em^R$ | This study |
| NTD1 + CTD1 | UA159:: (*rnaseJ1*p-$RenG_{1\text{-}155}$, *rnaseJ2*p-$RenG_{156\text{-}314}$), $Em^R$, $Km^R$ | This study |
| NTD2 + CTD2 | UA159:: (*rnaseJ1*p-$RenG_{1\text{-}229}$, *rnaseJ2*p-$RenG_{230\text{-}314}$), $Em^R$, $Km^R$ | This study |
| MecAIFDC2 | UA159:: (*mecA*-IFDC2), $Em^R$ | This study |
| ΔM | UA159 Δ*brsM*, 4-$CP^S$ | Qin et al, 2021 |
| MecAIFDC2/ΔM | UA159 Δ*brsM*:: (*mecA*-IFDC2), 4-$CP^S$, $Em^R$ | This study |
| MecA-NTD | UA159:: (*mecA*-$RenG_{1\text{-}155}$), 4-$CP^S$ | This study |
| MecA-CTD | UA159:: (*mecA*-$RenG_{156\text{-}314}$), 4-$CP^S$ | This study |
| MecA-NTD/ΔM | UA159 Δ*brsM*:: (*mecA*-$RenG_{1\text{-}155}$), 4-$CP^S$ | This study |
| MecA-CTD/ΔM | UA159 Δ*brsM*:: (*mecA*-$RenG_{156\text{-}314}$), 4-$CP^S$ | This study |
| IFDC2-MecA | UA159:: (IFDC2-*mecA*), $Em^R$ | This study |
| IFDC2-MecA/ΔM | UA159 Δ*brsM*:: (IFDC2-*mecA*), 4-$CP^S$, $Em^R$ | This study |
| NTD-MecA | UA159:: ( $RenG_{1\text{-}155}$-*mecA*), 4-$CP^S$ | This study |
| CTD-MecA | UA159:: ( $RenG_{156\text{-}314}$-*mecA*), 4-$CP^S$ | This study |
| NTD-MecA/ΔM | UA159 Δ*brsM*:: ( $RenG_{1\text{-}155}$-*mecA*), 4-$CP^S$ | This study |
| CTD-MecA/ΔM | UA159 Δ*brsM*:: ( $RenG_{156\text{-}314}$-*mecA*), 4-$CP^S$ | This study |
| MecA-CTD + ClpC-NTD | UA159:: (*mecA*-$RenG_{156\text{-}314}$, *clpC*-$RenG_{1\text{-}155}$), 4-$CP^S$, $Em^R$ | This study |
| CTD-MecA + ClpC-NTD | UA159:: ( $RenG_{156\text{-}314}$-*mecA*, *clpC*-$RenG_{1\text{-}155}$), 4-$CP^S$, $Em^R$ | This study |
| MecA-NTD + ClpC-CTD | UA159:: (*mecA*-$RenG_{1\text{-}155}$, *clpC*-$RenG_{156\text{-}314}$), 4-$CP^S$, $Em^R$ | This study |
| NTD-MecA + ClpC-CTD | UA159:: ( $RenG_{1\text{-}155}$-*mecA*, *clpC*-$RenG_{156\text{-}314}$), 4-$CP^S$, $Em^R$ | This study |
| MecA-CTD + | UA159:: (*mecA*-$RenG_{156\text{-}314}$, $RenG_{1\text{-}155}$-*clpC*), 4-$CP^S$, $Em^R$ | This study |

| | | |
|---|---|---|
| NTD-ClpC | | |
| CTD-MecA + NTD-ClpC | UA159:: ( $RenG_{156-314}$-*mecA*, $RenG_{1-155}$-*clpC*), 4-$CP^S$, $Em^R$ | This study |
| MecA-NTD + CTD-ClpC | UA159:: (*mecA*-$RenG_{1-155}$, $RenG_{156-314}$-*clpC*), 4-$CP^S$, $Em^R$ | This study |
| NTD-MecA + CTD-ClpC | UA159:: ( $RenG_{1-155}$-*mecA*, $RenG_{156-314}$-*clpC*), 4-$CP^S$, $Em^R$ | This study |
| MecA-NTD + ComX-IFDC2/ΔM | UA159 Δ*brsM*:: (*mecA*-$RenG_{1-155}$, *comX*-IFDC2), 4-$CP^S$, $Em^R$ | This study |
| MecA-CTD + ComX-IFDC2/ΔM | UA159 Δ*brsM*:: (*mecA*-$RenG_{156-314}$, *comX*-IFDC2), 4-$CP^S$, $Em^R$ | This study |
| NTD-MecA + ComX-IFDC2/ΔM | UA159 Δ*brsM*:: ($RenG_{1-155}$-*mecA*, *comX*-IFDC2), 4-$CP^S$, $Em^R$ | This study |
| CTD-MecA + ComX-IFDC2/ΔM | UA159 Δ*brsM*:: ($RenG_{156-314}$-*mecA*, *comX*-IFDC2), 4-$CP^S$, $Em^R$ | This study |
| MecA-CTD + ComX-NTD | UA159 Δ*brsM*:: (*mecA*-$RenG_{156-314}$, *comX*-$RenG_{1-155}$), 4-$CP^S$ | This study |
| CTD-MecA + ComX-NTD | UA159 Δ*brsM*:: ($RenG_{156-314}$-*mecA*, *comX*-$RenG_{1-155}$), 4-$CP^S$ | This study |
| MecA-NTD + ComX-CTD | UA159 Δ*brsM*:: (*mecA*-$RenG_{1-155}$, *comX*-$RenG_{156-314}$), 4-$CP^S$ | This study |
| NTD-MecA + ComX-CTD | UA159 Δ*brsM*:: ($RenG_{1-155}$-*mecA*, *comX*-$RenG_{156-314}$), 4-$CP^S$ | This study |
| MecA-NTD + IFDC2-ComX /ΔM | UA159 Δ*brsM*:: (*mecA*-$RenG_{1-155}$, IFDC2-*comX*), 4-$CP^S$, $Em^R$ | This study |
| MecA-CTD + IFDC2-ComX /ΔM | UA159 Δ*brsM*:: (*mecA*-$RenG_{156-314}$, IFDC2-*comX*), 4-$CP^S$, $Em^R$ | This study |
| NTD-MecA + IFDC2-ComX /ΔM | UA159 Δ*brsM*:: ($RenG_{1-155}$-*mecA*, IFDC2-*comX*), 4-$CP^S$, $Em^R$ | This study |
| CTD-MecA + IFDC2-ComX /ΔM | UA159 Δ*brsM*:: ($RenG_{156-314}$-*mecA*, IFDC2-*comX*), 4-$CP^S$, $Em^R$ | This study |
| MecA-CTD + NTD-ComX | UA159 Δ*brsM*:: (*mecA*-$RenG_{156-314}$, $RenG_{1-155}$-*comX*), 4-$CP^S$ | This study |
| CTD-MecA + NTD-ComX | UA159 Δ*brsM*:: ($RenG_{156-314}$-*mecA*, $RenG_{1-155}$-*comX*), 4-$CP^S$ | This study |
| MecA-NTD + CTD-ComX | UA159 Δ*brsM*:: (*mecA*-$RenG_{1-155}$, $RenG_{156-314}$-*comX*), 4-$CP^S$ | This study |
| NTD-MecA + CTD-ComX | UA159 Δ*brsM*:: ($RenG_{1-155}$-*mecA*, $RenG_{156-314}$-*comX*), 4-$CP^S$ | This study |
| MecA-NTD + ThiI-CTD | UA159:: (*mecA*-$RenG_{1-155}$, *thiI*-$RenG_{156-314}$), 4-$CP^S$, $Em^R$ | This study |
| MecA-NTD + CTD-ThiI | UA159:: (*mecA*-$RenG_{1-155}$, $RenG_{156-314}$-*thiI*), 4-$CP^S$, $Em^R$ | This study |

| | | | |
|---|---|---|---|
| NTD-MecA + ThiI-CTD | UA159:: | (RenG$_{1-155}$-*mecA*, *thiI*-RenG$_{156-314}$), 4-CP$^S$, Em$^R$ | This study |
| NTD-MecA + CTD-ThiI | UA159:: | (RenG$_{1-155}$-*mecA*, RenG$_{156-314}$-*thiI*), 4-CP$^S$, Em$^R$ | This study |
| MecA-NTD + GltB-IFDC2 | UA159:: | (*mecA*-RenG$_{1-155}$, *gltB*-IFDC2), 4-CP$^S$, Em$^R$ | This study |
| NTD-MecA + GltB-IFDC2 | UA159:: | (RenG$_{1-155}$-*mecA*, *gltB*-IFDC2), 4-CP$^S$, Em$^R$ | This study |
| MecA-NTD + GltB-CTD | UA159:: | (*mecA*-RenG$_{1-155}$, *gltB*-RenG$_{156-314}$), 4-CP$^S$ | This study |
| NTD-MecA + GltB-CTD | UA159:: | (RenG$_{1-155}$-*mecA*, *gltB*-RenG$_{156-314}$), 4-CP$^S$ | This study |
| MecA-NTD + IFDC2-GltB | UA159:: | (*mecA*-RenG$_{1-155}$, IFDC2-*gltB*), 4-CP$^S$, Em$^R$ | This study |
| NTD-MecA + IFDC2-GltB | UA159:: | (RenG$_{1-155}$-*mecA*, IFDC2-*gltB*), 4-CP$^S$, Em$^R$ | This study |
| MecA-NTD + CTD-GltB | UA159:: | (*mecA*-RenG$_{1-155}$, RenG$_{156-314}$-*gltB*), 4-CP$^S$ | This study |
| NTD-MecA + CTD-GltB | UA159:: | (RenG$_{1-155}$-*mecA*, RenG$_{156-314}$-*gltB*), 4-CP$^S$ | This study |
| MecA-NTD + AlaS-CTD | UA159:: | (*mecA*-RenG$_{1-155}$, *alaS*-RenG$_{156-314}$), 4-CP$^S$, Km$^R$ | This study |
| MecA-NTD + IFDC2-AlaS | UA159:: | (*mecA*-RenG$_{1-155}$, IFDC2-*alaS*), 4-CP$^S$, Em$^R$ | This study |
| MecA-NTD + CTD-AlaS | UA159:: | (*mecA*-RenG$_{1-155}$, RenG$_{156-314}$-*alaS*), 4-CP$^S$ | This study |
| MecA-NTD + DnaG-CTD | UA159:: | (*mecA*-RenG$_{1-155}$, *dnaG*-RenG$_{156-314}$), 4-CP$^S$, Km$^R$ | This study |
| IFDC2-DnaG | UA159:: | (IFDC2-*dnaG*), Em$^R$ | This study |
| MecA-NTD + IFDC2-DnaG | UA159:: | (*mecA*-RenG$_{1-155}$, IFDC2-*dnaG*), 4-CP$^S$, Em$^R$ | This study |
| MecA-NTD + CTD-DnaG | UA159:: | (*mecA*-RenG$_{1-155}$, RenG$_{156-314}$-*dnaG*), 4-CP$^S$ | This study |
| MecA-NTD + GyrA-CTD | UA159:: | (*mecA*-RenG$_{1-155}$, *gyrA*-RenG$_{156-314}$), 4-CP$^S$, Km$^R$ | This study |
| MecA-NTD + IlvD-CTD | UA159:: | (*mecA*-RenG$_{1-155}$, *ilvD*-RenG$_{156-314}$), 4-CP$^S$, Km$^R$ | This study |
| MecA-NTD + IFDC2-IlvD | UA159:: | (*mecA*-RenG$_{1-155}$, IFDC2-*ilvD*), 4-CP$^S$, Em$^R$ | This study |
| MecA-NTD + CTD-IlvD | UA159:: | (*mecA*-RenG$_{1-155}$, RenG$_{156-314}$-*ilvD*), 4-CP$^S$ | This study |
| MecA-NTD + MnmG-CTD | UA159:: | (*mecA*-RenG$_{1-155}$, *mnmG*-RenG$_{156-314}$), 4-CP$^S$, Km$^R$ | This study |
| MecA-NTD + IFDC2-MnmG | UA159:: | (*mecA*-RenG$_{1-155}$, IFDC2-*mnmG*), 4-CP$^S$, Em$^R$ | This study |
| MecA-NTD + CTD-MnmG | UA159:: | (*mecA*-RenG$_{1-155}$, RenG$_{156-314}$-*mnmG*), 4-CP$^S$ | This study |
| MecA-NTD + PcrA-CTD | UA159:: | (*mecA*-RenG$_{1-155}$, *pcrA*-RenG$_{156-314}$), 4-CP$^S$, Km$^R$ | This study |
| MecA-NTD + IFDC2-PcrA | UA159:: | (*mecA*-RenG$_{1-155}$, IFDC2-*pcrA*), 4-CP$^S$, Em$^R$ | This study |
| MecA-NTD + CTD-PcrA | UA159:: | (*mecA*-RenG$_{1-155}$, RenG$_{156-314}$-*pcrA*), 4-CP$^S$ | This study |

| | | | |
|---|---|---|---|
| MecA-NTD + PheT-CTD | UA159:: | (*mecA*-RenG$_{1\text{-}155}$, *pheT*-RenG$_{156\text{-}314}$), 4-CP$^S$, Km$^R$ | This study |
| MecA-NTD + SpaP-CTD | UA159:: | (*mecA*-RenG$_{1\text{-}155}$, *spaP*-RenG$_{156\text{-}314}$), 4-CP$^S$, Km$^R$ | This study |
| MecA-NTD + IFDC2-SpaP | UA159:: | (*mecA*-RenG$_{1\text{-}155}$, IFDC2-*spaP*), 4-CP$^S$, Em$^R$ | This study |
| MecA-NTD + CTD-SpaP | UA159:: | (*mecA*-RenG$_{1\text{-}155}$, RenG$_{156\text{-}314}$-*spaP*), 4-CP$^S$ | This study |
| MecA-NTD + TopA-CTD | UA159:: | (*mecA*-RenG$_{1\text{-}155}$, *topA*-RenG$_{156\text{-}314}$), 4-CP$^S$, Km$^R$ | This study |
| MecA-NTD + IFDC2-TopA | UA159:: | (*mecA*-RenG$_{1\text{-}155}$, IFDC2-*topA*), 4-CP$^S$, Em$^R$ | This study |
| MecA-NTD + CTD-TopA | UA159:: | (*mecA*-RenG$_{1\text{-}155}$, RenG$_{156\text{-}314}$-*topA*), 4-CP$^S$ | This study |
| MecA-NTD + UvrA-CTD | UA159:: | (*mecA*-RenG$_{1\text{-}155}$, *uvrA*-RenG$_{156\text{-}314}$), 4-CP$^S$, Km$^R$ | This study |
| MecA-NTD + IFDC2-UvrA | UA159:: | (*mecA*-RenG$_{1\text{-}155}$, IFDC2-*uvrA*), 4-CP$^S$, Em$^R$ | This study |
| MecA-NTD + CTD-UvrA | UA159:: | (*mecA*-RenG$_{1\text{-}155}$, RenG$_{156\text{-}314}$-*uvrA*), 4-CP$^S$ | This study |
| MecA-NTD + XseA-CTD | UA159:: | (*mecA*-RenG$_{1\text{-}155}$, *xseA*-RenG$_{156\text{-}314}$), 4-CP$^S$, Km$^R$ | This study |
| MecA-NTD + IFDC2-XseA | UA159:: | (*mecA*-RenG$_{1\text{-}155}$, IFDC2-*xseA*), 4-CP$^S$, Em$^R$ | This study |
| MecA-NTD + CTD-XseA | UA159:: | (*mecA*-RenG$_{1\text{-}155}$, RenG$_{156\text{-}314}$-*xseA*), 4-CP$^S$ | This study |
| MecA-NTD + SrtA-IFDC2 | UA159:: | (*mecA*-RenG$_{1\text{-}155}$, *srtA*-IFDC2), 4-CP$^S$, Em$^R$ | This study |
| NTD-MecA + SrtA-IFDC2 | UA159:: | (RenG$_{1\text{-}155}$-*mecA*, *srtA*-IFDC2), 4-CP$^S$, Em$^R$ | This study |
| MecA-NTD + SrtA-CTD | UA159:: | (*mecA*-RenG$_{1\text{-}155}$, *srtA*-RenG$_{156\text{-}314}$), 4-CP$^S$ | This study |
| NTD-MecA + SrtA-CTD | UA159:: | (RenG$_{1\text{-}155}$-*mecA*, *srtA*-RenG$_{156\text{-}314}$), 4-CP$^S$ | This study |
| MecA-NTD + IFDC2-SrtA | UA159:: | (*mecA*-RenG$_{1\text{-}155}$, IFDC2-*srtA*), 4-CP$^S$, Em$^R$ | This study |
| NTD-MecA + IFDC2-SrtA | UA159:: | (RenG$_{1\text{-}155}$-*mecA*, IFDC2-*srtA*), 4-CP$^S$, Em$^R$ | This study |
| MecA-NTD + CTD-SrtA | UA159:: | (*mecA*-RenG$_{1\text{-}155}$, RenG$_{156\text{-}314}$-*srtA*), 4-CP$^S$ | This study |
| NTD-MecA + CTD-SrtA | UA159:: | (RenG$_{1\text{-}155}$-*mecA*, RenG$_{156\text{-}314}$-*srtA*), 4-CP$^S$ | This study |
| RMRenG | UA159:: | (*brsRM*-RenG), Em$^R$ | Qin et al, 2021 |
| J1-Luc | UA159:: | (*rnase J1*-RenG), Em$^R$ | This study |
| J2-Luc | UA159:: | (*rnase J2*-RenG), Em$^R$ | This study |
| MecA-Luc | UA159:: | (*mecA*-RenG), 4-CP$^S$ | This study |
| Luc-ClpC | UA159:: | (RenG-*clpC*), Em$^R$ | This study |

| | | |
|---|---|---|
| ComX-Luc | UA159 Δ*brsM*:: ( *comX*-RenG), 4-CP$^{S}$ | This study |
| ThiI-Luc | UA159:: (*thiI*-RenG), Km$^{R}$ | This study |
| GltB-Luc | UA159:: (*gltB*-RenG), Km$^{R}$ | This study |
| AlaS-Luc | UA159:: (*alaS*-RenG), Km$^{R}$ | This study |
| Luc-DnaG | UA159:: (RenG-*dnaG*), 4-CP$^{S}$ | This study |
| GyrA-Luc | UA159:: (*gyrA*-RenG), Km$^{R}$ | This study |
| IlvD-Luc | UA159:: (*ilvD*-RenG), Km$^{R}$ | This study |
| MnmG-Luc | UA159:: (*mnmG*-RenG), Km$^{R}$ | This study |
| PcrA-Luc | UA159:: (*pcrA*-RenG), Km$^{R}$ | This study |
| PheT-Luc | UA159:: (*pheT*-RenG), Km$^{R}$ | This study |
| SpaP-Luc | UA159:: (*spaP*-RenG), Km$^{R}$ | This study |
| TopA-Luc | UA159:: (*topA*-RenG), Km$^{R}$ | This study |
| UvrA-Luc | UA159:: (*uvrA*-RenG), Km$^{R}$ | This study |
| XseA-Luc | UA159:: (*xseA*-RenG), Km$^{R}$ | This study |
| Barstar-NTD | UA159:: (*ldh*p-*barstar*-RenG$_{1-155}$), Em$^{R}$ | This study |
| Barstar-NTD + Barnase-CTD | UA159:: (*ldh*p-*barstar*-RenG$_{1-155}$, *rpoB*p-*barnase*-RenG$_{156-314}$), Em$^{R}$, Km$^{R}$ | This study |
| Barstar-Luc | UA159:: (*ldh*p-*barstar*), Em$^{R}$ | This study |
| Barnase-Luc | UA159:: (*rpoB*p-*barnase*-RenG), Km$^{R}$ | This study |
| MecAFG | UA159:: (*mecA*-FLAG3), Em$^{R}$ | This study |
| RdprA$^{OE}$/ΔM | UA159 Δ*brsM*:: (*brsR*-FLAG3, *ldh*-*dprA*-HA3), 4-CP$^{S}$ | Qin et al, 2021 |
| MecAFG + HA-ClpC | UA159:: (*mecA*-FLAG3, HA3-*clpC*), Em$^{R}$, Km$^{R}$ | This study |
| MecAFG + GyrAHA | UA159:: (*mecA*-FLAG3, *gyrA*-HA3), Em$^{R}$, Km$^{R}$ | This study |
| MecAFG + ThiIHA | UA159:: (*mecA*-FLAG3, *thiI*-HA3), Em$^{R}$, Km$^{R}$ | This study |
| MecAFG + GltBHA | UA159:: (*mecA*-FLAG3, *gltB*-HA3), Em$^{R}$, Km$^{R}$ | This study |
| MecAFG + IFDC2-DnaG | UA159:: (*mecA*-FLAG3, IFDC2-*dnaG*), Em$^{R}$, 4-CP$^{S}$ | This study |
| MecAFG + HA-DnaG | UA159:: (*mecA*-FLAG3, HA3-*dnaG*), Em$^{R}$, 4-CP$^{S}$ | This study |
| MecAFG + AlaSHA | UA159:: (*mecA*-FLAG3, *alaS*-HA3), Em$^{R}$, Km$^{R}$ | This study |
| MecAFG + SrtAHA | UA159:: (*mecA*-FLAG3, *srtA*-HA3), Em$^{R}$, Km$^{R}$ | This study |
| LDHFG | UA159:: (*ldh*-FLAG), Spec$^{R}$ | Qin et al, 2021 |
| ΔMecA | UA159 Δ*mecA*:: (*ldh*-FLAG), Em$^{R}$, Spec$^{R}$ | This study |
| HA-ClpC | UA159:: (HA3-*clpC*, *ldh*-FLAG), Km$^{R}$, Spec$^{R}$ | This study |
| HA-ClpC/ ΔMecA | UA159 Δ*mecA*:: (HA3-*clpC*, *ldh*-FLAG), Em$^{R}$, Km$^{R}$, Spec$^{R}$, | This study |
| GN03XC | UA159 Δ*irvR* Δ*clpC* Δ*clpX*, pGN03, Km$^{R}$, Em$^{R}$, Tet$^{R}$, Spec$^{R}$ | Niu et al, 2010 |
| HA-ClpC/ ΔAP | UA159 Δ*mecA* Δ*clpP*:: (HA3-*clpC*, *ldh*-FLAG), Em$^{R}$, Tet$^{R}$, Km$^{R}$, Spec$^{R}$ | This study |
| GltBHA | UA159:: (*gltB*-HA3, *ldh*-FLAG), Km$^{R}$, Spec$^{R}$ | This study |
| GltBHA/ ΔMecA | UA159 Δ*mecA*:: (*gltB*-HA3, *ldh*-FLAG), Em$^{R}$, Km$^{R}$, Spec$^{R}$, | This study |

| | | |
|---|---|---|
| GltBHA/ ΔAP | UA159 Δ*mecA* Δ*clpP*:: (*gltB*-HA3, *ldh*-FLAG), Em$^R$, Tet$^R$, Km$^R$, Spec$^R$ | This study |
| ThiIHA | UA159:: (*thiI*-HA3, *ldh*-FLAG), Km$^R$, Spec$^R$ | This study |
| ThiIHA/ ΔMecA | UA159 Δ*mecA*:: (*thiI*-HA3, *ldh*-FLAG), Em$^R$, Km$^R$, Spec$^R$, | This study |
| ThiIHA/ ΔAP | UA159 Δ*mecA* Δ*clpP*:: (*thiI*-HA3, *ldh*-FLAG), Em$^R$, Tet$^R$, Km$^R$, Spec$^R$ | This study |
| HA-DnaG | UA159:: (HA3-*dnaG*, *ldh*-FLAG), Km$^R$, Spec$^R$ | This study |
| HA-DnaG/ ΔMecA | UA159 Δ*mecA*:: (HA3-*dnaG*, *ldh*-FLAG), Em$^R$, Km$^R$, Spec$^R$, | This study |
| HA-DnaG/ ΔAP | UA159 Δ*mecA* Δ*clpP*:: (HA3-*dnaG*, *ldh*-FLAG), Em$^R$, Tet$^R$, Km$^R$, Spec$^R$ | This study |
| AlaSHA | UA159:: (*alaS*-HA3, *ldh*-FLAG), Km$^R$, Spec$^R$ | This study |
| AlaSHA/ ΔMecA | UA159 Δ*mecA*:: (*alaS*-HA3, *ldh*-FLAG), Em$^R$, Km$^R$, Spec$^R$, | This study |
| AlaSHA/ ΔAP | UA159 Δ*mecA* Δ*clpP*:: (*alaS*-HA3, *ldh*-FLAG), Em$^R$, Tet$^R$, Km$^R$, Spec$^R$ | This study |
| GyrAHA | UA159:: (*gyrA*-HA3, *ldh*-FLAG), Km$^R$, Spec$^R$ | This study |
| GyrAHA/ ΔMecA | UA159 Δ*mecA*:: (*gyrA*-HA3, *ldh*-FLAG), Em$^R$, Km$^R$, Spec$^R$, | This study |
| GyrAHA/ ΔAP | UA159 Δ*mecA* Δ*clpP*:: (*gyrA*-HA3, *ldh*-FLAG), Em$^R$, Tet$^R$, Km$^R$, Spec$^R$ | This study |
| SrtAHA | UA159:: (*srtA*-HA3, *ldh*-FLAG), Km$^R$, Spec$^R$ | This study |
| SrtAHA/ ΔMecA | UA159 Δ*mecA*:: (*srtA*-HA3, *ldh*-FLAG), Em$^R$, Km$^R$, Spec$^R$, | This study |
| SrtAHA/ ΔAP | UA159 Δ*mecA* Δ*clpP*:: (*srtA*-HA3, *ldh*-FLAG), Em$^R$, Tet$^R$, Km$^R$, Spec$^R$ | This study |
| TopAHA | UA159:: (*topA*-HA3, *ldh*-FLAG), Km$^R$, Spec$^R$ | This study |
| TopAHA/ ΔMecA | UA159 Δ*mecA*:: (*topA*-HA3, *ldh*-FLAG), Em$^R$, Km$^R$, Spec$^R$, | This study |
| TopAHA/ ΔAP | UA159 Δ*mecA* Δ*clpP*:: (*topA*-HA3, *ldh*-FLAG), Em$^R$, Tet$^R$, Km$^R$, Spec$^R$ | This study |
| IlvDHA | UA159:: (*ilvD*-HA3, *ldh*-FLAG), Km$^R$, Spec$^R$ | This study |
| IlvDHA/ ΔMecA | UA159 Δ*mecA*:: (*ilvD*-HA3, *ldh*-FLAG), Em$^R$, Km$^R$, Spec$^R$, | This study |
| IlvDHA/ ΔAP | UA159 Δ*mecA* Δ*clpP*:: (*ilvD*-HA3, *ldh*-FLAG), Em$^R$, Tet$^R$, Km$^R$, Spec$^R$ | This study |
| PcrAHA | UA159:: (*pcrA*-HA3, *ldh*-FLAG), Km$^R$, Spec$^R$ | This study |
| PcrAHA/ ΔMecA | UA159 Δ*mecA*:: (*pcrA*-HA3, *ldh*-FLAG), Em$^R$, Km$^R$, Spec$^R$, | This study |
| PcrAHA/ ΔAP | UA159 Δ*mecA* Δ*clpP*:: (*pcrA*-HA3, *ldh*-FLAG), Em$^R$, Tet$^R$, Km$^R$, Spec$^R$ | This study |
| MnmGHA | UA159:: (*mnmG*-HA3, *ldh*-FLAG), Km$^R$, Spec$^R$ | This study |
| MnmGHA/ ΔMecA | UA159 Δ*mecA*:: (*mnmG*-HA3, *ldh*-FLAG), Em$^R$, Km$^R$, Spec$^R$, | This study |
| MnmGHA/ ΔAP | UA159 Δ*mecA* Δ*clpP*:: (*mnmG*-HA3, *ldh*-FLAG), Em$^R$, Tet$^R$, Km$^R$, Spec$^R$ | This study |
| PheTHA | UA159:: (*pheT*-HA3, *ldh*-FLAG), Km$^R$, Spec$^R$ | This study |
| PheTHA/ ΔMecA | UA159 Δ*mecA*:: (*pheT*-HA3, *ldh*-FLAG), Em$^R$, Km$^R$, Spec$^R$, | This study |

| | | |
|---|---|---|
| PheTHA/ ΔAP | UA159 Δ*mecA* Δ*clpP*:: (*pheT*-HA3, *ldh*-FLAG), $Em^R$, $Tet^R$, $Km^R$, $Spec^R$ | This study |
| XseAHA | UA159:: (*xseA*-HA3, *ldh*-FLAG), $Km^R$, $Spec^R$ | This study |
| XseAHA/ ΔMecA | UA159 Δ*mecA*:: (*xseA*-HA3, *ldh*-FLAG), $Em^R$, $Km^R$, $Spec^R$, | This study |
| XseAHA/ ΔAP | UA159 Δ*mecA* Δ*clpP*:: (*xseA*-HA3, *ldh*-FLAG), $Em^R$, $Tet^R$, $Km^R$, $Spec^R$ | This study |
| UvrAHA | UA159:: (*uvrA*-HA3, *ldh*-FLAG), $Km^R$, $Spec^R$ | This study |
| UvrAHA/ ΔMecA | UA159 Δ*mecA*:: (*uvrA*-HA3, *ldh*-FLAG), $Em^R$, $Km^R$, $Spec^R$, | This study |
| UvrAHA/ ΔAP | UA159 Δ*mecA* Δ*clpP*:: (*uvrA*-HA3, *ldh*-FLAG), $Em^R$, $Tet^R$, $Km^R$, $Spec^R$ | This study |

| **Plasmid** | **Relevant characteristics** | **Reference** |
|---|---|---|
| pJY4164 | *E. coli*-*Streptococcus* shuttle vector, *Streptococcus* suicide vector, $Em^R$ | Achen et al, 1986 |
| pIFDC2 | pDL278:: (P*ldh*-*mpheS**-*ermB*), $Spec^R$, $Em^R$ | Xie et al, 2011 |
| pWVKTs | Temperature sensitive *E. coli*-*Streptococcus* shuttle vector, $Km^R$ | Gutierrez et al, 1996 |
| pJYRFG | pJY4164:: (P*ldh*-*brsR*-FLAG3), $Em^R$ | Qin et al, 2021 |

**Table S2. Primers used in this study**

| **Primer name** | **Sequence 5′➔3′** |
|---|---|
| MecA-LF | GGAATACCGATTATGTTTGGTGCT |
| MecA(TAP)-R | CTCCGCCACCCGAGCCCCCTCCACCTCCAATCATTTGTAATTC TTGCAGA |
| (erm)MecA dn-F | AGTTATCTATTATTTAACGGGAGGAAATAAGCTAGATGATAC CAATTACCTTAAAATTTG |
| HA3-FLAG-C-F | GGTGGAGGGGGCTCGGGTGGCGGA |
| J25 FLAG-R | TTACTTATCGTCATCGTCCTTGT |
| (TAP)erm-F | CGATTACAAGGACGATGACGATAAGTAAGATATAATGGGAG ATAAGACGGTTCG |
| erm-R | TTATTTCCTCCCGTTAAATAATAGATAACTAT |
| J1-LF | TCCAGAGCCTCTAGGAATTGTCAT |
| J1 (IFDC2)-R | CTATGCTATGAGTGTTATTGTTGCTCGGTTAAGACTTATCTGG TGTCAATATCATAGGA |
| (IFDC2)J1 dn-F | AGATAAATTATTAGGTATACTACTGACAGCTTCCTATGATAT TGACACCAGATAAGTC |
| J1-RR | TTAAGGTTGGACCTGGTTTCAAAG |
| IFDC2-F | CCGAGCAACAATAACACTCATAGC |

| | |
|---|---|
| IFDC2-R | GAAGCTGTCAGTAGTATACCTAATAATTTATCTACA |
| J1-linker-R | TGATCCTCCACCACCAGATCCACCTCCACCAGACTTATCTGGTGTCAATATCATAGGAAT |
| J1 dn-F | TAAATCATAATTGTAAGCAGCTATTATTAAAGAG |
| (linker)RenG-F-2 | GGTGGAGGTGGATCTGGTGGTGGAGGATCAGCTAGTAAAGTTTATGATCCTGAACAACG |
| RenG155(J1 dn)-R | CTTTAATAATAGCTGCTTACAATTATGATTTAACCCATCCAAGATTCAATAACATCA |
| RenG229 (J1 dn)-R | CTTTAATAATAGCTGCTTACAATTATGATTTATCCACCTTTAACTAAAGGAATTTCACG |
| J2-LF | GCACTAAAGAAGGCAATATTGTTTATACAG |
| J2 (IFDC2)-R | TATGCTATGAGTGTTATTGTTGCTCGGTTATCTAACTTCCATAACTACTGGTAAAATAGC |
| (IFDC2) J2 dn-F | AGATAAATTATTAGGTATACTACTGACAGCTTCCAGCTATTTTACCAGTAGTTATGGAAG |
| J2-RR | CTGTCAAATAAAACCAGAAAAACTTACCT |
| J2-linker-R | TCCTCCACCACCAGATCCACCTCCACCTCTAACTTCCATAACTACTGGTAAAATAGCTG |
| (linker) RenG 156-F | GGTGGAGGTGGATCTGGTGGTGGAGGATCATGGCCAGATATTGAAGAAGAATTG |
| RenG 156-314 (J2 dn)-R | TTGGAAACAGTATCAATCAAGCCTCTTTTAGATAGAACGTTGCTCATTCTTCAAAAC |
| J2 dn-F | TAAAAGAGGCTTGATTGATACTGTTTC |
| (linker) RenG 230-F | GGTGGAGGTGGATCTGGTGGTGGAGGATCAAAGCCTGATGTTGTTGCTATTGTTC |
| J1(rbs-RenG1-155)-R | TAGCCATTCTCTAAACATCTCCTTCTTAAGACTTATCTGGTGTCAATATCATAGGAATG |
| (LDH-rbs)RenG-F | GAAGGAGATGTTTAGAGAATGGCTAGTAAAGTTTATGATCCTGAACA |
| RenG1-155(erm)-R-2 | TATATTTTTGTTCATGTAATCACTCCTTCTTAACCCATCCAAGATTCAATAACATCAAC |
| (erm)J1 dn-F | GTTATCTATTATTTAACGGGAGGAAATAAATCATAATTGTAAGCAGCTATTATTAAAGAG |
| erm-F | GAAGGAGTGATTACATGAACAAAAATATAA |
| RenG1-229(erm)-R | TATTTTTGTTCATGTAATCACTCCTTCTTATCCACCTTTAACTAAAGGAATTTCACGAG |
| J2(rbs-RenG156-314)-R | GCCACATTCTCTAAACATCTCCTTCTTATCTAACTTCCATAACTACTGGTAAAATAGCTG |
| (kan)J2 dn-F-2 | TATTATATTTTACTGGATGAATTGTTTTAGTAAAAGAGGCTTGATTGATACTGTTTCCA |

| | |
|---|---|
| (LDH rbs)RenG156-314-F | GAAGGAGATGTTTAGAGAATGTGGCCAGATATTGAAGAAGA ATTG |
| RenG-R | TTAGATAGAACGTTGCTCATTCTTCAA |
| (RenG)kan-F-2 | TTTGAAGAATGAGCAACGTTCTATCTAATGTAGAAAAGAGG AAGGAAATAATAAATGGC |
| KanR-R-2 | CTAAAACAATTCATCCAGTAAAATATAATATTTTATTTTCTCC |
| J2(LDH rbs)-R | TCTCTAAACATCTCCTTCTTATCTAACTTCCATAACTACTGGT AAAATAGCTG |
| (LDH rbs)RenG230-314-F | TAGATAAGAAGGAGATGTTTAGAGAATGAAGCCTGATGTTG TTGCTATT |
| MecA-LF-1 | GAGCAACTATTTTAGGTGGTATTTTGC |
| MecA(IFDC2)-R | TGCTATGAGTGTTATTGTTGCTCGGTTATCCAATCATTTGTAA TTCTTGCAG |
| (IFDC2)MecA dn-F | TAAATTATTAGGTATACTACTGACAGCTTC GCTAGATGATACCAATTACCTTAAAATTTG |
| MecA-RR-1 | CAATAGTCTGAAGAGTGGTGTTCTTTTAG |
| MecA(linker)-R | ACCAGATCCACCTCCACCAGAACCTCCACCTCCTCCAATCAT TTGTAATTCTTGCAGAG |
| (RenG1-155)MecA dn-F | GATGTTATTGAATCTTGGATGGGTTAAGCTAGATGATACCAA TTACCTTAAAATTTG |
| (MecA)linker-F | TTACAAATGATTGGAGGAGGTGGAGGTTCTGGTGGAGGTGG ATCTGGTGGT |
| RenG1-155-R | TTAACCCATCCAAGATTCAATAACATC |
| (RenG)MecA dn-F | TTTGAAGAATGAGCAACGTTCTATCTAAGCTAGATGATACCA ATTACCTTAAAATTTG |
| MecA-LF-2 | GCAGATTTATCAACCTATACCTCAACTG |
| MecA up(IFDC2)-R | TGCTATGAGTGTTATTGTTGCTCGGAGTCTTTACCTCATACGA GTAGTTATTTCATT |
| (IFDC2)MecA-F | GTAGATAAATTATTAGGTATACTACTGACAGCTTCATGGAAA TGAAACAAATCAGCGA |
| MecA-RR-2 | GAATCTTAAAACTATCAAATCGAAAGTCTG |
| MecA up(RenG1-155)-R | TTCAGGATCATAAACTTTACTAGCCATAGTCTTTACCTCATAC GAGTAGTTATTTCATT |
| (linker)MecA-F | GGTGGAGGTGGATCTGGTGGTGGAGGATCAGAAATGAAACA AATCAGCGAAACA |
| RenG-F-2 | ATGGCTAGTAAAGTTTATGATCCTGAACA |
| RenG1-155(linker)-R | TGATCCTCCACCACCAGATCCACCTCCACCACCCATCCAAGA TTCAATAACATCA |
| MecA up (RenG156-314)-R | CCAATTCTTCTTCAATATCTGGCCACATAGTCTTTACCTCATA CGAGTAGTTATTTCATT |
| RenG156-314-F | ATGTGGCCAGATATTGAAGAAGAATTGG |

| | |
|---|---|
| RenG156-314(linker)-R | TGATCCTCCACCACCAGATCCACCTCCACCGATAGAACGTTGCTCATTCTTCAAAA |
| ClpC-LF-2 | AGATCATGCTCTTATCACTAACGAT |
| ClpC(linker)-R | CCACCAGAACCTCCACCTCCTACAATATCAAATTTTAATTTTTCTTTTTGAATACCTATT |
| (erm)clpC dn-F | ATCTATTATTTAACGGGAGGAAATAATTATCTAGTCTATTTTTCATAAATTTGTTAGCTC |
| ClpC-RR-2 | CTTCACCAGTCTTAGCATCCT |
| linker15-F | GGAGGTGGAGGTTCTGGT |
| RenG1-155-R | TTAACCCATCCAAGATTCAATAACATC |
| (RenG1-155)erm-F | TGTTATTGAATCTTGGATGGGTTAAGATATAATGGGAGATAAGACGGTTCG |
| (RenG)erm-F | GAAGAATGAGCAACGTTCTATCTAAGATATAATGGGAGATAAGACGGTTCG |
| ClpC-LF-1 | AAGATATCAAGGCAGAGCTTAAAG |
| ClpC up(erm)-R | TATATTTTTGTTCATGTAATCACTCCTTCTCATAGATGGTATCCTTTTCTATCTAAACG |
| (linker)ClpC-F | AGGTGGATCTGGTGGTGGAGGATCAACCGATTACTCATTAAAAATGCAGG |
| ClpC-RR-1 | CTTCACTTGGTTCTTCAATAGTTACC |
| erm(rbs)-R-1 | ACTAGCCATGCTTCCAACTCCTTCCTTATTTCCTCCCGTTAAATAATAGATAACTAT |
| (rbs)RenG-F | GGAAGGAGTTGGAAGCATGGCTAGTAAAGTTTATGATCCTGAACA |
| erm(rbs)-R-2 | TCTGGCCACATGCTTCCAACTCCTTCCTTATTTCCTCCCGTTAAATAATAGATAACTAT |
| (rbs)RenG156-314-F | GGAAGGAGTTGGAAGCATGTGGCCAGATATTGAAGAAGA |
| ComX-LF | TACTAATCGCTCGAGGACTTATCCA |
| ComX(IFDC2)-R | TGCTATGAGTGTTATTGTTGCTCGGCTATTTTTCCTTAAAATCACTTAATTTTTTACG |
| (IFDC2)ComX dn-F | TAGATAAATTATTAGGTATACTACTGACAGCTTCTTAAAAAGGGAAAGAATGGAACATGT |
| ComX-RR | TTCAAAGTTTTCACAACAGTAAGCTTTAA |
| ComX(linker)-R | TCCACCTCCACCAGAACCTCCACCTCCTTTTTCCTTAAAATCACTTAATTTTTTACG |
| (RenG1-155)ComX dn-F | TGATGTTATTGAATCTTGGATGGGTTAATTAAAAAGGGAAAGAATGGAACATGT |
| (RenG)ComX dn-F | TTGAAGAATGAGCAACGTTCTATCTAATTAAAAAGGGAAAGAATGGAACATGT |
| ComX up(IFDC2)-R | TGCTATGAGTGTTATTGTTGCTCGGCTATTACGATGACCTCCTTTTATAATAAAATTAT |

| | |
|---|---|
| (IFDC2)ComX-F | ATAAATTATTAGGTATACTACTGACAGCTTCATGGAAGAAGATTTTGAAATTGTTTTTA |
| ComX up(RenG1-155)-R | CAGGATCATAAACTTTACTAGCCATCTATTACGATGACCTCCTTTTATAATAAAATTAT |
| (linker)ComX-F | GTGGAGGTGGATCTGGTGGTGGAGGATCAGAAGAAGATTTTGAAATTGTTTTTAATAAGG |
| ComX up (RenG156-314)-R | ATTCTTCTTCAATATCTGGCCACATCTATTACGATGACCTCCTTTTATAATAAAATTAT |
| Thil-LF-1 | TATCTATCTTAATGGAGCAAACTATGTTC |
| Thil(linker)-R | CACCTCCACCAGAACCTCCACCTCCCAATAAATCTTCGATTAAGACATCAACTT |
| (erm)Thil dn-F | CTATTATTTAACGGGAGGAAATAAAAACAAGAAAAATCATATATTAATCATTTTCTTCAT |
| Thil-RR-1 | ATTTTGTAAATTCTCATAGGACTGGTT |
| Thil-LF-2 | GGACAGATGAAGTCTTTTTTACTTCAG |
| Thil up (erm)-R | ATTTTTGTTCATGTAATCACTCCTTCAGTAAAAATACAAGGTCATTGAATCATTAG |
| (linker)Thil-F | AGGTGGATCTGGTGGTGGAGGATCACAGTATTCAGAAATTATGGTGCGT |
| Thil-RR-2 | CTCCACATTCTTAATTTTAGGATTGGT |
| erm(Thil rbs)-R | ACATAGTAAAAATACAAGGTCATTGAATCTTATTTCCTCCCGTTAAATAATAGATAACT |
| (Thil rbs)RenG156-F | AATAAGATTCAATGACCTTGTATTTTTACTATGTGGCCAGATATTGAAGAAGA |
| linker15-R | TGATCCTCCACCACCAGATC |
| GltB-LF-2 | GAAAAGGCCTGTACAGAAGGCTTA |
| GltB(IFDC2)-R | TGCTATGAGTGTTATTGTTGCTCGGCTCTGTAACCATCATTCTGAGGTTTTG |
| (IFDC2)GltB dn-F | ATAAATTATTAGGTATACTACTGACAGCTTCGAGTAGAACAAAAGAGACAGGACTTTCG |
| GltB-RR-2 | CAGAACGTGAACCTATTATCATTCCTAT |
| GltB(linker)-R | CTCCACCACCAGATCCACCTCCACCCTCTGTAACCATCATTCTGAGGTTTTG |
| (RenG)GltB dn-F | TTGAAGAATGAGCAACGTTCTATCTAAGAGTAGAACAAAAGAGACAGGACTTTCG |
| GltB-LF-1 | GCTCTGTTGATGAAATGGTTGGT |
| GltB up(IFDC2)-R | TGCTATGAGTGTTATTGTTGCTCGGCTCATTTCCCTCCTGTTACACTAGC |
| (IFDC2)GltB-F | GTAGATAAATTATTAGGTATACTACTGACAGCTTCGCAGATCCGTTTGGATTTTTAAAG |
| GltB-RR-1 | ATTCTACTGTTGACGTGATATAGTCCGT |

| | |
|---|---|
| GltB up(RenG156-314)-R | CCAATTCTTCTTCAATATCTGGCCACATCTCATTTCCCTCCTG TTACACTA |
| (linker)GltB-F | AGGTGGATCTGGTGGTGGAGGATCAGCAGATCCGTTTGGATT TTTAAAG |
| AlaS-LF-1 | GACACTGTTGCTCGTGTGAC |
| AlaS(linker)-R | CACCTCCACCAGAACCTCCACCTCCCAAATTCTCTGCTACTG CGGC |
| (kan)AlaS dn-F | TTTTACTGGATGAATTGTTTTAGAACTTATCTTTGCAAAAGAA CTATTTTATAACTAATC |
| AlaS-LF-2 | GGAAAGCTTATAAGAAACGTCTGAAAG |
| AlaS up(IFDC2)-R | TGCTATGAGTGTTATTGTTGCTCGGGACAGATACTTGTATCTT AGGCAGACTATCT |
| (IFDC2)AlaS up-F | TTATTAGGTATACTACTGACAGCTTCGTCAGATTCCTAATTTT CTGTTCAATTACAGTC |
| AlaS-RR-2 | GTCTGAAAATATCTTTTCCTTCAAGCGT |
| AlaS up(RenG156-314)-R | CCAATTCTTCTTCAATATCTGGCCACATATTTAGATATAGAGT CCGACTGTAATTGAAC |
| (linker)AlaS-F | AGGTGGATCTGGTGGTGGAGGATCAAAACAATTAACATCAG CTCAAGTCCG |
| DnaG-LF | TGGAAGGCTTTATGGATGTTATTGC |
| DnaG(linker)-R | CACCTCCACCAGAACCTCCACCTCCTTCCATATTTCTTTTTTG AGCGATCAGG |
| (Kan)DnaG dn-F-2 | TATTTTACTGGATGAATTGTTTTAGAGGAAAAATGGTAAATA ATAAGAAAAAAACATCAA |
| DnaG-RR | AATTTCTTCGGGAGTATAGTTTTTATCTGC |
| DnaG-LF-2 | TCATAGCAATACCACAGGTGGC |
| DnaG up(IFDC2)-R | TGCTATGAGTGTTATTGTTGCTCGGTGCCTTTTATTATAACAT TTTAGAAGTTCCTTGT |
| (IFDC2)DnaG up-F | TATTAGGTATACTACTGACAGCTTCATGCTAAAAAGCAATTT TGAGGTGG |
| DnaG-RR-2 | CTTCTGAATTCTCCTTGATAAATTCATCTGG |
| DnaG up(RenG156-314)-R | CCAATTCTTCTTCAATATCTGGCCACATAAGTCATCACCTCCA CCTCAAA |
| (linker)DnaG-F | GGTGGATCTGGTGGTGGAGGATCAGTTGTTGATAAAGAGAA GATTAAAGAAATCAAAGAC |
| GyrA-LF | GGATGAACTTGAAGATATCAAACGCA |
| GyrA(linker)-R | CACCTCCACCAGAACCTCCACCTCCCTCATTTTCATTATTATT TAGTGAAGTGCCTTC |
| (kan)GyrA dn-F | ATTATATTTTACTGGATGAATTGTTTTAGAATGAAAATGAGT AGCTAGCAATGAAAAAAG |
| GyrA-RR | CAAGTTTGTAATCATGACGTTTCGG |

| | |
|---|---|
| IlvD-LF | CGCTGTTGTCAAAATGCTCG |
| IlvD(linker)-R | CACCTCCACCAGAACCTCCACCTCCTTTTTTGCCAGTTTCTTCAGGCTTC |
| (kan)IlvD dn-F | AATATTATATTTTACTGGATGAATTGTTTTAGAATCAAACATTAAAAAGACCGCTTGAA |
| IlvD-RR | TAAGCAATCAGTCCTGCGTTG |
| IlvD-LF-2 | CATCTATCTGTCCTGCTTCTGC |
| IlvD up(IFDC2)-R | TGCTATGAGTGTTATTGTTGCTCGGAGTCATTCAATGTAGAAATTTTACACACCAT |
| (IFDC2)IlvD up-F | TATTAGGTATACTACTGACAGCTTCATATGATGTGATGAAAGGAAGCGG |
| IlvD-RR-2 | CAGGTGTTAAATCATCAAAAGCCTTC |
| IlvD up(RenG156-314)-R | CCAATTCTTCTTCAATATCTGGCCACATTTGACCGCTTCCTTTCATCAC |
| (linker)IlvD-F | AGGTGGATCTGGTGGTGGAGGATCAACTGACAAAAAAACTCTTAAAGACTTAAGAAATC |
| MnmG-LF | GCTGACAAAGATCGTCATCAG |
| MnmG(linker)-R | CACCTCCACCAGAACCTCCACCTCCTCTTGTACGACTTCTATTCTTTCCTTCAAG |
| (kan)MnmG dn-F | AATATTATATTTTACTGGATGAATTGTTTTAGTAAAATGAAAAGAATTCCCCTTTGTTCC |
| MnmG-RR | TGTCCTACAACAAAGACACTATCTACTG |
| MnmG-LF-2 | GGTTTCTATCATGACGCACTTATGTC |
| MnmG up(IFDC2)-R | TGCTATGAGTGTTATTGTTGCTCGGTCGAAAATTGTGGGTCAAAAGACC |
| (IFDC2)MnmG up-F | TATTAGGTATACTACTGACAGCTTCGGGTTTTGGCTATTTTTGTGTTTGG |
| MnmG-RR-2 | TCATATTCAATGGCATAGCCAGTC |
| MnmG up(RenG156-314)-R | CCAATTCTTCTTCAATATCTGGCCACATTTCTATCTCCCAAACACAAAAATAGCC |
| (linker)MnmG-F | AGGTGGATCTGGTGGTGGAGGATCAATGACACACGAATTTACTGAAAACTATGA |
| PcrA-LF | GCATGAGTCTGTTAGAAGCTTCTG |
| PcrA(linker)-R | CACCTCCACCAGAACCTCCACCTCCTGTCTCCTTTTTCTCAAGGGGTG |
| (kan)PcrA dn-F | ATATTATATTTTACTGGATGAATTGTTTTAGGAAGAGAAAAGGTAAAGGACTTTGTTGC |
| PcrA-RR | AATGTCATGATAGGAAGGATCTGGT |
| PcrA-LF-2 | GACGAGAGTATCAATAAAAGTGCCT |
| PcrA up(IFDC2)-R | TGCTATGAGTGTTATTGTTGCTCGGTGTTTTTCATTCATTCCATGTAATAAAGGGT |

| | |
|---|---|
| (IFDC2)PcrA up-F | TATTAGGTATACTACTGACAGCTTCAGCGCAAGCTGTGAAAACA |
| PcrA-RR-2 | GTGTAGGGAATATTGGACTTAAGCAGAG |
| PcrA up(RenG156-314)-R | CCAATTCTTCTTCAATATCTGGCCACATAATCAAAAGCGGACCTTCCG |
| (linker)PcrA-F | AGGTGGATCTGGTGGTGGAGGATCAGCAGGTGCAGGGTCGGGTAA |
| PheT-LF | TGGTTTTGGTCTTTCTGGCAA |
| PheT(linker)-R | CACCTCCACCAGAACCTCCACCTCCTCTCACTTCAGCACCAACTTTTTCG |
| (kan)PheT dn-F | ATATTTTACTGGATGAATTGTTTTAGCGAAAAAATCTACCTTTAAAGGTAGATTTTTTTG |
| PheT-RR | ACAGTTTACCTTAGTTGTTTTAGGAACAC |
| SpaP-LF | GCTTGAATTGCGTCAGGATTTAG |
| SpaP(linker)-R | CACCTCCACCAGAACCTCCACCTCCATCTTTCTTAGCCTTTAAGCCAAGC |
| (Kan)SpaP dn-F | TTTACTGGATGAATTGTTTTAGCAGCATAGATATTACATTAGAATTAAAAAGTGAGATAG |
| SpaP-RR | CCACGCAGTTTTAATCCTCTTGT |
| SpaP-LF-2 | GCACAAGGAACCTATCACTTTACTAAAC |
| SpaP up(IFDC2)-R | TGAGTGTTATTGTTGCTCGGCTTCCAAATATATTTTTGTAATATACGTTAAAAAATTGT |
| (IFDC2)SpaP up-F | TATTAGGTATACTACTGACAGCTTCATTTATTCAGATTTGGAGGATTTATGTGGC |
| SpaP-RR-2 | TCTCATTGCGTTGCTTAATTGC |
| SpaP up(RenG156-314)-R | CCAATTCTTCTTCAATATCTGGCCACATAAATCCTCCAAATCTGAATAAATCTTCCA |
| (linker)SpaP-F | AGGTGGATCTGGTGGTGGAGGATCAAAAGTCAAAAAAACTTACGGTTTTCGT |
| TopA-LF | CGTCCATCTAATGTTAATCATACACCT |
| TopA(linker)-R | CACCTCCACCAGAACCTCCACCTCCTTTAACAGCTTTTTCCTTATAGTCACACTCC |
| (Kan)TopA dn-F | TATTATATTTTACTGGATGAATTGTTTTAGAGAAAGTGATGAAGTTGATGTATTGTACTT |
| TopA-RR | GGTATTCCAGACCAACTGGTTTCA |
| TopA-LF-2 | AGAAGATGGAAATAAATAGTATGACAAGCG |
| TopA up(IFDC2)-R | CTATGAGTGTTATTGTTGCTCGGTGATATATCATTTTTAGAGTACTGTCAAGAACAAAC |
| (IFDC2)TopA up-F | AGGTATACTACTGACAGCTTCTTCTAGAAGTTTATAACTTATAGGAAGTGTAGAATAATG |
| TopA-RR-2 | CATTGCGAACATGATTTCCGTG |

| | |
|---|---|
| TopA up(RenG156-314)-R | CTTCTTCAATATCTGGCCACATTATTCTACACTTCCTATAAGT TATAAACTTCTAGAATG |
| (linker)TopA-F | AGGTGGATCTGGTGGTGGAGGATCAACAAGTAAAACAACGA CAACAGTAAAAAAG |
| UvrA-LF | CCCAAGGCAAGTTGCTAAAAATAAAAAG |
| UvrA(linker)-R | CACCTCCACCAGAACCTCCACCTCCTTTTTCATTTAGTCTTCC CTTCAAATACTGTC |
| (kan)UvrA dn-F | AAATATTATATTTTACTGGATGAATTGTTTTAGATACAGGAC AGTATTTGAAGGGAAGAC |
| UvrA-RR | GTCAACACTTCACAACCATCCT |
| UvrA-LF-2 | AGGACATTTTCATAGATACCTGCC |
| UvrA up(IFDC2)-R | TGCTATGAGTGTTATTGTTGCTCGGGTGACGAAACAATACGT TTCTAAGCA |
| (IFDC2)UvrA up-F | TATTAGGTATACTACTGACAGCTTCAATTTTCCTATTTTCCTT TGGGGTGT |
| UvrA-RR-2 | CCTTCAAAAGGAATATCAATATCCCTGAC |
| UvrA up(RenG156-314)-R | CCAATTCTTCTTCAATATCTGGCCACATAATTAAGATACCAA GCCGTTAAACACC |
| (linker)UvrA-F | AGGTGGATCTGGTGGTGGAGGATCACAAGATAAATTAATTAT TCGTGGAGCGC |
| XseA-LF | CCATCATCATTGAAAAGGCAGAG |
| XseA(linker)-R | CACCTCCACCAGAACCTCCACCTCCAATGTTTTCTTCTTGTTT GACATTTTCTACC |
| (Kan) XseA dn-F | TAAAATATTATATTTTACTGGATGAATTGTTTTAGTGAAAGA TGGGCAGGTTCAAG |
| XseA-RR | TCAATTCCAGTAGCATTAAATTGTTCCTC |
| XseA-LF-2 | ACTGTCTCATTATTGATCTTGAGCG |
| XseA up(IFDC2)-R | TGCTATGAGTGTTATTGTTGCTCGGGAATCCTGTTCCAAAGTC TAAAACAAATGAC |
| (IFDC2)XseA up-F | TATTAGGTATACTACTGACAGCTTCAAGAATACCTTTTTCCTT TGTCTTTGGT |
| XseA-RR-2 | CAATCTTTGTCTCAGCAAATGTTCC |
| XseA up(RenG156-314)-R | CCAATTCTTCTTCAATATCTGGCCACATAAATCTCATCTAAAG ACCAAAGACAAAGG |
| (linker)XseA-F | AGGTGGATCTGGTGGTGGAGGATCATCAGATTATTTATCAGT ATCCTCTTTAACCAA |
| SrtA-LF | TGCAGATCACATAGCAGATAGTC |
| SrtA dn (IFDC2)-R | TGCTATGAGTGTTATTGTTGCTCGGTTAAAATGATATTTGATT ATAGGACTGCCTAAA |
| (IFDC2) SrtA dn-F | ATAAATTATTAGGTATACTACTGACAGCTTCAATAAGAAACT CGCTCTAACGAG |

| | |
|---|---|
| SrtA-RR | CATTTCGATACATATCCTGTGTCAC |
| SrtA (linker)-R | CTCCACCACCAGATCCACCTCCACCAAATGATATTTGATTATAGGACTGCCT |
| (RenG)SrtA dn-F | TGAAGAATGAGCAACGTTCTATCTAAAATAAGAAACTCGCTCTAACGAG |
| SrtA-LF-2 | GCTAAACCAGAACGTGTTATCAC |
| SrtA up(IFDC2)-R | TGCTATGAGTGTTATTGTTGCTCGGTGCTAGCTACTCATTTTCATTATTATTTAGT |
| (IFDC2)SrtA up-F | ATAAATTATTAGGTATACTACTGACAGCTTCAAAAAAGAACGTCAATCTAGGAAAAAAC |
| SrtA-RR-2 | GACTTCAAAACCCAATTGATTAACAT |
| SrtA up (RenG156-314)-R | ATTCTTCTTCAATATCTGGCCACATTGCTAGCTACTCATTTTCATTATTATTTAGT |
| (linker)SrtA-F | AGGTGGATCTGGTGGTGGAGGATCAAAAAAAGAACGTCAATCTAGGAAAAAAC |
| (erm)J1 dn-F | GTTATCTATTATTTAACGGGAGGAAATAAATCATAATTGTAAGCAGCTATTATTAAAGAG |
| (erm)J2 dn-F | AGTTATCTATTATTTAACGGGAGGAAATAAAAGAGGCTTGATTGATACTGTTT |
| (linker)RenG-F | AGGTGGATCTGGTGGTGGAGGATCAGCTAGTAAAGTTTATGATCCTGAACAAC |
| (kan)ThiI dn-F | CGAGGAATTTGTATCGACTCTAGAAACAAGAAAAATCATATATTAATCATTTTCTTCATG |
| RenG(kan)-R | CAAGCTATAAGGTTATTGTCCTGGGTTAGATAGAACGTTGCTCATTCTTCAAA |
| Kan-F | CCCAGGACAATAACCTTATAGCTTGTA |
| Kan-R | CTAGAGTCGATACAAATTCCTCGTAGG |
| (kan)GltB dn-F | CTACGAGGAATTTGTATCGACTCTAGGAGTAGAACAAAAGAGACAGGACT |
| AlaS(linker)-R-2 | CTCCACCACCAGATCCACCTCCACCCAAATTCTCTGCTACTGCGGC |
| DnaG up (RenG)-R | GTTCAGGATCATAAACTTTACTAGCCATAAGTCATCACCTCCACCTCAAA |
| (DnaG up)RenG-F | TTTGAGGTGGAGGTGATGACTTATGGCTAGTAAAGTTTATGATCCTGAACAACG |
| GyrA(linker)-R-2 | CTCCACCACCAGATCCACCTCCACCCTCATTTTCATTATTATTTAGTGAAGTGCCTTC |
| IlvD(linker)-R-2 | CTCCACCACCAGATCCACCTCCACCTTTTTTGCCAGTTTCTTCAGGCTTC |
| MnmG(linker)-R-2 | CTCCACCACCAGATCCACCTCCACCTCTTGTACGACTTCTATTCTTTCCTTCAAG |

| | |
|---|---|
| PcrA(liner)-R-2 | CTCCACCACCAGATCCACCTCCACCTGTCTCCTTTTTCTCAAGGGGTG |
| PheT(linker)-R-2 | CTCCACCACCAGATCCACCTCCACCTCTCACTTCAGCACCAACTTTTTCG |
| SpaP(linker)-R-2 | CTCCACCACCAGATCCACCTCCACCATCTTTCTTAGCCTTTAAGCCAAGC |
| TopA(linker)-R-2 | CTCCACCACCAGATCCACCTCCACCTTTAACAGCTTTTTCCTTATAGTCACACTCC |
| UvrA(linker)-R-2 | CTCCACCACCAGATCCACCTCCACCTTTTTCATTTAGTCTTCCCTTCAAATACTGTC |
| XseA(linker)-R-2 | CTCCACCACCAGATCCACCTCCACCAATGTTTTCTTCTTGTTTGACATTTTCTACC |
| LDH-F | ATGACTGCAACTAAACAACATAAAAAAGT |
| LDH(rbs-Barstar)-R | CCTTCTTCATTCTCTAAACATCTCCTTCTTAGTTACGAGCTGCAGCAGC |
| (erm)LDH dn-F | ATCTATTATTTAACGGGAGGAAATAAACAATAAAAATCCATAAAAATACCAAGCATTGTG |
| LDH-RR | CGTTTTCGCACTTATTTAATAGCATC |
| (rbs)Barstar-F | GAAGGAGATGTTTAGAGAATGAAGAAGGC |
| RpoB-LF | GGTGACAAATCACGTGAAGTTC |
| RpoB(rbs-Barnase)-R | CTTGAGCCATGTTTTAGCTCCTTCTTCTTTATTTCTCATCACTATTTGCAAATTCAGCA |
| (Kan)RpoB dn-F | TTTACTGGATGAATTGTTTTAGGTGATTTAGTAACTTAGTTTATCTTTTATGATTGGTAC |
| RpoB-RR | CCAAACGATTGTTCCGATTGATC |
| (rbs)Barnase-F | AGAAGAAGGAGCTAAAACATGGCT |
| MecA(linker)-R-2 | TAGTCAGAACCACCAGAACCACCAGAACCACCTCCAATCATTTGTAATTCTTGCAGAGC |
| (erm)MecA dn-F-2 | TATCTATTATTTAACGGGAGGAAATAAGCTAGATGATACCAATTACCTTAAAATTTGTTC |
| C-Flag3-F | GGTGGTTCTGGTGGTTCTGGTG |
| C-Flag3-R | CTTATCGTCATCGTCCTTGTAATCG |
| (FLAG3)erm-F | CGATTACAAGGACGATGACGATAAGTAAGAAGGAGTGATTACATGAACAAAAATATAAAA |
| ClpC up(kan)-R | TTTATTATTTCCTTCCTCTTTTCTACATCATAGATGGTATCCTTTTCTATCTAAACGTTG |
| (ClpC up)kan-F | TTTAGATAGAAAAGGATACCATCTATGATGTAGAAAAGAGGAAGGAAATAATAAATGGC |
| kan(rbs)-R | GCTTCCAACTCCTTCCCTAAAACAATTCATCCAGTAAAATATAATATTTTATTTTCTCC |

| | |
|---|---|
| (kan)HA3-F | GGATGAATTGTTTTAGGGAAGGAGTTGGAAGCATGTACCCAT ACGATGTTCCTGACTAT |
| HA3(linker)-R | TGATCCTCCACCACCAGATCCACCTCCACCAGCGTAATCTGG AACGTCATATGG |
| (linker)C-HA3-F | GGAGGTGGAGGTTCTGGTGGAGGTGGATCTTACCCATACGAT GTTCCTGACTAT |
| C-HA3-R | AGCGTAATCTGGAACGTCATATGG |
| (HA3)kan-F-2 | TCCATATGACGTTCCAGATTACGCTTAATGTAGAAAAGAGGA AGGAAATAATAAATGGC |
| Thil(linker)-R-2 | AGAACCACCAGAACCACCAGAACCACCCAATAAATCTTCGA TTAAGACATCAACTTCATC |
| (kan)Thil dn-F | CGAGGAATTTGTATCGACTCTAGAAACAAGAAAAATCATAT ATTAATCATTTTCTTCATG |
| C-HA3-F | GGTGGTTCTGGTGGTTCTGGTGGTTCTTACCCATACGATGTTC CTGACTATGC |
| (HA3)kan-F | TCCATATGACGTTCCAGATTACGCTTAACCCAGGACAATAAC CTTATAGCTTGTA |
| GltB(linker)-R-2 | AACCACCAGAACCACCAGAACCACCCTCTGTAACCATCATTC TGAGGTTTTG |
| DnaG up(HA3)-R | CATAGTCAGGAACATCGTATGGGTACATAAGTCATCACCTCC ACCTCA |
| N-HA3-F | ATGTACCCATACGATGTTCCTGACTATGC |
| N-HA3(linker)-R | TGATCCTCCACCACCAGATCCACCTCCACCAGCGTAATCTGG AACGTCATATGG |
| SrtA(linker)-R-2 | AGAACCACCAGAACCACCAGAACCACCAAATGATATTTGAT TATAGGACTGCCTAAAAGC |
| (kan)SrtA dn-F | CCTACGAGGAATTTGTATCGACTCTAGAATAAGAAACTCGCT CTAACGAGTTTC |
| MecA up(erm)-R | TTTTATATTTTTGTTCATGTAATCACTCCTTCCGCTGATTTGTT TCATTTCCATAGTCT |
| (erm)MecA dn-F | AGTTATCTATTATTTAACGGGAGGAAATAACTGCAAGAATTA CAAATGATTGGATAAGCT |
| ClpP-LF | CCGCTAGAAGAGGTTGAGATTG |
| Clp(Tet)-R | TCTTCATGTGATTTTCCTCCATTCAATCATAAGAGCGTTCACC ACG |
| (Tet) ClpP dn-F | TGTTCAATAAAATAACTTAGTGCGGGCTTCATCGATGAAATC ATGG |
| ClpP-RR | AAGCTGCAAGAACAGCTGCACTGA |
| Tet-F | TGAATGGAGGAAAATCACATGAAGA |
| Tet-R | CGCACTAAGTTATTTTATTGAACA |

**Table S3. Normalized luciferase values of strains employed in the study**

| Strains | Luciferase signal (LUC/OD, ×$10^3$) | Strains | Luciferase signal (LUC/OD, ×$10^3$) |
|---|---|---|---|
| UA159(BG) | 3.19 ± 0.41 | MecA-NTD + CTD-MnmG | 35.36 ± 5.24 |
| NTD1 + CTD1 | 3.67 ± 0.30 | MecA-NTD + PcrA-CTD | 1102.57 ± 29.86 |
| NTD2 + CTD2 | 3.59 ± 0.29 | MecA-NTD + CTD-PcrA | 3.70 ± 0.20 |
| J1-NTD1 + J2-CTD1 | 41050.29 ± 5574.41 | MecA-NTD + PheT-CTD | 3111.67 ± 72.25 |
| J1-NTD2 + J2-CTD2 | 31417.60 ± 212.07 | MecA-NTD + CTD-PheT | ND* |
| MecA-NTD + ClpC-CTD | 3852.81 ± 58.67 | MecA-NTD + SpaP-CTD | 298.81 ± 10.12 |
| MecA-NTD + CTD-ClpC | 4599.76 ± 359.45 | MecA-NTD + CTD-SpaP | 202.65 ± 24.65 |
| NTD-MecA + ClpC-CTD | 363.04 ± 9.44 | MecA-NTD + TopA-CTD | 1498.29 ± 28.26 |
| NTD-MecA + CTD-ClpC | 284.21 ± 8.59 | MecA-NTD + CTD-TopA | 548.03 ± 22.87 |
| MecA-CTD + ClpC-NTD | 2701.43 ± 150.76 | MecA-NTD + UvrA-CTD | 1361.54 ± 11.53 |
| MecA-CTD + NTD-ClpC | 3148.89 ± 172.23 | MecA-NTD + CTD-UvrA | 89.39 ± 2.75 |
| CTD-MecA + ClpC-NTD | 342.78 ± 9.32 | MecA-NTD + XseA-CTD | 471.75 ± 36.11 |
| CTD-MecA + NTD-ClpC | 264.56 ± 9.01 | MecA-NTD + CTD-XseA | 17.35 ± 0.67 |
| MecA-NTD + ComX-CTD | 412.26 ± 14.25 | MecA-NTD + SrtA-CTD | 3.34 ± 0.12 |
| MecA-NTD + CTD-ComX | 177.70 ± 20.09 | MecA-NTD + CTD-SrtA | 2.78 ± 0.12 |
| NTD-MecA +ComX-CTD | 22.55 ± 2.11 | NTD-MecA + SrtA-CTD | 2.77 ± 0.28 |
| NTD-MecA + CTD-ComX | 14.14 ± 0.31 | NTD-MecA + CTD-SrtA | 3.65 ± 0.74 |
| MecA-CTD + ComX-NTD | 260.23 ± 56.93 | J1-Luc | 42566.67 ± 2758.02 |
| MecA-CTD + NTD-ComX | 11.86 ± 0.54 | J2-Luc | 41987.67 ± 1289.06 |
| CTD-MecA + ComX-NTD | 34.40 ± 4.10 | MecA-Luc | 17333.33 ± 725.72 |
| CTD-MecA + NTD-ComX | 11.50 ± 0.44 | Luc-ClpC | 8330.01 ± 259.62 |
| MecA-NTD + ThiI-CTD | 944.81 ± 31.20 | ComX-Luc | 6546.67 ± 1036.64 |
| MecA-NTD + CTD-ThiI | 19.48 ± 0.55 | ThiI-Luc | 10596.67 ± 1353.15 |
| NTD-MecA + ThiI-CTD | 50.44 ± 2.46 | GltB-Luc | 16366.67 ± 601.39 |
| NTD-MecA + CTD-ThiI | 2.81 ± 0.36 | AlaS-Luc | 123596.40 ± 3291.47 |
| MecA-NTD + GltB-CTD | 1219.41 ± 23.02 | Luc-DnaG | 23116.23 ± 2137.88 |
| MecA-NTD + CTD-GltB | 619.89 ± 13.08 | GyrA-Luc | 52331.23 ± 652.84 |
| NTD-MecA + GltB-CTD | 144.99 ± 5.42 | IlvD-Luc | 136369.01 ± 5541.43 |
| NTD-MecA + CTD-GltB | 63.78 ± 1.57 | MnmG-Luc | 46741.51 ± 3846.13 |

| | | | |
|---|---|---|---|
| MecA-NTD + AlaS-CTD | 953.33 ± 33.27 | PcrA-Luc | 20718.61 ± 778.58 |
| MecA-NTD + CTD-AlaS | 3.59 ± 0.27 | PheT-Luc | 96772.74 ± 1348.15 |
| MecA-NTD + DnaG-CTD | 458.21 ± 14.18 | SpaP-Luc | 215359.53 ± 13684.56 |
| MecA-NTD + CTD-DnaG | 1207.89 ± 34.04 | TopA-Luc | 29296.11 ± 738.19 |
| MecA-NTD + GyrA-CTD | 2230.36 ± 118.32 | UvrA-Luc | 25205.97 ± 578.74 |
| MecA-NTD + CTD-GyrA | ND* | XseA-Luc | 14920.21 ± 684.53 |
| MecA-NTD + IlvD-CTD | 2272.11 ± 70.63 | Barstar-NTD + Barnase-CTD | 2594.30 ± 54.47 |
| MecA-NTD + CTD-IlvD | 871.87 ± 74.8 | RpoB-Barnase-Luc | 4886.77 ± 16.14 |
| MecA-NTD + MnmG-CTD | 1386.12 ± 24.78 | LDH-Barstar-Luc | 115410.6 ± 16917.08 |

***Not determined; presumably these constructs are lethal and cannot be generated**

### Supplemental Figures

**Figure S1.**

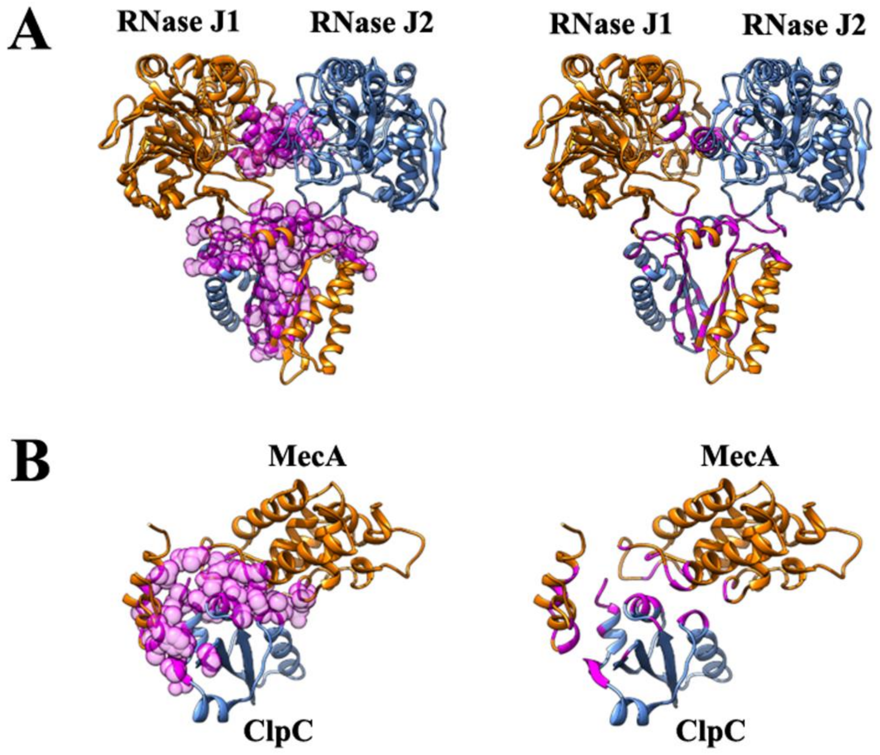


**Figure S1. Models of J1-J2 and MecA-ClpC highlighting the molecular interfaces.** (A) Model of the RNase J1-J2 heterodimer interface (2571 Å$^2$) with the interacting residues shown in magenta spheres (left) or only colored at the ribbon level (right). (B) A truncated model of the *S. mutans* MecA-ClpC complex (988 Å$^2$) showing the MecA (blue) and ClpC (orange) interface. Interacting residues are shown in spheres on the left and as a colored ribbon on the right.

**Figure S2.**

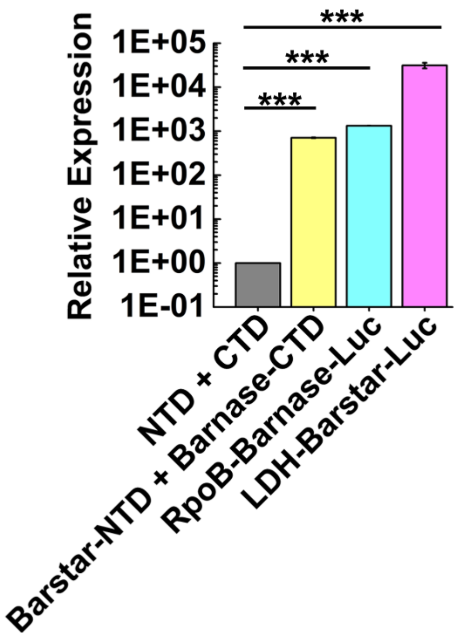


**Figure S2. SLCA and whole luciferase measurements of the barnase-barstar complex.** The barnase-barstar complex was used as a positive control due to its well-characterized high affinity interaction. Barstar luciferase fusions were expressed using the lactate dehydrogenase promoter P*ldh*, whereas barnase luciferase fusions were expressed using the promoter for the Beta subunit of RNA polymerase P*rpoB*. Cultures were grown to mid-logarithmic growth phase, normalized to the measured optical density ($OD_{600}$) values, and then expressed relative to the luciferase values of the unfused NTD and CTD split luciferase fragments, which were arbitrarily assigned values of 1. All luciferase data are expressed as the means ± s.d. (indicated by error bars) derived from three or four biological replicates. ***$p < 0.001$ Unpaired two-tailed Student's *t*-test with Welch's correction.
